# Temporal transcriptomic remodeling after controlled cortical impact reveals delayed AQP4/SNTA1 expression imbalance associated with ion-homeostatic remodeling

**DOI:** 10.64898/2026.09.10.750706

**Authors:** Benu George, Eric B. Emmons, Qiang Zhang

## Abstract

Traumatic brain injury (TBI) is often viewed as a progressively evolving molecular response, yet the molecular and cellular responses that dictate acute TBI may differ fundamentally from those that define the chronically remodeled brain. We integrated temporally-resolved transcriptomic analyses with independent mouse, human, proteomic, and spatial datasets to determine how neural-circuit and astrocyte-homeostatic transcriptional responses reorganize after controlled cortical impact (CCI). In GSE269748, a mouse CCI transcriptomic dataset spanning acute to chronic post-injury timepoints, transcriptomic remodeling was non-monotonic. The number of genes showing large expression changes was greatest at 7 days, with extensive transcriptional changes persisting at 6 months. A more selective astrocytic phenotype emerged at 7 days and persisted at 6 months, with *Aqp4* increasing disproportionately relative to *Snta1*, producing a sustained AQP4/SNTA1 expression imbalance. This imbalance occurred alongside heterogeneous endfoot remodeling and could not be explained by inflammatory stress alone. Among 14 prespecified biological processes, potassium and ion homeostasis showed the most specific and robust association with the AQP4/SNTA1 imbalance after adjustment for broader reactivity and AP-1 activity; this observation remained stable to gene, sample, and injury-model perturbation. Circuit-associated analyses provided complementary context, showing that dopamine recipient changes occurred within broader neurotransmitter system remodeling rather than as a unique signal. Integrative temporal analysis distinguished acute stress-dominant remodeling from delayed astrocyte-homeostatic changes, with AQP4/SNTA1 imbalance and potassium and ion homeostatic remodeling becoming most prominent at later stages. External datasets provided limited contextual support while defining clear limits to generalization. Together, these findings identify delayed AQP4/SNTA1 expression imbalance within potassium and ion homeostatic remodeling as a testable feature of chronic post-traumatic astrocyte biology.

## Introduction

Traumatic brain injury (TBI) is increasingly recognized as a heterogeneous and evolving neurological condition rather than a single acute event [1-3]. Patients with apparently similar injuries can follow markedly different cognitive and functional trajectories, and chronic consequences may persist for months or years. Major TBI frameworks therefore emphasize biologically informed characterization beyond conventional severity categories [1-3]. Yet, the molecular counterpart of this heterogeneity remains incompletely resolved. In particular, it is unclear whether acute and chronic post-traumatic tissue primarily differ in the magnitude of a common injury response or instead reflect changing contributions of distinct molecular and cellular responses over time.

This distinction has translational relevance because pharmacological responses after TBI vary with clinical phenotype and recovery interval [4-8]. Amantadine can improve recovery during a defined post-injury window [4,5], whereas methylphenidate is used for selected cognitive deficits [6,7], and molecular imaging demonstrates dopaminergic abnormalities in only a subset of cognitively impaired patients after TBI [8]. Together, these observations raise the question of whether dopamine-associated abnormalities represent isolated circuit phenotypes or components of broader neurotransmitter remodeling. These observations do not establish that molecular profiling can guide treatment, but they illustrate a broader problem: clinically defined TBI populations can contain biologically different forms of circuit dysfunction while treatment remains largely phenotype- and time-based [3,10].

Transcriptomic studies increasingly support a multidimensional view of injury biology. Human single-nucleus profiling and a large murine single-cell TBI atlas have revealed marked cell-, model-, region-, and time-dependent responses [9,10]. Astrocytes are particularly relevant because they regulate glutamate uptake, extracellular potassium regulation, metabolic exchange, water transport, and the neurovascular interface. Classic controlled cortical impact (CCI) studies have demonstrated post-traumatic reductions in GLT-1 and GLAST [11], while recent longitudinal and perturbational work shows that astrocyte responses remain dynamic into chronic injury and that inflammatory signaling can be dissociated from homeostatic function [12,13]. These observations argue against treating reactive astrogliosis as a fixed molecular state.

The astrocytic perivascular endfoot provides a biologically compelling point of convergence. AQP4 (aquaporin-4) is enriched at astrocytic endfeet, where its polarized localization depends in part on alpha-syntrophin, encoded by SNTA1, and the dystrophin-associated complex [14-16]. Alpha-syntrophin loss disrupts perivascular AQP4 localization and delays extracellular K+ clearance after neuronal activation [17], directly linking endfoot organization to ion homeostasis. TBI can also increase total AQP4 while reducing polarized perivascular localization [18], and experimental injury has been associated with altered glymphatic transport [19,20]. These studies make the AQP4-SNTA1 axis mechanistically relevant, but they also establish an important boundary: transcript abundance does not measure AQP4 polarization, physical protein coupling, potassium clearance, or glymphatic function.

Here, we first asked whether the acute-to-chronic CCI time course is characterized by temporally distinct transcriptional responses and whether a delayed AQP4/SNTA1 expression imbalance emerges within astrocyte-associated remodeling. We then tested whether this imbalance is preferentially associated with ion homeostasis rather than generalized inflammatory reactivity. As a complementary analysis, we examined circuit-associated transcriptional changes to determine whether dopamine-related alterations represent a selective post-traumatic signal or occur within broader multi-neurotransmitter remodeling processes. We addressed these questions using temporally resolved analysis of the GSE269748 single-cell RNA-sequencing dataset together with complementary mouse, human, proteomic, perturbational, neuropathological, and spatial datasets to assess the robustness, biological context, and generalizability of the findings.

## Results

### CCI induces non-monotonic, temporally structured transcriptomic remodeling

GSE269748 contains 36 GEO (gene expression omnibus) samples across several experimental strata; the primary temporal analysis used the 21 metadata-defined male focal CCI/Naive samples (Naive: n = 7, 24 h: n = 6, 7 d: n = 4, 6 mo: n = 4; Figure 1A; Supplementary Figure 1A,B; Supplementary Table S01). Supplementary Figure 1A summarizes the overall evidence-building strategy, whereas Supplementary Figure 1B places GSE269748 within the broader cross-dataset framework used for contextual testing and evidence-boundary analyses. Whole-transcriptome PCA showed marked stage-associated organization, with PC1 and PC2 explaining 42.8% and 23.2% of variance, respectively (Figure 1B; Supplementary Table S04). Hierarchical sample-correlation analysis provided a concordant representation of this structure (Supplementary Figure 1C; Supplementary Table S07).

**Figure 1.**
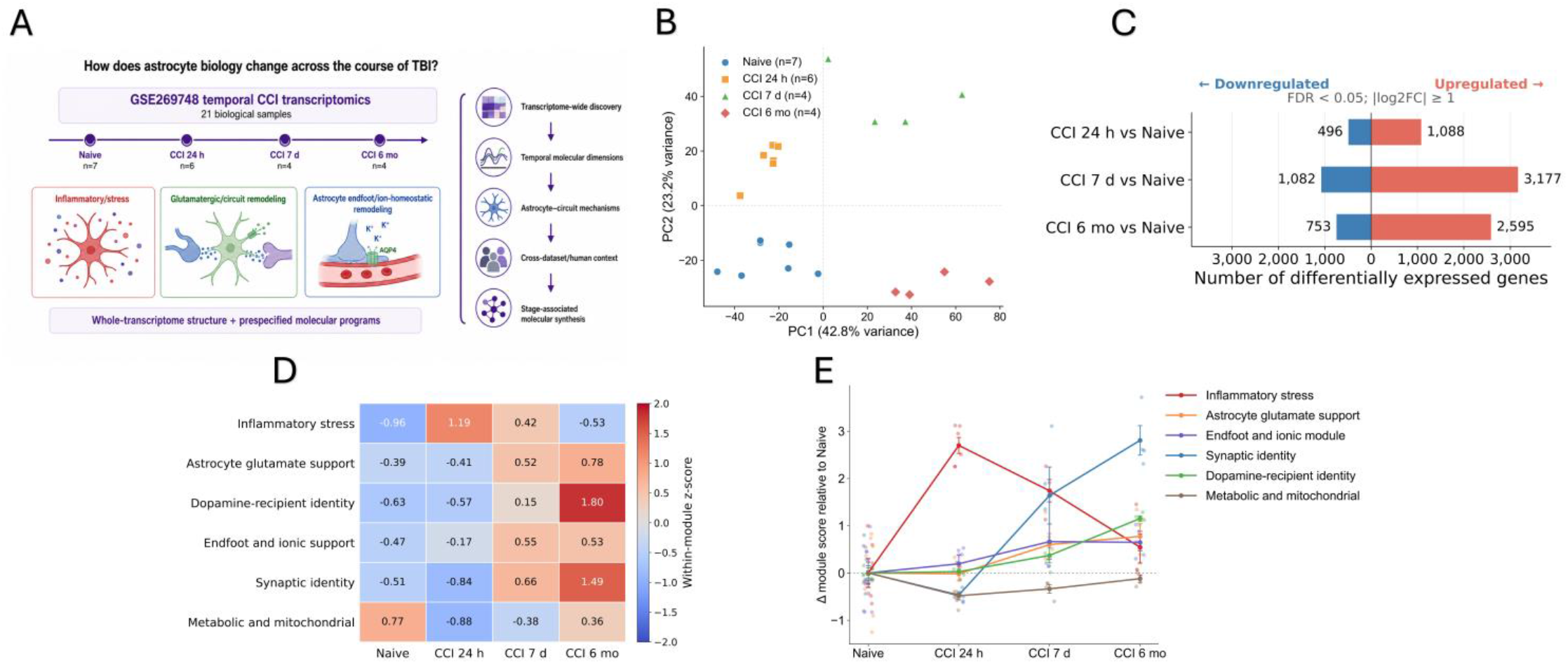
Temporal transcriptomic landscape and biological-response organization after CCI. (A) Overview of the study design and analytical framework. The primary temporal analysis used the 21 metadata-defined male focal CCI/Naive samples from GSE269748 and animal-level astrocyte-like pseudobulk profiles: Naive (n = 7), CCI 24 h (n = 6), CCI 7 d (n = 4), and CCI 6 mo (n = 4). The study was structured to determine whether post-CCI transcriptional remodeling followed a uniform progressive response or resolved into temporally distinct molecular dimensions. (B) Whole-transcriptome principal component analysis (PCA) of the 21 astrocyte-like pseudobulk samples. PC1 and PC2 explained 42.8% and 23.2% of total variance, respectively, and revealed marked stage-associated organization of Naive, 24-h, 7-d, and 6-mo samples. (C) Transcriptome-wide differential-expression burden for each CCI stage relative to Naive. Differentially expressed genes were defined using FDR < 0.05 and |log2FC| ≥ 1 within a common 16,209-gene testing universe. At 24 h, 1,088 genes were upregulated and 496 were downregulated; at 7 d, 3,177 were upregulated and 1,082 were downregulated; and at 6 mo, 2,595 were upregulated and 753 were downregulated. The largest large-effect differential-expression burden occurred at 7 d, with extensive remodeling persisting at 6 mo. (D) Temporal landscape of six predefined neural and astrocyte-associated transcriptional responses: inflammatory stress, astrocyte glutamate support, dopamine-recipient identity, endfoot and ionic module, synaptic identity, and metabolic and mitochondrial function. Values represent standardized scores across the four injury stages, illustrating distinct rather than uniformly progressive temporal patterns. (E) Sample-level trajectories of the same six transcriptional responses expressed relative to the Naive reference. Individual sample values are shown together with stage-level summaries, demonstrating heterogeneous temporal responses. Inflammatory-stress remodeling was most prominent acutely, whereas astrocyte glutamate support, endfoot and ionic module, synaptic, and dopamine-recipient responses became more prominent at later stages. Connecting lines summarize cross-sectional stage patterns and do not represent longitudinal measurements from the same animals.

Differential-expression gene burden (DEG-burden) was non-monotonic (Figure 1C). At FDR < 0.05 and |log2FC| ≥ 1, 1,584 genes were altered at 24 h, 4,259 at 7 d, and 3,348 at 6 mo relative to Naive (Supplementary Table S05). The ordering 7 d > 6 mo > 24 h was preserved both without a fold-change threshold and at |log2FC| ≥ 0.5, indicating that the non-monotonic DEG-burden pattern was not dependent on a single effect-size cutoff (Supplementary Figure 1D; Supplementary Table S05). Chronic gene-level effects were also robust to stricter astrocyte-like selection: across 10,326 commonly tested genes, primary- and strict-gate effects correlated at ρ = 0.782, and 520 of 523 genes significant in both analyses showed concordant directions (99.4%; Supplementary Figure 1E; Supplementary Table S06). This sensitivity analysis supports robustness of the chronic Naive-versus-6-mo transcriptional signal to astrocyte-selection stringency but does not constitute replication of the complete time course.

Six predefined transcriptional responses further revealed heterogeneous temporal organization (Figure 1D,E; Supplementary Table S08). Inflammatory stress-related activity was most prominent acutely, whereas astrocyte glutamate support, endfoot and ionic support, synaptic identity, and dopamine-recipient identity became more prominent at later stages; the metabolic and mitochondrial response followed a distinct pattern. These divergent trajectories established a temporally heterogeneous molecular framework for the focused circuit and astrocyte analyses that followed.

### Circuit-associated remodeling provides context across multiple neurotransmitter systems

To determine whether dopamine-associated changes represented a selective signal or one component of a broader neural response, we examined circuit-associated transcriptional remodeling using the matched all-QC pseudobulk. Transcriptome-wide enrichment showed acute negative enrichment of several neurotransmissionrelated pathways, followed by broad positive neuronal and synaptic enrichment at 7 d and 6 mo (Figure 2A; Supplementary Table S11). Across seven transmitter systems, dopaminergic enrichment was not uniquely dominant (Figure 2B). By 6 mo, GABAergic, glutamatergic, dopaminergic, serotonergic, adrenergic, and cholinergic gene sets were all positively enriched, whereas the purinergic/adenosine gene set showed a distinct negative pattern (Supplementary Table S12). Removing genes shared across transmitter systems reproduced this stage structure in non-overlapping modules, and sample-level trajectories showed the same delayed increase across most transmitter-associated modules (Figure 2C; Supplementary Figure 2E; Supplementary Tables S13-S15), arguing against pathway overlap as the sole explanation.

**Figure 2.**
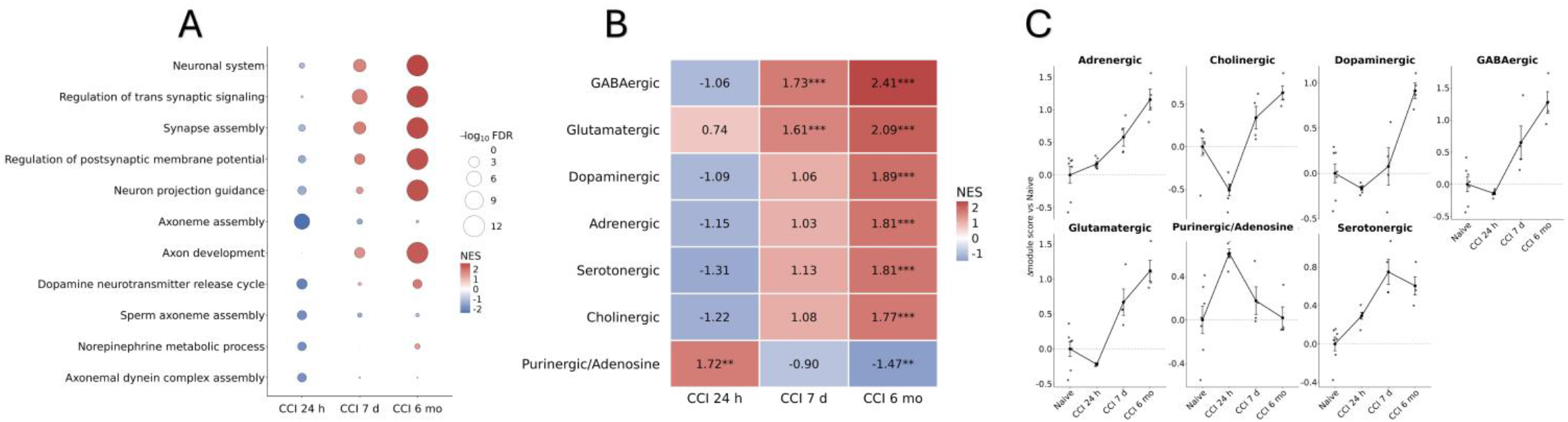
Stage-structured circuit and multi-neurotransmitter transcriptional remodeling after CCI. (A) Transcriptome-wide enrichment of neural, synaptic, and neurotransmitter-associated pathways across CCI 24 h, 7 d, and 6 mo relative to Naive. Circle color indicates normalized enrichment score (NES), and circle size indicates −log10 FDR. Acute CCI showed negative enrichment of several neurotransmission-associated processes, followed by broader positive neuronal and synaptic enrichment at 7 d and 6 mo (Supplementary Table S11). (B) Gene set enrichment across seven neurotransmitter-associated systems. Heatmap values indicate NES relative to Naive. At 6 mo, positive enrichment was observed across GABAergic, glutamatergic, dopaminergic, serotonergic, adrenergic, and cholinergic gene sets, whereas the purinergic/adenosine gene set followed a distinct negative pattern. Dopaminergic enrichment therefore occurred within a broader multi-neurotransmitter response rather than as a uniquely dominant signal (Supplementary Table S12). (C) Individual-sample trajectories for seven non-overlapping neurotransmitter-associated modules derived after removing genes shared across transmitter systems. Points represent individual biological samples; larger points and connecting lines indicate group means with SEM. Most transmitter-associated modules increased at delayed stages, whereas the purinergic/adenosine module followed a distinct temporal pattern (Supplementary Tables S13-S15). All analyses used matched GSE269748 all-QC animal-level pseudobulk profiles (Naive, n = 7; CCI 24 h, n = 6; CCI 7 d, n = 4; CCI 6 mo, n = 4). These transcriptional measures do not directly assess neurotransmitter release, receptor activation, synaptic transmission, or circuit function. Additional gene-level, leave-one-gene-out, and cellular-composition sensitivity analyses are shown in Supplementary Figure 2 and Supplementary Tables S16-S18.

Additional targeted analyses supported the same interpretation. Prespecified synaptic-identity and dopaminerecipient module trajectories showed delayed increases (Supplementary Figure 2A,B; Supplementary Table S17), while targeted gene-level analysis showed predominantly increased expression at 7 d and 6 mo with heterogeneity among individual genes (Supplementary Figure 2F; Supplementary Table S16). The chronic dopamine-recipient module remained positive after sequential omission of every constituent gene (Supplementary Figure 2C; Supplementary Table S18). Because all-QC pseudobulk can reflect cell composition, outcome-independent samplelevel composition PCs were derived from 278,108 QC-passing cells. Adjustment for composition PC1 retained the overall stage association for synaptic identity and dopamine-recipient identity and preserved positive 6-mo effects, whereas several earlier coefficients were attenuated (Supplementary Figure 2D; Supplementary Table S18). Together, these analyses identify broad multi-neurotransmitter transcriptional remodeling rather than a selective dopaminergic response.

### Delayed CCI is associated with disproportionate *Aqp4* relative to *Snta1* expression

We next focused on *Aqp4* and *Snta1* as an astrocytic endfoot-associated axis. In the astrocyte-like pseudobulk, *Aqp4* increased at delayed stages and was elevated at 6 mo (+1.85 log2[CPM+1] versus Naive, FDR = 0.0182), whereas *Snta1* was reduced at 24 h, 7 d, and 6 mo (all FDR ≤ 0.0164; Figure 3A; Supplementary Tables S20-S22). Importantly, neither transcript was used in the positive astrocyte gate.

**Figure 3.**
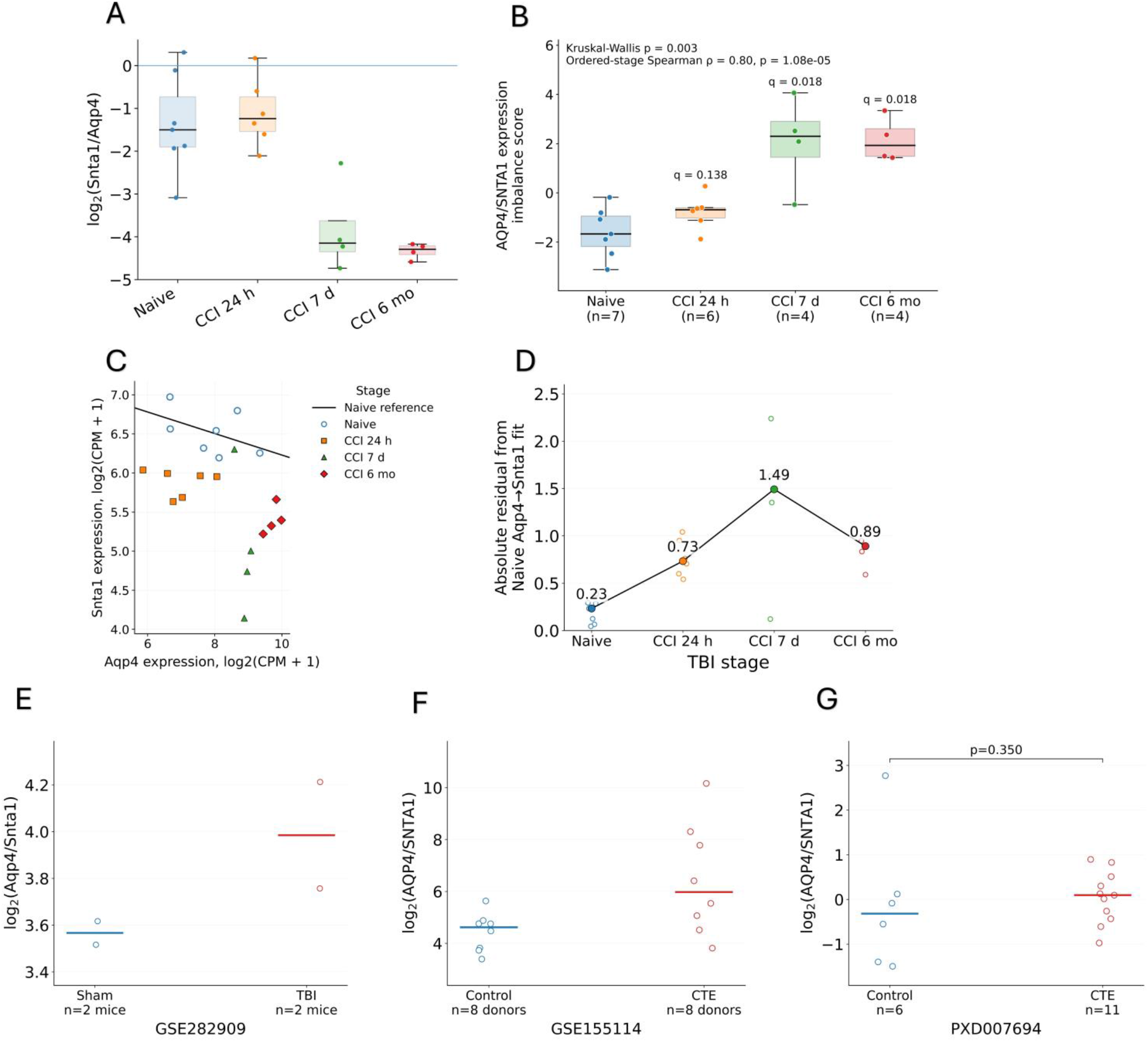
Delayed AQP4/SNTA1 expression imbalance and external evidence boundaries. (A) Temporal relationship between *Aqp4* and *Snta1* expression in GSE269748 astrocyte-like pseudobulk samples. Individual biological samples and group summaries are shown for Naive (n = 7), CCI 24 h (n = 6), CCI 7 d (n = 4), and CCI 6 mo (n = 4). Delayed stages were characterized by increasing *Aqp4* relative to *Snta1* expression (Supplementary Tables S20-S22). (B) Naive-referenced AQP4/SNTA1 expression-imbalance score across the CCI time course. Higher values indicate greater *Aqp4* expression relative to *Snta1*. The imbalance was not increased at 24 h but was significantly elevated at 7 d and 6 mo. Exact two-sided permutation tests were used for comparisons with Naive, with Benjamini-Hochberg correction as specified in the analysis plan (Supplementary Tables S21-S23). (C) Naive-reference *Aqp4*-*Snta1* expression relationship. The regression line was estimated from Naive samples and subsequently used as a reference for evaluating post-injury deviation. Symbols identify injury stages. This analysis evaluates an expression relationship and does not imply physical molecular coupling. (D) Absolute deviation from the Naive *Aqp4*-*Snta1* reference relationship. Points show individual biological samples and connected group summaries show median absolute residuals. Departure was greatest at 7 d and remained elevated at 6 mo (Supplementary Table S24). (E) Mouse-level spatial transcriptomic context from GSE282909. Values represent astrocyte-enriched log2(*Aqp4*/*Snta1*) summaries for Sham and TBI mice. Sections from the same mouse are connected, and horizontal bars denote mouse-level group means. With n = 2 mice per condition, these data provide trend-level spatial context only (Supplementary Table S26). (F) Chronic human transcriptomic context from GSE155114. Donor-level log2(AQP4/SNTA1) summaries are shown for Control and CTE donors (n = 8 per group). The observed direction is presented as contextual human support rather than independent mechanistic validation (Supplementary Table S27). (G) Human CTE proteomic context from PXD007694. Sample-level log2(AQP4/SNTA1) protein contrasts are shown for Control (n = 6) and CTE (n = 11) groups. The group difference was not statistically significant (twosided Mann-Whitney p = 0.350), and the analysis is therefore interpreted as directional cross-modal context rather than proteomic validation (Supplementary Table S27).

The Naive-referenced AQP4/SNTA1 imbalance was unchanged at 24 h but increased at 7 d (+2.46, FDR = 0.0164) and 6 mo (+2.97, FDR = 0.0091; Figure 3B; Supplementary Tables S21-S23). Alternative difference and zscore formulations reproduced the delayed separation (Supplementary Figure 3A-C). Within Naive samples, *Aqp4* and *Snta1* were essentially uncorrelated, but post-injury samples increasingly departed from the Naive reference relationship, with the largest median absolute residual at 7 d (Figure 3C,D; Supplementary Table S24).

External mouse and human datasets provided limited and context-dependent support for the delayed AQP4/SNTA1 expression imbalance. The prespecified 7-d comparison in GSE180862 an independent single-cell mouse mTBI dataset sampling hippocampus and frontal cortex [21], did not reproduce the discovery direction (Supplementary Figure 3D), whereas GSE282909, a Visium spatial-transcriptomic dataset from a mild, diffuse CHIMERA TBI model [26], provided only trend-level spatial context because n = 2 mice per condition (Figure 3E). Human chronic traumatic encephalopathy (CTE) transcriptomic (GSE155114) and proteomic (PXD007694) datasets showed directionally higher AQP4/SNTA1 contrasts, but the proteomic difference was not significant (p = 0.350; Figure 3F,G). We therefore interpret the delayed imbalance as a robust transcriptional feature of the discovery cohort with limited external support from spatial mouse data and directional human transcriptomic and proteomic findings, rather than as a universal TBI signature or evidence of physical AQP4-SNTA1 uncoupling.

### AQP4/SNTA1 expression imbalance accompanies heterogeneous endfoot remodeling and is not explained by inflammatory stress

The delayed imbalance occurred alongside heterogeneous remodeling of the broader endfoot and homeostatic network rather than uniform transcriptional loss (Figure 4A-C). At 6 mo, *Aqp4, Dmd, Dtna, Kcnj10, Gja1, Gjb6*, and *Slc1a2* were increased while *Snta1* remained below the Naive reference. Module trajectories likewise separated acute inflammatory stress activity from later endfoot and ionic support and glutamate-support remodeling.

**Figure 4.**
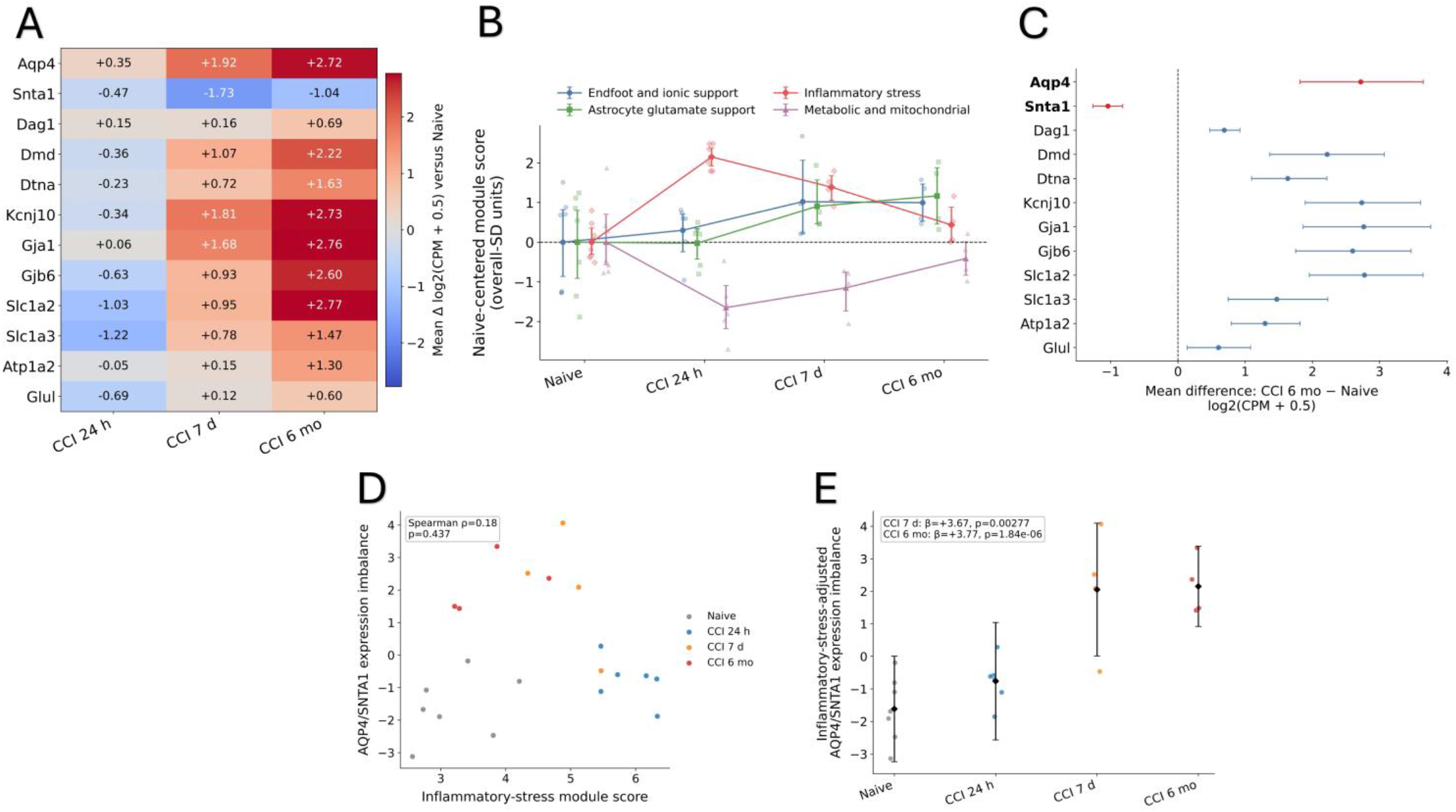
Endfoot and homeostatic transcriptional context of AQP4/SNTA1 expression imbalance after CCI. (A) Temporal expression changes in selected astrocyte endfoot and homeostatic genes across the GSE269748 CCI time course. Heatmap values represent mean expression change relative to each gene-specific Naive mean using log2(CPM + 0.5) values at CCI 24 h, 7 d, and 6 mo. Genes include *Aqp4, Snta1, Dag1, Dmd, Dtna, Kcnj10, Gja1, Gjb6, Slc1a2, Slc1a3, Atp1a2*, and *Glul*. The delayed response is heterogeneous, with increased *Aqp4* and multiple homeostatic genes occurring alongside persistently reduced *Snta1*. (B) Temporal trajectories of four validated astrocyte-associated transcriptional responses: endfoot and ionic support, astrocyte glutamate support, inflammatory stress, and metabolic and mitochondrial function. Validated Figure 1 scores are reused here and expressed as Naive-centered overall-SD units. Points represent individual biological samples; group summaries and bootstrap 95% confidence intervals are shown. Inflammatory stress activity was most prominent at 24 h, whereas endfoot and ionic support and glutamate support responses became more prominent at delayed stages, indicating temporally distinct components of astrocytic remodeling. (C) Chronic gene-level expression effects at CCI 6 mo relative to Naive. Points show mean differences in log2(CPM + 0.5), with horizontal bars indicating bootstrap 95% confidence intervals. *Aqp4* and *Snta1* are highlighted to emphasize their opposing chronic expression pattern. Multiple endfoot, ion-handling, gap-junction, and glutamate-support genes showed positive chronic effects, demonstrating that the AQP4/SNTA1 imbalance occurs within heterogeneous remodeling rather than uniform loss of the broader endfoot and homeostatic network. (D) Relationship between AQP4/SNTA1 expression imbalance and the validated inflammatory stress score across individual biological samples. Points are colored by CCI stage. The association was weak and not statistically significant (Spearman ρ = 0.18, p = 0.437); the fitted line is shown for visualization. Thus, variation in the measured inflammatory stress response alone did not account for the AQP4/SNTA1 expression imbalance. (E) AQP4/SNTA1 expression imbalance after statistical adjustment for inflammatory stress activity. Individual samples are shown together with model-adjusted stage estimates and 95% confidence intervals. The delayed imbalance remained evident after adjustment, including positive stage effects at CCI 7 d (β = +3.67, p = 0.00277) and CCI 6 mo (β = +3.77, p = 1.84 × 10^−6^). These results indicate that the delayed expression imbalance is not adequately explained by the measured inflammatory stress dimension, but they do not exclude a contribution of inflammatory biology.

AQP4/SNTA1 imbalance was only weakly related to the validated inflammatory stress score (Spearman ρ = 0.18, p = 0.437; Figure 4D), and delayed stage effects remained after inflammatory stress adjustment (7 d β = +3.67, p = 0.00277; 6 mo β = +3.77, p = 1.84 × 10^−6^; Figure 4E). Human CTE, acute severe TBI, and repetitive head impacts or early CTE datasets showed heterogeneous perturbation of the same broad homeostatic domain but did not reproduce a uniform cross-context state (Supplementary Figure 4). Thus, the imbalance was not adequately explained by generalized inflammatory reactivity alone.

### Potassium and ion homeostatic remodeling remains associated with AQP4/SNTA1 imbalance beyond broader reactivity

We next screened 14 prespecified transcriptional gene set scores representing biological processes for stageadjusted association with AQP4/SNTA1 imbalance. Four remained supported after permutation-FDR correction: hypoxia and angiogenic response (β = 1.164, q = 0.013), potassium and ion homeostasis (β = 0.962, q = 0.0014), endfoot anchoring and dystrophin-associated protein complex (DAPC; β = 0.831, q = 0.010), and astrocyte glutamate support (β = 0.797, q = 0.016; Figure 5A; Supplementary Figure 5A; Supplementary Table S38). Reactive gliosis and the remaining inflammatory, metabolic, oxidative-stress, and proteostasis processes were unsupported after correction.

**Figure 5.**
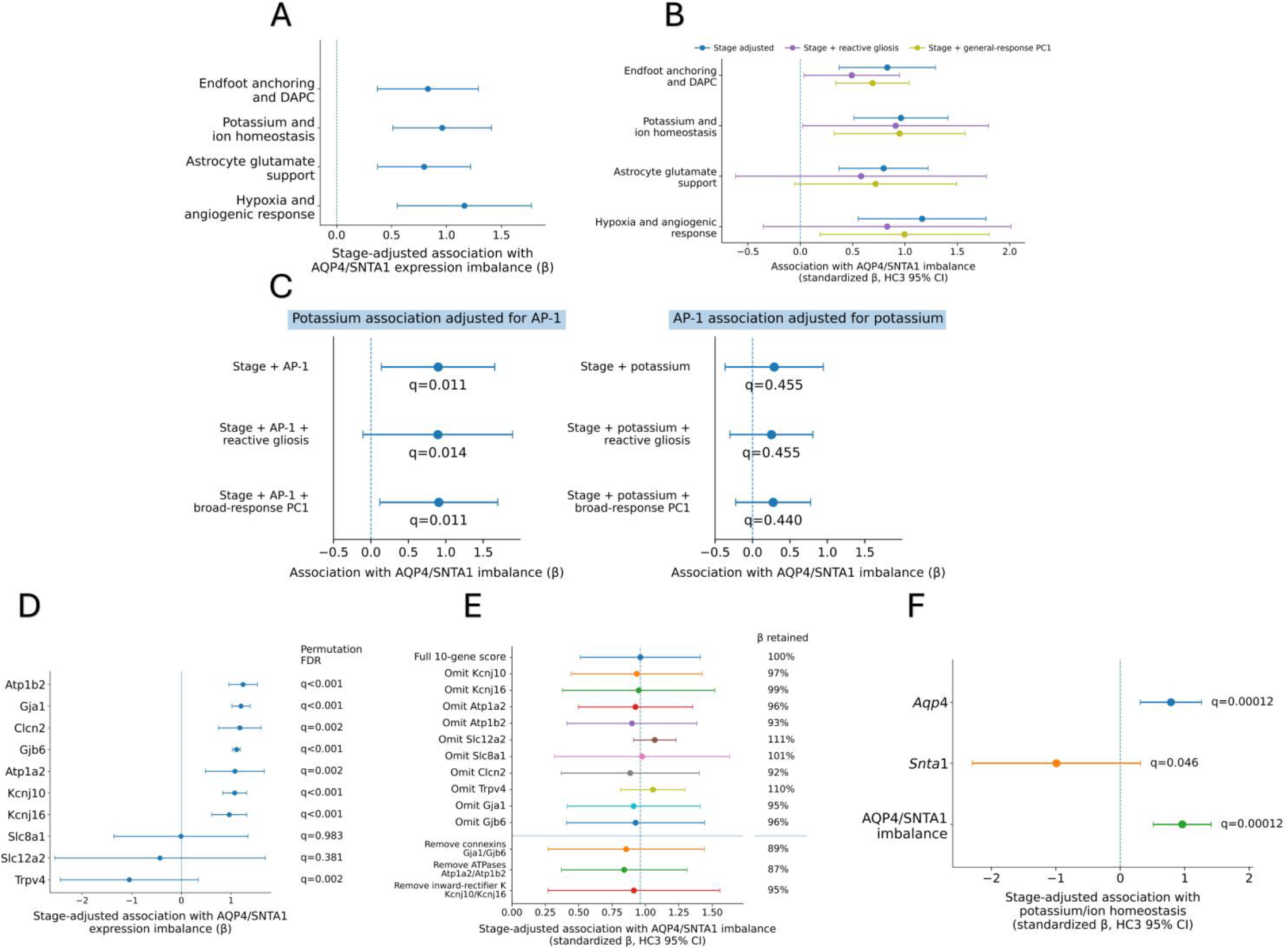
Specificity and robustness of potassium and ion homeostatic remodeling associated with AQP4/SNTA1 expression imbalance. (A) Stage-adjusted associations between AQP4/SNTA1 expression imbalance and the four biological processes that remained significant in the prespecified screen. Variables considering endfoot anchoring and DAPC, potassium and ion homeostasis, astrocyte glutamate support, and hypoxia and angiogenic response were positively associated with the imbalance after permutation-based multiple-testing correction. Points indicate standardized regression coefficients (β), horizontal bars indicate HC3 95% confidence intervals, and q values denote permutation-based FDR. (B) Specificity of the four supported associations after adjustment for broader reactive transcriptional structure. For each process, stage-adjusted estimates are compared with models additionally adjusted for reactive gliosis or the general-response PC1. Potassium and ion homeostasis showed particularly stable effect estimates across these adjustments, supporting its prioritization within the correlated homeostatic response. Covariate-adjusted associations indicate statistical specificity and do not establish biological independence or causality. (C) Reciprocal specificity analysis of potassium and ion homeostasis and AP-1 activity. Left, association of potassium and ion homeostasis with AQP4/SNTA1 expression imbalance after adjustment for AP-1 activity, with additional models incorporating reactive gliosis or the broad-response PC1. The potassium association remained supported across models (permutation q = 0.011, 0.014, and 0.011, respectively). Right, reciprocal association of AP-1 activity after adjustment for potassium and ion homeostasis. AP-1 was no longer independently supported after potassium adjustment (q = 0.455, 0.455, and 0.440, respectively). Points indicate standardized β coefficients and horizontal bars indicate HC3 95% confidence intervals. These analyses assess comparative statistical specificity and do not establish mediation, causal ordering, or direct transcriptional regulation. (D) Gene-level associations within the locked 10-gene potassium and ion homeostasis score comprising *Atp1b2, Gja1, Clcn2, Gjb6, Atp1a2, Kcnj10, Kcnj16, Slc8a1, Slc12a2*, and *Trpv4*. Several potassium-, ATPase-, chloride-, and connexin-associated genes showed positive associations with AQP4/SNTA1 imbalance, whereas *Slc8a1* was essentially null and *Slc12a2* and *Trpv4* showed negative estimates. Points indicate stage-adjusted β coefficients, horizontal bars indicate HC3 95% confidence intervals, and permutation-FDR values are shown at right. The heterogeneous gene-level pattern indicates that the score-level association does not reflect uniform regulation of all constituent genes. (E) Robustness of the potassium and ion homeostasis association to individual and grouped gene omission. The full 10-gene score was recalculated after sequential removal of each constituent gene and after removal of selected functional groups, including connexins (*Gja1*/*Gjb6*), Na+/K+-ATPase components (*Atp1a2*/*Atp1b2*), and inward-rectifier potassium channels (*Kcnj10*/*Kcnj16*). The association remained positive across all perturbations. Values at right indicate the percentage of the full-model β retained, demonstrating that the association was not driven by a single gene or narrow functional subgroup. (F) Parallel stage-adjusted models testing whether potassium and ion homeostasis was associated with Aqp4 expression, Snta1 expression, or the AQP4/SNTA1 expression imbalance. Points indicate standardized regression coefficients (β), horizontal bars indicate HC3 95% confidence intervals, and q values denote stage-stratified permutation FDR. Potassium and ion homeostasis was strongly associated with Aqp4 (β = 0.786, q = 0.00012) and with AQP4/SNTA1 imbalance (β = 0.962, q = 0.00012), whereas the Snta1-only model showed inconsistent support across inferential approaches (β = −0.989; HC3 p = 0.137; permutation q = 0.046). The imbalance model showed the largest partial R^2^, supporting the relative Aqp4-Snta1 expression relationship as the most informative summary of the potassium-associated transcriptional phenotype.

The four supported processes shared substantial stage-independent transcriptional structure, with pairwise residual correlations of 0.69-0.86 and a shared PC1 explaining 85.6% of residual variance (Supplementary Figure 5B; Supplementary Tables S39-S40). Potassium and ion homeostasis nevertheless showed the greatest statistical specificity: 95% of its stage-adjusted coefficient was retained after adjustment for reactive gliosis, 99% after broadresponse-PC1 adjustment, and 93% when both were included, with permutation-FDR support retained (Figure 5B; Supplementary Table S41).

Reciprocal models further distinguished potassium and ion homeostatic remodeling from AP-1 activity. The potassium association remained supported after AP-1 adjustment and additional reactivity covariates, whereas AP-1 was no longer independently supported after potassium adjustment (Figure 5C; Supplementary Table S42). Genelevel effects were heterogeneous, but the association remained positive after every single-gene omission and after removal of connexin, Na+/K+-ATPase, or inward-rectifier potassium-channel subsets (Figure 5D,E). Leave-onesample-out, bootstrap, strict-gate, and matched injury-model analyses also retained a positive association (Supplementary Figure 5; Supplementary Tables S43-S46). These analyses prioritize potassium and ion homeostasis as the most specific and internally robust transcriptional correlate of the imbalance, without establishing altered potassium flux or causality.

To determine whether this association was driven primarily by Aqp4, Snta1, or their relative expression imbalance, we fit three parallel stage-adjusted models (Figure 5F). Potassium and ion homeostasis was strongly associated with Aqp4 (standardized β = 0.786, HC3 p = 0.00125; permutation q = 0.00012) and with AQP4/SNTA1 imbalance (β = 0.962, HC3 p = 2.74 × 10^−5^; permutation q = 0.00012), whereas the Snta1-only model showed inconsistent support across inferential approaches (β = −0.989, HC3 p = 0.137; permutation q = 0.046). The imbalance model showed the largest partial R^2^ (0.775 versus 0.708 for Aqp4 and 0.463 for Snta1), indicating that the relative Aqp4-Snta1 expression relationship provided the most informative summary of the potassium-associated transcriptional phenotype. In a nested model containing Snta1 and the AQP4/SNTA1 imbalance, the imbalance remained independently associated with potassium and ion homeostasis, whereas Snta1 did not. An analogous nested model including Aqp4 and the AQP4/SNTA1 imbalance model was not interpreted because of severe multicollinearity between Aqp4 and the imbalance.

An independent astrocyte gate based only on Aldh1l1 and Sox9 preserved the direction of the potassium/ionhomeostasis association but substantially attenuated its magnitude and statistical support (β = 0.55, 95% CI −0.89 to 1.99; permutation p = 0.075). Count-matched downsampling suggested that reduced cell number could account for part, but not all, of this attenuation. Thus, this analysis supports directional robustness but not independent statistical confirmation.

### Temporal synthesis identifies delayed AQP4/SNTA1 and ion-homeostatic remodeling with defined external boundaries

The relationship between AQP4/SNTA1 expression imbalance and potassium and ion homeostasis was not confined to the primary CCI subgroup within GSE269748. In the matched 24-h dataset containing Naive, repetitive closed-head injury (rCHI), CCI, and CCI plus hemorrhagic shock samples, the injury-model-adjusted association remained strong (β = 0.98, 95% CI 0.61-1.35; permutation p < 0.001), with no supported model-by-imbalance interaction (p = 0.628; Figure 6A; Supplementary Table S48). Because the model-specific groups were small, this analysis supports robustness within GSE269748 rather than equivalence across injury paradigms or independent replication.

**Figure 6.**
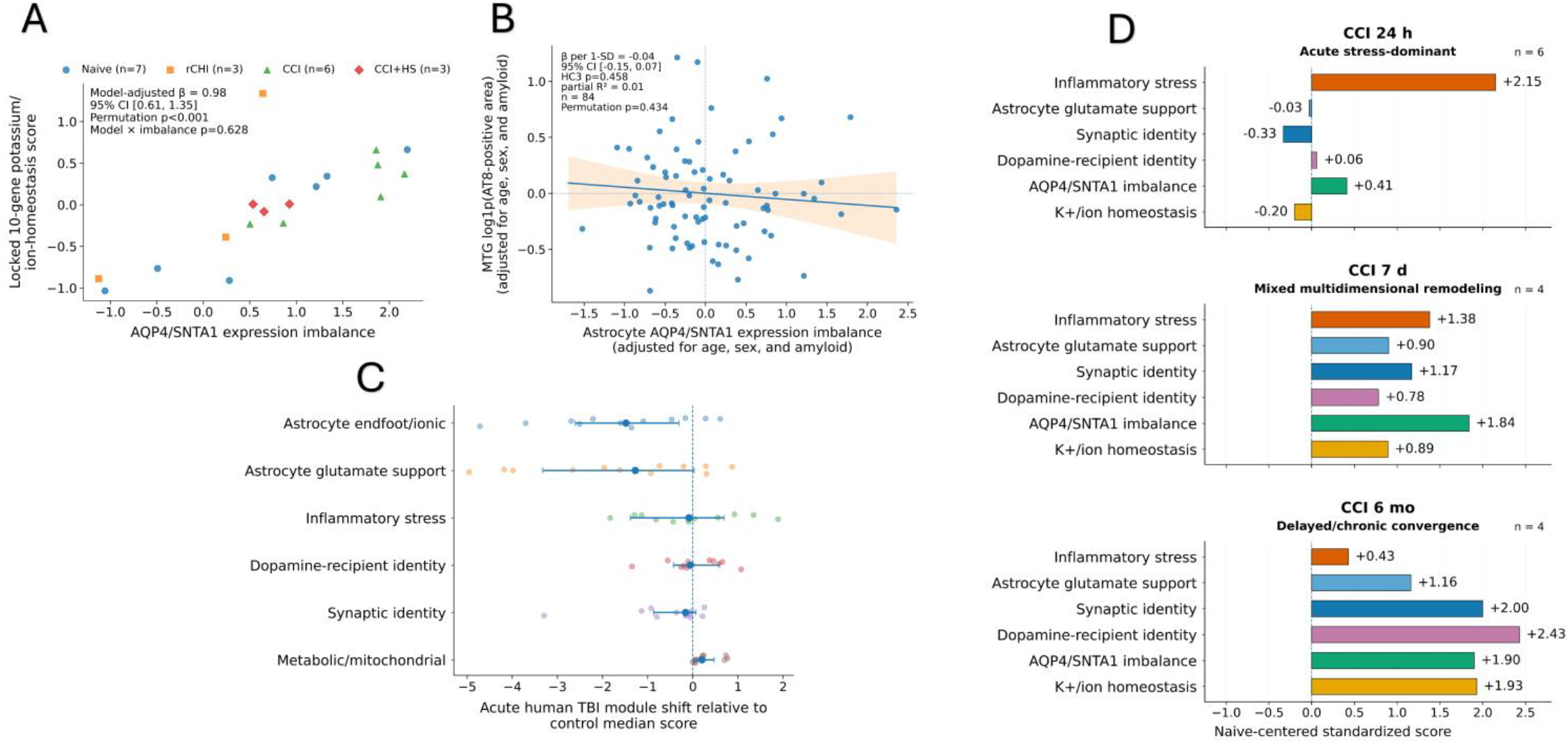
AQP4/SNTA1 imbalance and potassium and ion homeostatic remodeling across injury contexts and time. (A) Cross-injury-model robustness of the relationship between AQP4/SNTA1 expression imbalance and the locked 10-gene potassium and ion homeostasis score in the matched 24-h GSE269748 subset. Points represent individual biological samples from Naive (n = 7), rCHI (n = 3), CCI (n = 6), and CCI plus hemorrhagic shock (CCI+HS; n = 3). The association remained positive after adjustment for injury model (β = 0.98, 95% CI 0.61-1.35; permutation p < 0.001), with no supported model-by-imbalance interaction (p = 0.628). (B) Human neuropathology evidence boundary in SEA-AD middle temporal gyrus. Donor-level astrocyte AQP4/SNTA1 expression imbalance was tested for association with AT8-positive tau burden after adjustment for age, sex, and amyloid burden (n = 84). No association was detected (β per 1-SD = −0.04, 95% CI −0.15 to 0.07; HC3 p = 0.458; partial R^2^ = 0.01; permutation p = 0.434), indicating that the mouse-derived imbalance should not be interpreted as a general marker of greater human tau burden. (C) Acute human TBI contextual projection using GSE209552. Points show sample-level shifts in six transcriptional scores relative to the control median, with group summaries and uncertainty intervals. The largest negative shifts occurred in astrocyte endfoot and ionic support, as well as astrocyte glutamate support, whereas inflammatory stress, dopamine recipient, synaptic, and metabolic or mitochondrial responses were smaller. These data provide acute human context but do not directly replicate the delayed mouse CCI phenotype. (D) Integrated temporal configuration of six transcriptional responses in GSE269748. Bars show Naive-centered standardized scores at CCI 24 h, 7 d, and 6 mo. The 24-h profile was dominated by inflammatory stress, whereas the 7-d profile showed broader multidimensional remodeling. By 6 mo, AQP4/SNTA1 imbalance and potassium and ion homeostatic remodeling were prominent, as were changes to broader circuit-associated and astrocyte support responses. These profiles summarize transcriptional configurations and do not define discrete disease states.

Across the primary CCI time course, synaptic identity, dopamine-recipient identity, AQP4/SNTA1 imbalance, and potassium and ion homeostasis followed partially distinct stage-associated profiles (Supplementary Figure 6C). Synaptic identity was reduced acutely and increased at 7 d and 6 mo; dopamine-recipient identity and AQP4/SNTA1 imbalance became prominent at delayed stages; and potassium and ion homeostasis showed its strongest supported increase at 6 mo. All four delayed or chronic versus 24-h contrasts remained supported after global correction (q ≤ 0.0097; Supplementary Table S49). Formal profile testing rejected a single common trajectory (omnibus permutation p = 0.00247), with several pairwise profile differences remaining significant after correction (Supplementary Table S50).

The integrated six-response summary further illustrated this stage structure (Figure 6D). At 24 h, inflammatorystress activity was the strongest positive feature. By 7 d, broader remodeling was evident, including increased astrocyte glutamate support, synaptic identity, AQP4/SNTA1 imbalance, and potassium and ion homeostasis. At 6 mo, inflammatory stress activity was lower, whereas AQP4/SNTA1 imbalance and potassium and ion homeostatic remodeling remained prominent, as were broader circuit and astrocyte support-related changes. Post hoc correspondence with the unsupervised whole-transcriptome PCA was concordant with this organization. PC1 correlated strongly with AQP4/SNTA1 imbalance (ρ = 0.90), synaptic identity (ρ = 0.87), potassium and ion homeostasis (ρ = 0.82), and dopamine-recipient identity (ρ = 0.80), whereas inflammatory stress aligned more strongly with PC2 (ρ = 0.68; Supplementary Figure 6D). Because these transcriptional scores and PCA scores were derived from the same expression dataset, this analysis is interpreted as axis correspondence rather than independent validation.

The external datasets showed that this relationship was not consistently reproduced across contexts. In 84 SEAAD middle temporal gyrus donors, astrocyte AQP4/SNTA1 imbalance was not associated with AT8-positive tau burden after adjustment for age, sex, and amyloid burden (β per 1-SD = −0.04, 95% CI −0.15 to 0.07; HC3 p = 0.458; permutation p = 0.434; Figure 6B; Supplementary Figure 6A; Supplementary Tables S52-S53). Acute human TBI in GSE209552 showed the largest negative shifts in astrocyte endfoot and ionic support as well as astrocyte glutamate support, emphasizing the importance of injury stage and context (Figure 6C; Supplementary Table S51). In independent astrocytes isolated 365 d after TBI in the PRJNA1280541 dataset, neither the AQP4/SNTA1 imbalance nor its association with potassium and ion homeostasis were replicated, although the potassium score showed limited directional support (Supplementary Figure 6B; Supplementary Table S54). Spatial analyses in GSE282909 likewise showed non-random anatomical organization of individual transcriptional responses but weak local correspondence between AQP4/SNTA1 imbalance and potassium and ion homeostasis (Supplementary Figure 7A-D; Supplementary Tables S55-S58). Together, the data support a delayed AQP4/SNTA1 transcriptional imbalance that is most consistently associated with ion homeostatic remodeling in the discovery dataset, while the external analyses clearly suggest that this relationship is context-dependent.

## Discussion

This study supports a model of post-traumatic molecular remodeling in which acute and chronic CCI are not simply different magnitudes of a single injury response. Acute injury was dominated by stress-associated transcription, whereas delayed stages showed broader remodeling involving astrocyte support, circuit-associated responses, AQP4/SNTA1 imbalance, and ion homeostasis. Together, the stage-associated profiles support changing molecular configurations across the post-injury course rather than a linear progression of a single response.

The astrocyte findings provide a more focused example of this temporal reorganization. Contemporary frameworks reject binary definitions of astrocyte reactivity [41], and recent TBI studies show that astrocyte states continue to change into chronic injury [12,13]. Here, delayed endfoot and ionic remodeling and glutamate-support changes were poorly explained by inflammatory stress-related activity alone. Within this broader response, disproportionate *Aqp4* relative to *Snta1* emerged at 7 d and persisted at 6 mo as a focused transcriptional phenotype. The established alpha-syntrophin AQP4 literature provides strong biological context: alpha-syntrophin supports perivascular AQP4 localization [14-16] and its loss alters AQP4 organization, potassium clearance, and perivascular solute exchange [17,42,43]. TBI can also dissociate overall AQP4 abundance from perivascular localization [18-20]. Our data do not demonstrate these structural or physiological outcomes, but they identify a delayed expression configuration involving the same molecular axis.

Potassium and ion homeostatic remodeling provided the strongest specific transcriptional context for the imbalance. The association emerged from a prespecified screen, persisted after accounting for shared transcriptional structure, reactive gliosis, and AP-1 activity, and was stable to gene, sample, and injury model perturbations, while the independent minimal-marker gate produced a directionally consistent but attenuated and statistically inconclusive estimate. Parallel models further showed that the AQP4/SNTA1 imbalance explained more of the potassium-associated variation than either *Aqp4* or *Snta1* alone. Gene-level heterogeneity argues against uniform induction of a single pathway and instead points to distributed remodeling across channels, transporters, connexins, and ATPase-related components. The known relationship between alpha-syntrophin-dependent AQP4 organization and extracellular K+ clearance [17] makes ion handling a plausible experimental consequence, but the present data do not establish altered potassium concentrations, buffering, or causal ordering.

The circuit analyses provide complementary context rather than a separate mechanistic conclusion. Human imaging and pharmacological studies establish dopaminergic relevance in selected TBI phenotypes [4-8], but our analyses showed that dopamine recipient changes occurred within broader shifts in multi-neurotransmitter and synaptic remodeling. At 6 months, positive transcriptional enrichment extended across GABAergic, glutamatergic, dopaminergic, serotonergic, adrenergic, and cholinergic systems, while purinergic/adenosine-associated transcription followed a distinct, negative trajectory. This distributed remodeling provides a potential molecular context for the heterogeneous efficacy of neurotransmitter-directed therapies after TBI. It does not indicate that dopaminergic interventions are ineffective or explain individual treatment responses, but it suggests that dopamine-associated dysfunction may represent one component of a broader chronic circuit phenotype. Adjustment for the dominant sample-level composition axis preserved the overall stage effects and the chronic signal, while earlier coefficients were more composition-sensitive. Thus, the data support broad circuit-associated transcriptional remodeling rather than increased dopamine release, receptor activation, or a neuron-intrinsic dopaminergic state.

The external datasets did not provide consistent validation of these findings. Acute human TBI showed substantial astrocyte-support perturbation, whereas the independent one-year mouse astrocyte dataset did not replicate the AQP4/SNTA1 imbalance or its potassium association. SEA-AD provided no evidence that higher astrocyte imbalance predicts greater tau burden, and spatial analyses showed anatomical organization of individual transcriptional responses without strong local correspondence between AQP4/SNTA1 imbalance and potassium and ion homeostasis. These findings argue against a universal chronic injury signature and emphasize dependence on injury model, region, timepoint, tissue compartment, and analytical strategy. The available external evidence therefore provides limited contextual support rather than independent validation.

Several limitations follow from this design. The principal temporal and specificity analyses derive from one public discovery cohort with small delayed-stage groups. The astrocyte-like pseudobulk preserves animal-level inference and excludes *Aqp4* and *Snta1* from the positive gate, but it is not equivalent to experimentally isolated perivascular astrocytes. The all-QC circuit compartment can reflect both within-cell regulation and cell composition despite composition-adjusted sensitivity analyses. Timepoints are cross-sectional, not repeated measurements in the same animals. The primary discovery analysis was restricted to male mice by metadata, so the extent to which these transcriptional relationships generalize to females remains unknown. In addition, because the chronic cohort differed from the earlier cohorts in both age at tissue collection and CCI parameters, differences observed at 6 mo should not be attributed exclusively to post-injury timepoint. Most robustness tests remain internal to GSE269748, and external datasets differ substantially in model and sampling context. Finally, transcript abundance cannot establish protein abundance, protein-protein interaction, membrane localization, neurotransmission, potassium physiology, glymphatic exchange, or behavioral consequences.

These limitations define a focused experimental path forward. The immediate priority is time-resolved protein-level validation of AQP4, alpha-syntrophin, and KCNJ10 at vascular-associated astrocyte endfeet after CCI, followed by direct measurement of extracellular potassium-clearance kinetics if structural reorganization is confirmed. Causal testing would then require manipulation of *Snta1* or related endfoot machinery. More broadly, the present study identifies delayed AQP4/SNTA1 expression imbalance within potassium and ion homeostatic remodeling as a specific, falsifiable feature of chronic post-traumatic astrocyte biology while placing that phenotype within the broader temporal evolution of post-CCI molecular responses.

## Conclusion

CCI engages partially distinct molecular responses whose relative prominence changes across the post-injury course. Acute stress-associated transcription gives way to delayed astrocyte-homeostatic remodeling, including a sustained AQP4/SNTA1 expression imbalance and prominent potassium and ion homeostatic changes. The AQP4/SNTA1 imbalance was most specifically associated with ion-homeostatic transcription in the primary dataset, whereas the mouse, human, neuropathological, and spatial external analyses showed that this relationship was context dependent. Direct spatial protein and physiological studies are required to determine whether the transcriptional phenotype translates into altered perivascular endfoot organization or potassium handling.

## Materials and methods

### Study design and primary discovery dataset

This study reanalyzed publicly available transcriptomic, proteomic, and spatial datasets. GSE269748 was the primary discovery cohort. From 36 GEO samples, the primary temporal analysis was restricted a priori by metadata to the 21-sample male focal CCI/Naive series: Naive (n = 7), CCI 24 h (n = 6), CCI 7 d (n = 4), and CCI 6 mo (n = 4). Mice in the source study were 12 to 15 weeks old at experimental assignment and were therefore of comparable age at the time of CCI; however, age at tissue collection differed across post-injury timepoints. The remaining 15 samples represented a separate female cohort, CCI plus hemorrhagic shock, contralateral CCI tissue, or repetitive closed-head injury and were not excluded on the basis of expression. Biological animals, human donors, or original biological samples, rather than cells, nuclei, or spatial spots, were the unit of group-level inference. GSE269748 was used to define the primary temporal and association structure, whereas additional mouse and human datasets were used for prespecified replication, contextual support, sensitivity analysis, or evidence-boundary testing according to their experimental compatibility with the discovery analysis. Full dataset provenance, sample inclusion criteria, and analytical roles are provided in the supplementary tables.

### Astrocyte-like and all-QC pseudobulk construction

For astrocyte-focused analyses, cells that passed quality control were selected using an astrocyte-support score based on *Aldh1l1, Sox9, Slc1a3, Glul, Gja1, Gjb6*, and *Gfap*, together with an exclusion-lineage score. Neither *Aqp4* nor *Snta1* was included in the positive gate. Raw counts from retained cells were summed within each animal to generate astrocyte-like pseudobulk profiles, preserving the animal as the unit of inference. A stricter gate was used as a sensitivity analysis. To test whether overlap between selection markers and homeostatic gene sets could create circularity, an additional independent gate was reconstructed using only *Aldh1l1* and *Sox9*, followed by count-matched downsampling of the primary gate. Circuit analyses used matched all-QC animal-level pseudobulk without astrocyte selection, providing a broader sample-level representation of circuit-associated transcriptional remodeling.

### Differential expression and unsupervised transcriptomic structure

Analyses were conducted in R. Differential expression used edgeR with filterByExpr, trimmed mean of M-values normalization, robust quasi-likelihood inference, and Benjamini-Hochberg correction [29-32]. Whole-transcriptome principal component analysis and sample-correlation analyses were used as descriptive assessments of unsupervised structure. Complete R and package versions are reported in Supplementary Methods and the reproducibility package.

### Transcriptional gene sets and circuit analyses

Prespecified neural, inflammatory, astrocyte support, endfoot and ionic, metabolic, and neurotransmitter transcriptional gene sets were scored from sample-level expression in R using locked gene definitions. Circuit enrichment used fgsea and MSigDB collections [33-35], followed by systematically derived non-overlapping transmitter modules and targeted synaptic identity and dopamine recipient scores. Outcome-independent cell clustering was used to derive sample-level composition principal components [39]; the dominant composition axis was included in sensitivity models for the targeted synaptic and dopamine recipient scores. The targeted genes were excluded from the clustering features used to derive composition PCs.

### AQP4/SNTA1 expression imbalance

AQP4/SNTA1 expression imbalance was defined as log2(*Aqp4* CPM + 0.5) minus log2(*Snta1* CPM + 0.5), with higher values indicating greater *Aqp4* relative to *Snta1* expression. Alternative difference and z-score formulations and a Naive-reference residual analysis tested metric robustness. Targeted stage comparisons used biological-sample permutation tests and bootstrap confidence intervals.

### Specificity and robustness analyses

Fourteen prespecified biological processes were screened for stage-adjusted association with standardized AQP4/SNTA1 imbalance using HC3 regression and stage-stratified permutation inference. AQP4 and SNTA1 were excluded from pathway score construction. Potassium and ion homeostasis specificity was evaluated after adjustment for reactive gliosis, a broad-response principal component, and inferred AP-1 activity. Transcriptionfactor activity inference used established regulon-based approaches implemented through decoupleR/CollecTRI resources [36-38]. Robustness analyses included single-gene and functional-subgroup omission, leave-one-sample-out analysis, bootstrap resampling, strict-gate and independent-gate sensitivity analyses, count-matched downsampling, and cross-injury-model analysis. Parallel stage-adjusted models compared associations of potassium and ion homeostasis with *Aqp4, Snta1*, and the AQP4/SNTA1 imbalance. Temporal profile permutation testing compared the major molecular dimensions across stages, and predefined transcriptional scores were related post hoc to the unsupervised PCA axes.

### External datasets

External datasets were analyzed for replication, contextual support, or evidence boundary purposes according to their respective experimental compatibility with the discovery analysis dataset (GSE269748). These included GSE180862, GSE155114, PXD007694, GSE160763, GSE261807, GSE209552, GSE276182, GSE276422, PRJNA1280541, SEA-AD middle temporal gyrus resources, and GSE282909 [9,12,13,21-28]. Mouse, human, proteomic, perturbational, neuropathological, and spatial datasets were retained even when they did not reproduce the discovery phenotype.

### Statistical framework and interpretation boundaries

Exact or permutation-based inference was preferred for small groups. Regression models used HC3 robust covariance to provide coefficient uncertainty estimates. For stage-adjusted association analyses, stage-stratified permutation tests constituted the primary small sample inferential framework, with Benjamini-Hochberg falsediscovery-rate correction applied within the relevant testing families; HC3 p values were retained as complementary regression-based inference. Moran’s I was used for spatial autocorrelation where indicated [40]. Cross-sectional stage summaries were not interpreted as longitudinal trajectories in the same animals. Transcriptomic associations were not interpreted as direct measures of protein localization, neurotransmission, potassium flux, glymphatic function, or causality.

## Additional information

### Author contributions

Conceptualization: BG; Methodology: BG; Formal analysis: BG; Investigation: BG; Data curation: BG; Visualization: BG; Writing – original draft: BG; Writing – review and editing: BG, QZ, EBE; Supervision: BG, QZ, EBE; Project administration: QZ; Funding acquisition: QZ,. All authors reviewed and approved the final manuscript.

