## Supplementary Figure for "Temporal transcriptomic remodeling after controlled cortical impact reveals delayed AQP4/SNTA1 expression imbalance associated with ion-homeostatic remodeling"

### SUPPLEMENTARY FIGURES

*Traumatic brain injury transcriptomic study*

**Supplementary Figure 1. Study design, evidence framework, and robustness of temporal transcriptomic organization after CCI**

**A**

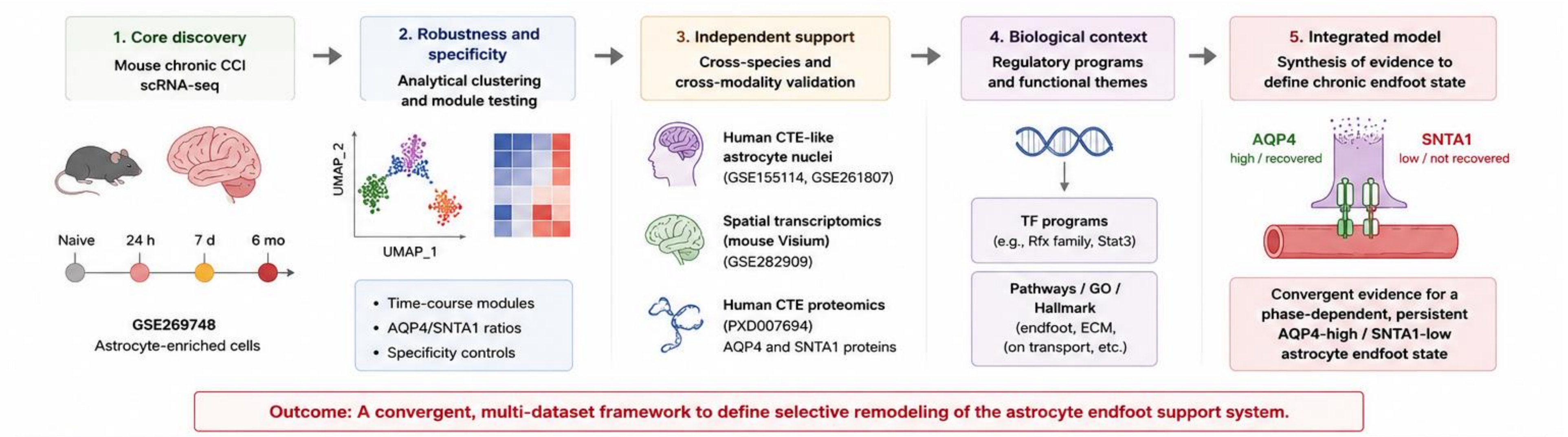


**B**

| Dataset | Species | Model / Context | Region / Tissue | Modality | Timepoints / Stage | Role in study | Evidence tier |
|---|---|---|---|---|---|---|---|
| GSE269748 | Mouse | CCI, rCHI | Cortex | scRNA-seq (10x) | Naive, 24 h, 7 d, 6 mo | Core discovery (timecourse) | ● Core |
| GSE180862 | Mouse | mTBI | Cortex | scRNA-seq (10x) | 24 h, 7 d | Independent mouse validation | ● Supporting |
| GSE155114 | Human | CTE | Frontal white matter | snRNA-seq (10x) | Chronic disease state | Main human validation (astrocytes) | ● Core |
| GSE209552 | Human | Severe TBI | Injured brain tissue | Bulk RNA-seq | Acute | Acute human context | ● Contextual |
| GSE104687 | Human | Postmortem TBI | Multi-region | Bulk RNA-seq | Chronic | Supplementary human context | ● Supporting |
| GSE261807 | Human | RHI / early CTE | Cortex | snRNA-seq (10x) | Chronic / early disease | Supplementary human context | ● Supporting |
| GSE282909 | Mouse | mTBI (CHIMERA Visium) | Brain sections | Spatial transcriptomics (Visium) | 7 d | Spatial validation of endfoot program | ● Supporting |
| PXD007694 | Human | CTE | Frontal cortex | Proteomics (DIA-MS) | Chronic disease state | Protein-level validation (AQP4, SNTA1) | ● Core |

● Core evidence (essential to the main conclusions) ● Supporting evidence (strengthens findings) ● Contextual evidence (provides biological context)

**C**

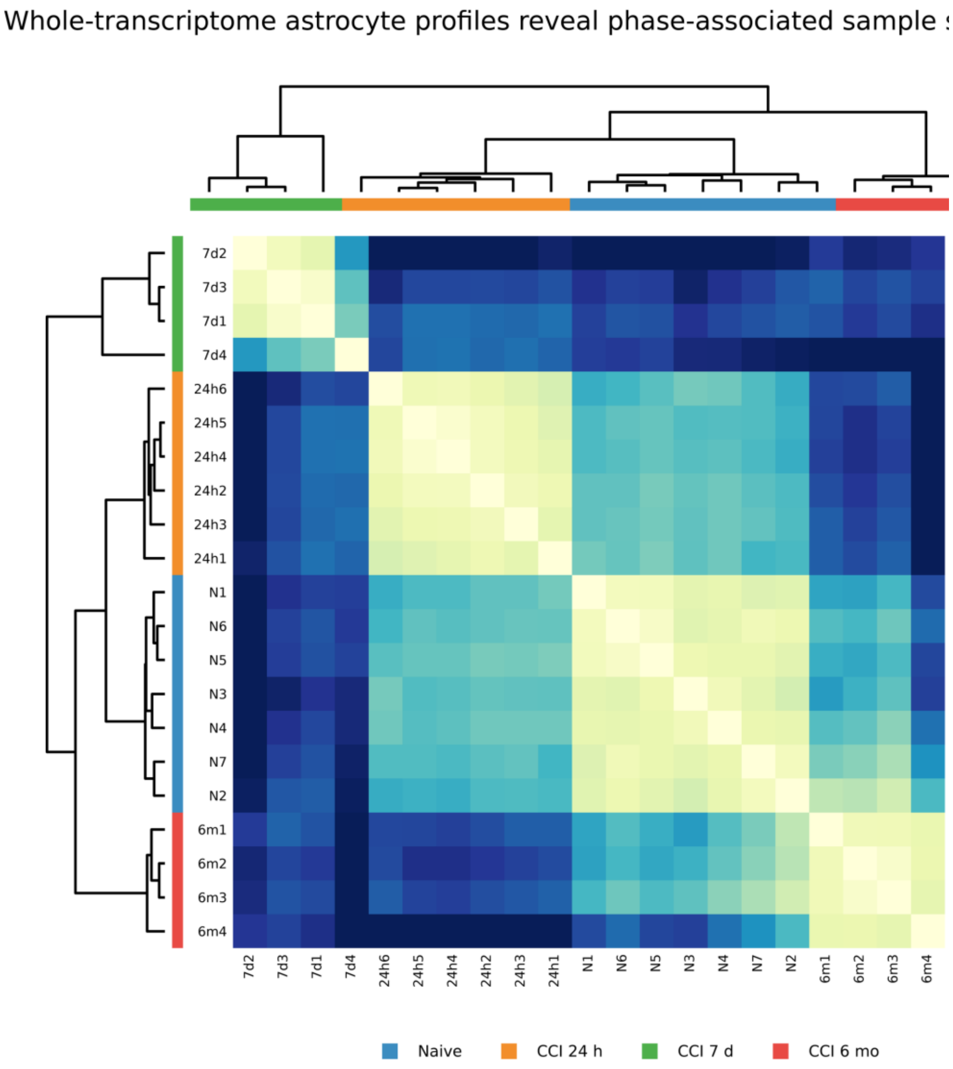
Whole-transcriptome astrocyte profiles reveal phase-associated sample
7d2
7d3
7d1
7d4
24h6
24h5
24h4
24h2
24h3
24h1
N1
N6
N5
N3
N4
N7
N2
6m1
6m2
6m3
6m4
Naive
CCI 24 h
CCI 7 d
CCI 6 mo

**D**

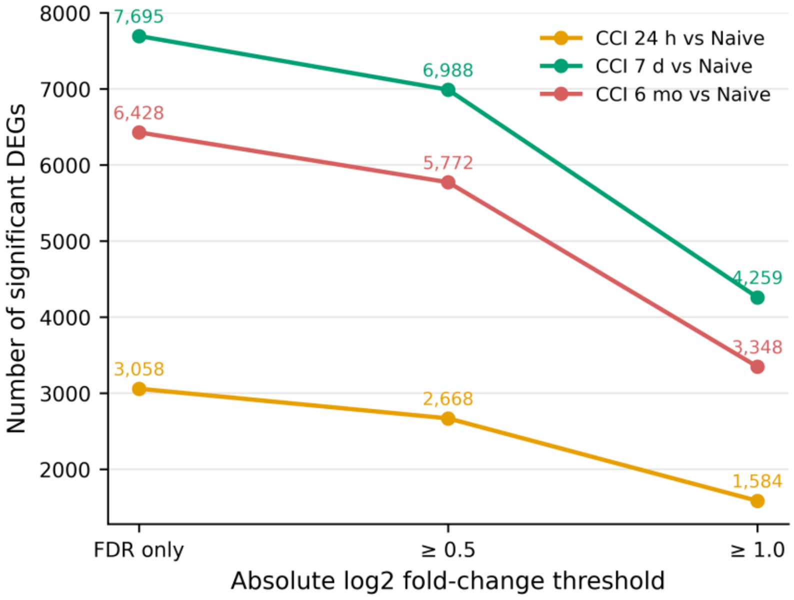
CCI 24 h vs Naive
CCI 7 d vs Naive
CCI 6 mo vs Naive
7,695
6,988
4,259
6,428
5,772
3,348
3,058
2,668
1,584
8000
7000
6000
5000
4000
3000
2000
Number of significant DEGs
FDR only
≥ 0.5
≥ 1.0
Absolute log2 fold-change threshold

E

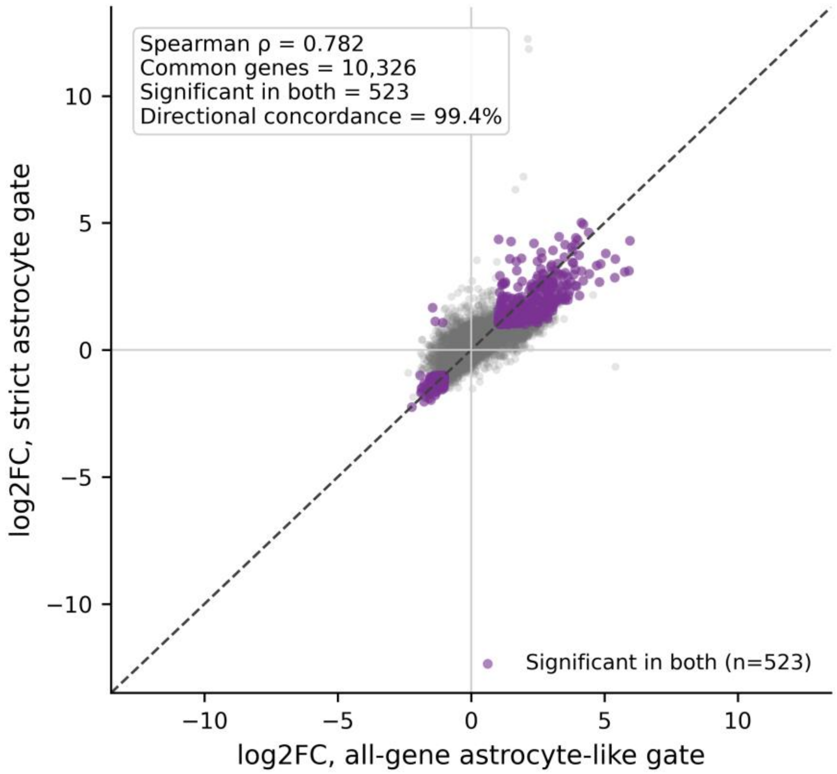


#### Legend

(A) Overview of the multi-level analytical strategy used to define temporal transcriptional remodeling after controlled cortical impact (CCI), test the robustness and specificity of the AQP4/SNTA1 expression imbalance, and evaluate its biological context and generalizability across complementary datasets. GSE269748 served as the primary temporal discovery cohort spanning Naive, 24 h, 7 d, and 6 mo after CCI. Internal robustness analyses included alternative astrocyte gates, gene- and sample-omission analyses, composition adjustment, count-matched downsampling, and cross-injury-model testing. External mouse, human, proteomic, neuropathological, and spatial datasets were used as replication tests, contextual-support datasets, or evidence boundaries according to their comparability with the discovery analysis. These analyses converged on delayed AQP4/SNTA1 transcriptional imbalance as a discovery-cohort phenotype most consistently associated with potassium and ion homeostatic remodeling, while also defining limits to its generalization across injury models, stages, tissues, and modalities.

(B) Inventory of datasets incorporated into the study, including species, injury or disease context, tissue, molecular modality, sampling stage, analytical role, and evidence category. The evidence hierarchy distinguishes the primary discovery dataset from internal robustness analyses, external contextual support, and datasets used to test boundaries of generalizability. External datasets were retained regardless of whether they reproduced the discovery phenotype and therefore should not be interpreted uniformly as independent validation cohorts.

(C) Complementary unsupervised sample-similarity analysis for the same 21 GSE269748 astrocyte-like pseudobulk profiles used in Figure 1. The Pearson sample-correlation matrix and hierarchical clustering provide a second representation of stage-associated transcriptomic organization alongside the whole-transcriptome PCA. Because both analyses use the same underlying expression matrix, this panel is a robustness view of sample structure rather than independent validation (Supplementary Table S07).

(D) Differential-expression threshold sensitivity. The common edgeR testing universe contained 16,209 genes. At FDR $< 0.05$ without a fold-change threshold, the numbers of significant genes were 3,058 at 24 h, 7,695 at 7 d, and 6,428 at 6 mo. With $|\log2FC| \geq 0.5$, the corresponding counts were 2,668, 6,988, and 5,772. With the primary large-effect threshold of FDR $< 0.05$ and $|\log2FC| \geq 1$, the counts were 1,584, 4,259, and 3,348. Thus, the ordering 7 d > 6 mo > 24 h was preserved across all three definitions, showing that the non-monotonic DEG-burden result is not dependent on a single effect-size threshold (Supplementary Table S05).

(E) Chronic astrocyte-selection sensitivity for the matched Naive versus CCI 6-mo comparison. Gene-level log2 fold-change estimates from the primary astrocyte-like pseudobulk and the stricter astrocyte gate were correlated across 10,326 common tested genes (Spearman $\rho = 0.782$). Of 523 genes significant in both analyses, 520 showed concordant directions, corresponding to 99.4% directional concordance. This analysis supports robustness of the chronic transcriptional signal to astrocyte-selection stringency, but it is restricted to the Naive-versus-6-mo contrast and is not a replication of the complete time course (Supplementary Table S06).

**Supplementary Figure 2. Targeted circuit-module robustness and cellular-composition sensitivity**

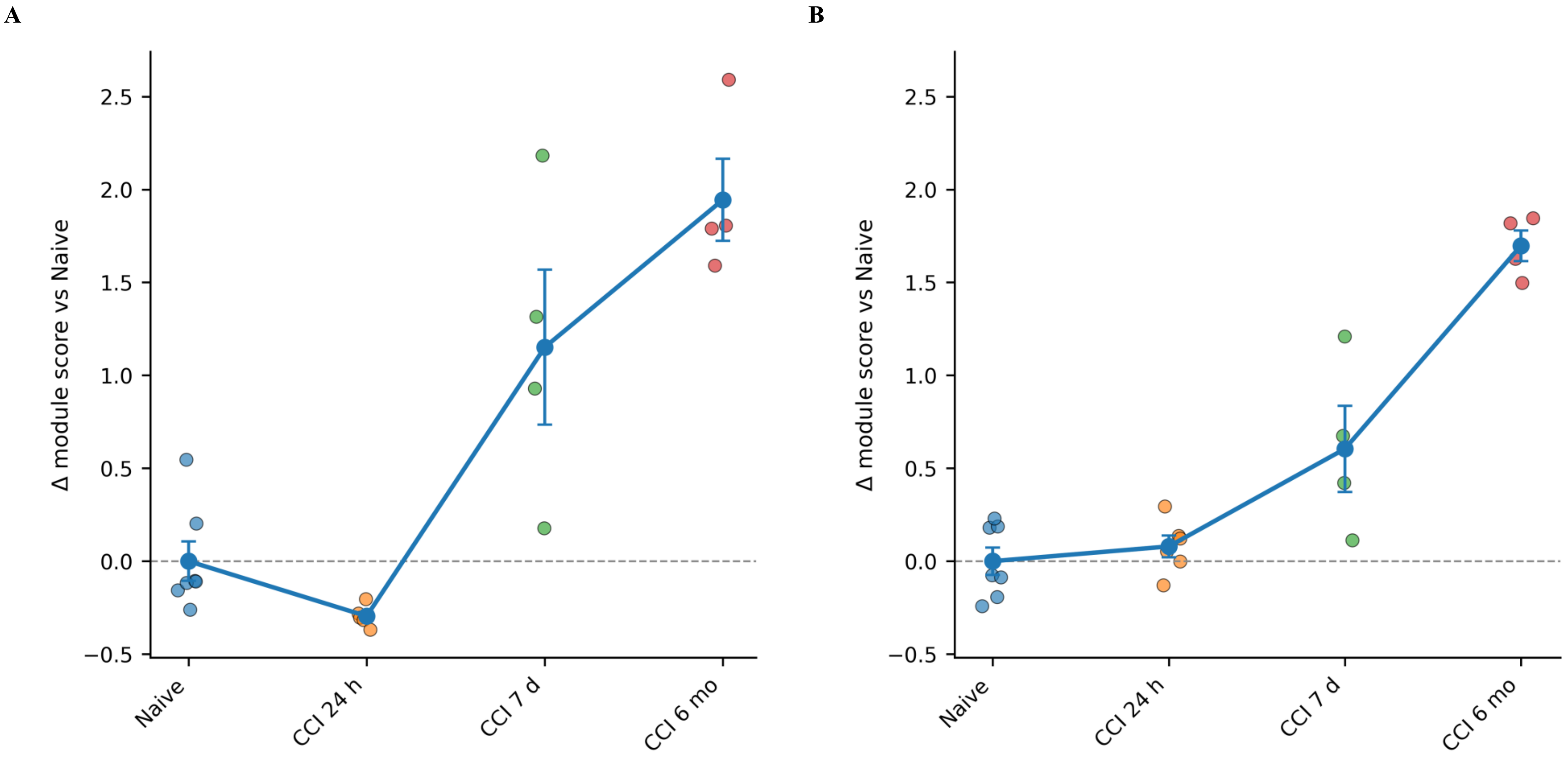

C

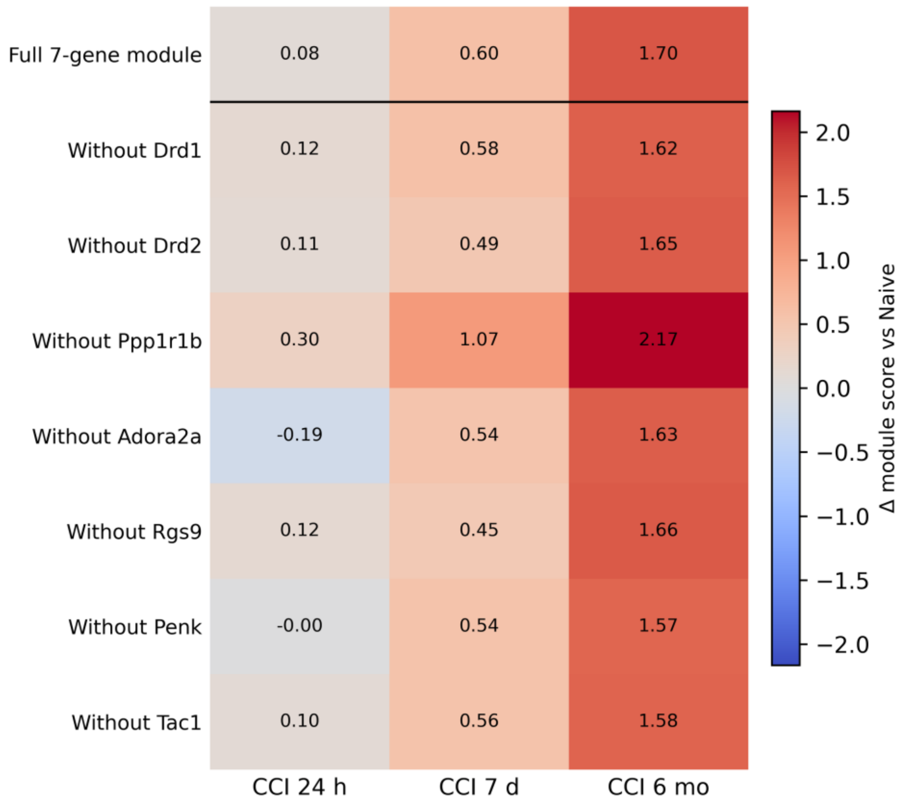
Full 7-gene module
Without Drd1
Without Drd2
Without Ppp1r1b
Without Adora2a
Without Rgs9
Without Penk
Without Tac1
0.08 0.60 1.70
0.12 0.58 1.62
0.11 0.49 1.65
0.30 1.07 2.17
-0.19 0.54 1.63
0.12 0.45 1.66
-0.00 0.54 1.57
0.10 0.56 1.58
CCI 24 h
CCI 7 d
CCI 6 mo
Δ module score vs Naive

D

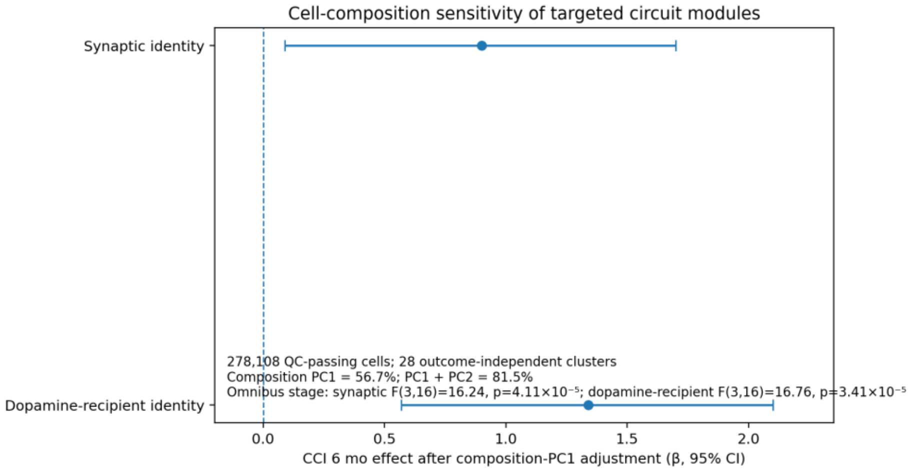
Cell-composition sensitivity of targeted circuit modules
Synaptic identity
Dopamine-recipient identity
278,108 QC-passing cells; 28 outcome-independent clusters
Composition PC1 = 56.7%; PC1 + PC2 = 81.5%
Omnibus stage: synaptic F(3,16)=16.24, p=4.11×10⁻⁵; dopamine-recipient F(3,16)=16.76, p=3.41×10⁻⁵
CCI 6 mo effect after composition-PC1 adjustment (β, 95% CI)

**E**

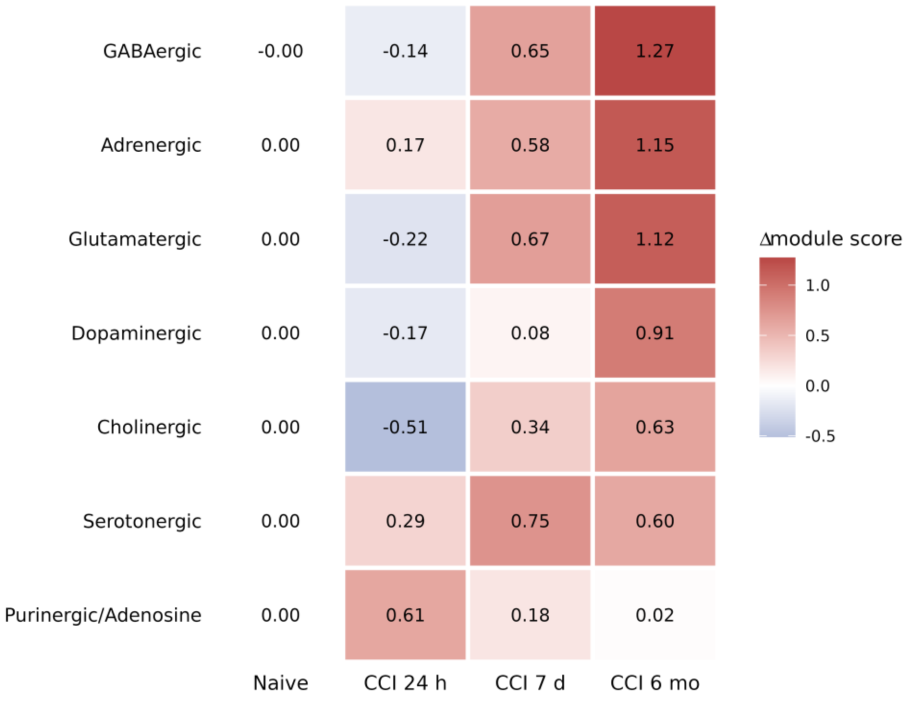


**F**

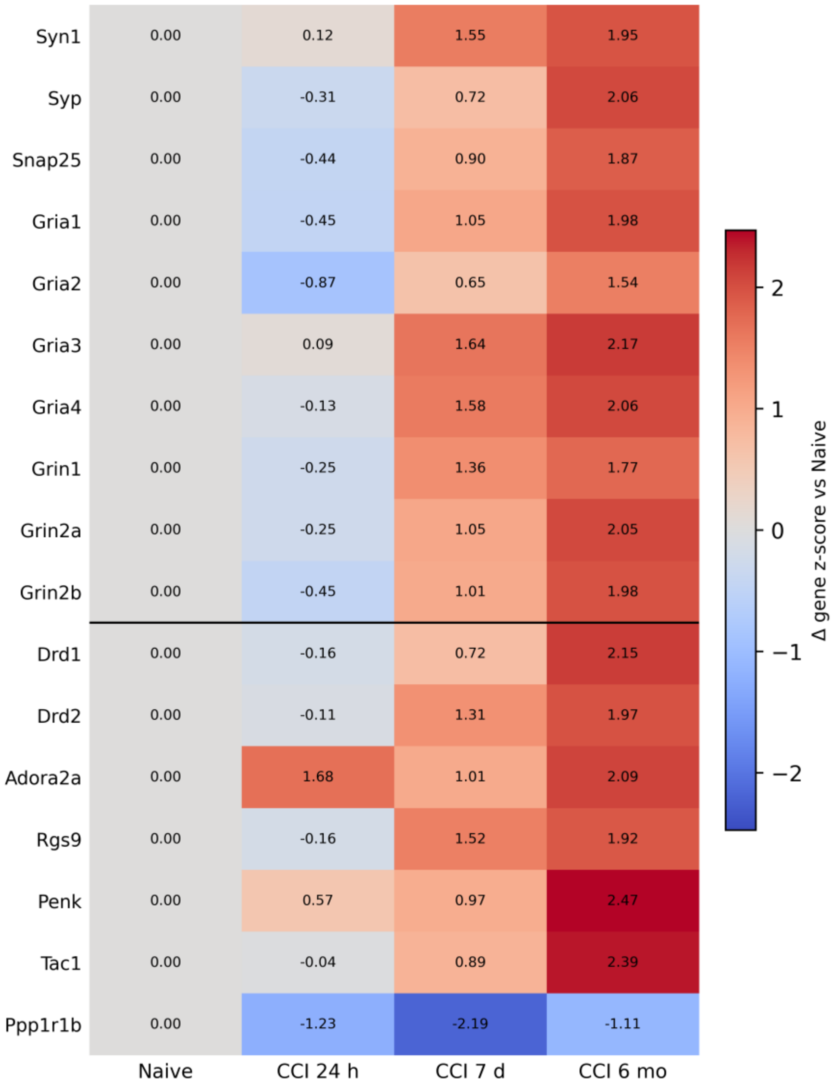


#### Legend

(A,B) Prespecified targeted module trajectories in the all-QC pseudobulk compartment. The 10-gene synaptic-identity module (Syn1, Syp, Snap25, Gria1, Gria2, Gria3, Gria4, Grin1, Grin2a, Grin2b) shifted from -0.30 relative to Naive at 24 h to +1.15 at 7 d and +1.94 at 6 mo. The seven-gene dopamine-recipient module (Drd1, Drd2, Ppp1r1b, Adora2a, Rgs9, Penk, Tac1) shifted from +0.08 at 24 h to +0.60 at 7 d and +1.70 at 6 mo. These are prespecified targeted follow-up scores, not unbiased pathway discoveries and not measurements of neurotransmitter release, receptor activation, or functional circuit activity (Supplementary Table S17).

(C) Leave-one-gene-out robustness of the dopamine-recipient module. Each version was rescored after omitting one constituent gene and re-centered to that version's own Naive reference. The full seven-gene module and all seven single-gene-omission versions remained positive at CCI 6 mo (8/8 positive versions), demonstrating that the chronic signal was distributed across the module rather than driven by a single dopamine-associated transcript (Supplementary Table S18).

(D) Cellular-composition sensitivity of the targeted circuit modules. The same 21 animals were independently reprocessed using all 278,108 QC-passing cells. The 17 genes constituting the synaptic and dopamine-recipient outcomes were excluded from feature selection and clustering, and outcome-independent clustering generated 28 clusters. Composition PC1 explained 56.7% of sample-level composition variance and PC1 plus PC2 explained 81.5%. After adjustment for composition PC1, the overall stage association remained supported for synaptic identity (HC3 $F[3,16] = 16.24$, $p = 4.11 \times 10^{-5}$) and dopamine-recipient identity ($F[3,16] = 16.76$, $p = 3.41 \times 10^{-5}$). The 6-mo coefficients remained positive for synaptic identity ($\beta = 0.90$, 95% CI 0.09 to 1.70, $p = 0.031$) and dopamine-recipient identity ($\beta = 1.34$, 95% CI 0.57 to 2.10, $p = 0.00195$), whereas the 24-h and 7-d coefficients were attenuated and unsupported. A secondary PC1+PC2 model retained significant omnibus stage effects but had substantial multicollinearity (maximum VIF approximately 53), so individual coefficients from that model were not strongly interpreted. An independently defined neuronal-like gate was also audited before outcome inspection, but one animal retained as few as 10 neuronal-like cells; neuron-only animal-level pseudobulk was therefore considered insufficiently stable for inference and no neuron-intrinsic claim is made (Supplementary Table S18).

Broader Figure 2 context retained in the supplementary tables: at 24 h, dopamine, serotonin, and glutamate neurotransmitter-release pathways were negatively enriched (dopamine NES = -2.09, FDR = 0.0010; serotonin NES = -1.86, FDR = 0.0165; glutamate NES = -1.81, FDR = 0.0191), whereas purinergic/adenosine was the only transmitter system with supported positive enrichment (NES = 1.73, FDR = 0.0016). At 7 d, GABAergic (NES = 1.73, FDR = $3.5 \times 10^{-4}$) and glutamatergic (NES = 1.61, FDR = $3.3 \times 10^{-4}$) enrichment was supported, while dopaminergic enrichment remained modest (NES = 1.06, FDR = 0.49). At 6 mo, GABAergic (NES = 2.41), glutamatergic (2.09), dopaminergic (1.88), serotonergic (1.80), adrenergic (1.80), and cholinergic (1.76) systems were positively enriched, whereas purinergic/adenosine was negatively enriched (NES = -1.47, FDR = 0.0082). Non-overlapping transmitter modules contained 138 glutamatergic, 69 GABAergic, 46 purinergic/adenosine, 32 dopaminergic, 27 cholinergic, 23 adrenergic, and 14 serotonergic genes (Supplementary Tables S11-S15).

(E) Naive-centered temporal changes across seven non-overlapping neurotransmitter-associated modules. Heatmap values represent mean module-score differences relative to Naive for GABAergic, adrenergic, glutamatergic, dopaminergic, cholinergic, serotonergic, and purinergic/adenosine systems across CCI 24 h, 7 d, and 6 mo.

(F) Targeted synaptic-identity and dopamine-recipient gene expression across the same CCI time course. Heatmap values represent group mean gene-wise z-score changes relative to Naive. Synaptic genes showed predominantly delayed increases, while dopamine-recipient genes were heterogeneous, including a divergent Ppp1r1b profile.

#### Supplementary Figure 3. Metric robustness and external context of delayed AQP4/SNTA1 expression imbalance

**A**

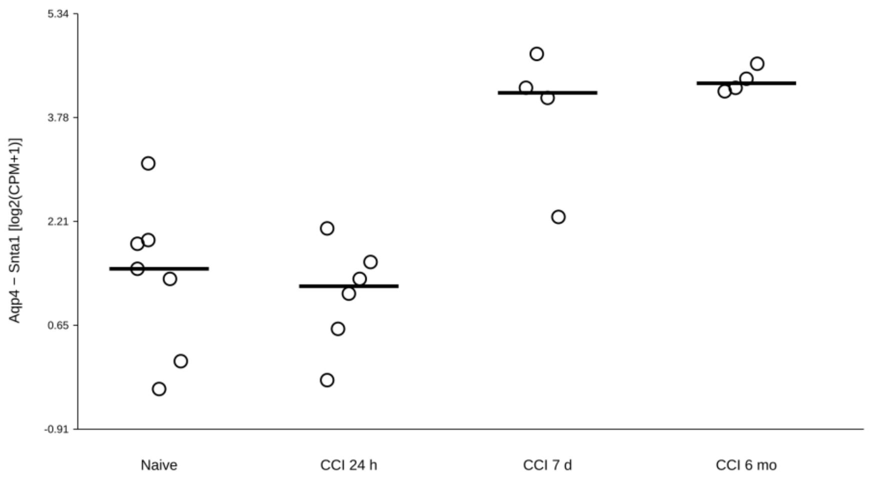


**B**

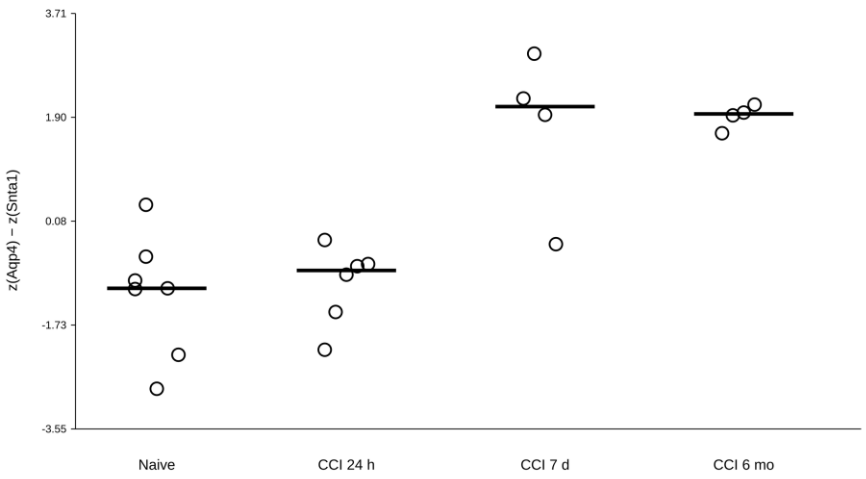

C

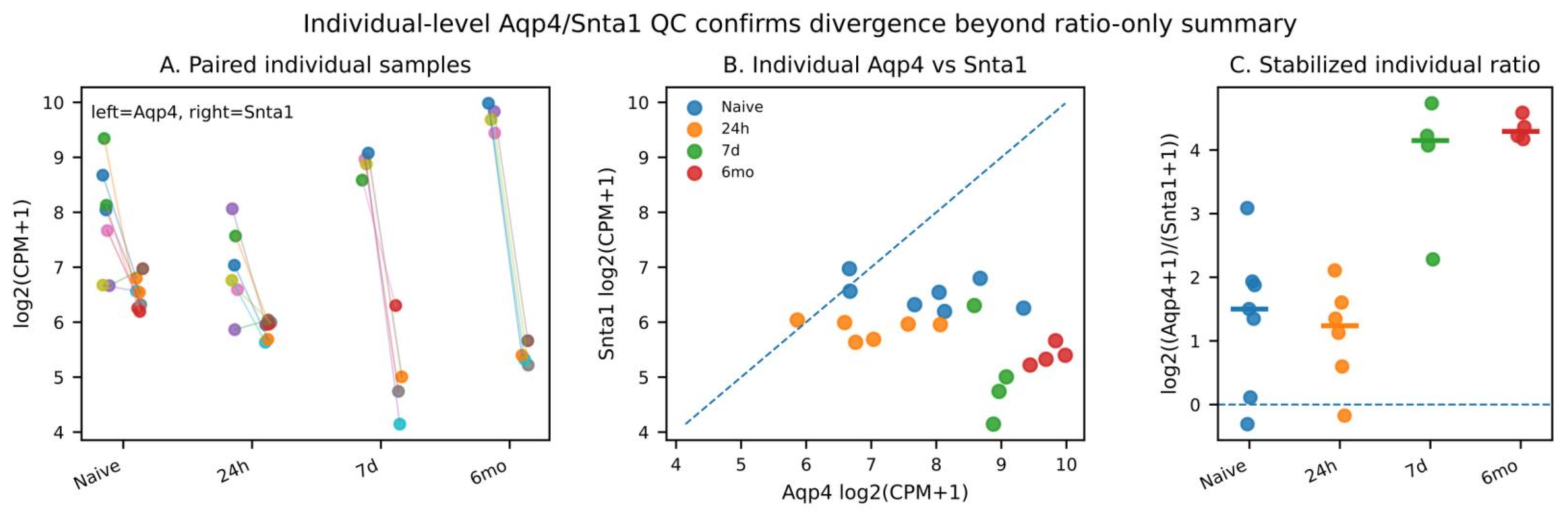

Individual-level Aqp4/Snta1 QC confirms divergence beyond ratio-only summary
A. Paired individual samples
left=Aqp4, right=Snta1
log2(CPM+1)
Naive
24h
7d
6mo
B. Individual Aqp4 vs Snta1
Naive
24h
7d
6mo
Snta1 log2(CPM+1)
Aqp4 log2(CPM+1)
C. Stabilized individual ratio
log2((Aqp4+1)/(Snta1+1))
Naive
24h
7d
6mo


D

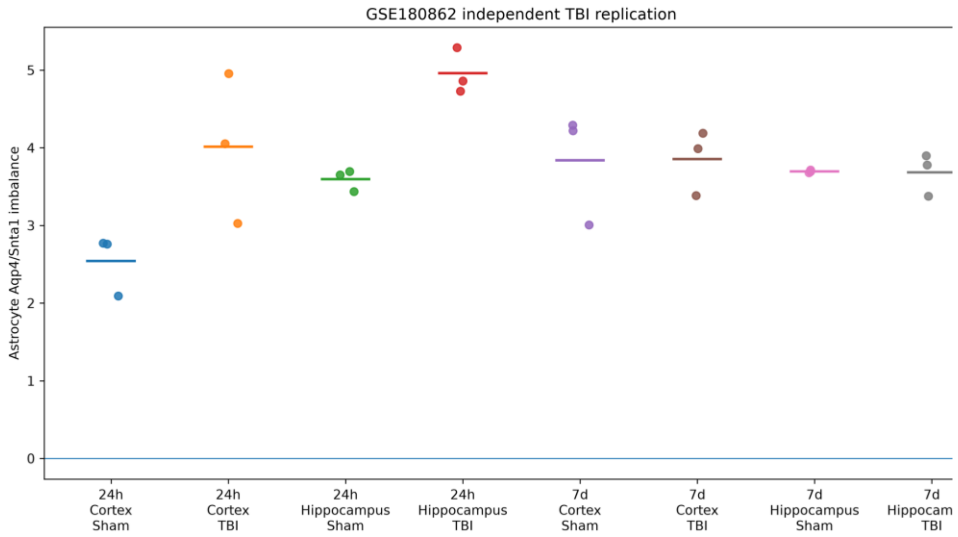

GSE180862 independent TBI replication
Astrocyte Aqp4/Snta1 imbalance
24h Cortex Sham
24h Cortex TBI
24h Hippocampus Sham
24h Hippocampus TBI
7d Cortex Sham
7d Cortex TBI
7d Hippocampus Sham
7d Hippocam TBI


E

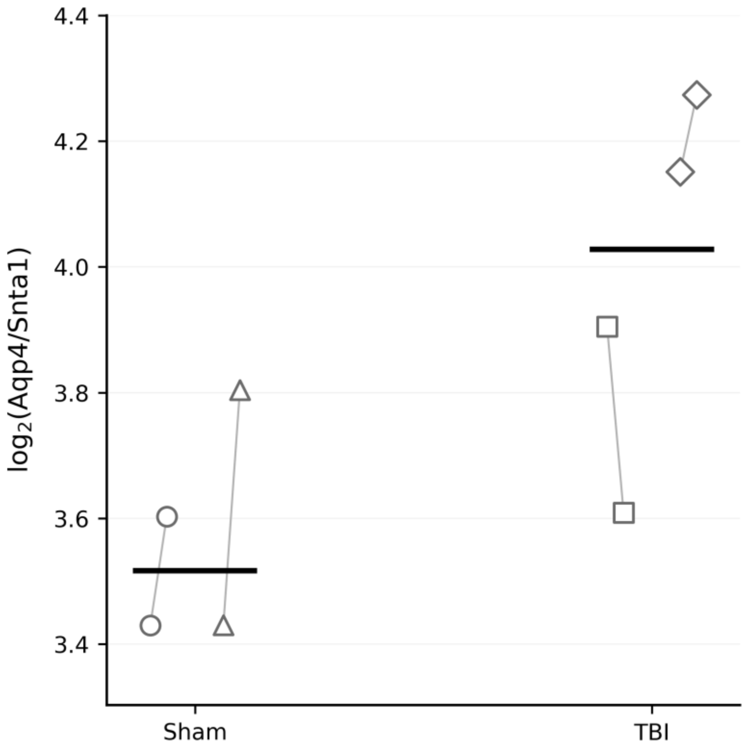

$log_2$(Aqp4/Snta1)
Sham
TBI

#### Legend

(A,B) Alternative definitions of the AQP4/SNTA1 transcriptional relationship reproduced the delayed separation seen with the primary score. Panel A uses the direct difference between Aqp4 and Snta1 on the log2(CPM + 1) scale; panel B uses the difference between standardized Aqp4 and Snta1 expression. These sensitivity formulations were evaluated to ensure that the delayed pattern did not depend on one pseudocount or scaling convention (Supplementary Table S23).

(C) Individual-sample QC demonstrates that the delayed signal reflects the underlying Aqp4 and Snta1 measurements rather than a ratio-only artifact. In the primary analysis, Aqp4 showed its clearest increase at 6 mo (+1.85 log2[CPM+1] versus Naive; exact $p = 0.0121$, FDR = 0.0182), whereas Snta1 was reduced at 24 h (-0.64, exact $p = 0.00117$, FDR = 0.0091), 7 d (-1.47, $p = 0.00909$, FDR = 0.0164), and 6 mo (-1.12, $p = 0.00303$, FDR = 0.0091). The Naive-referenced imbalance was unsupported at 24 h (-0.26, $p = 0.646$), but increased at 7 d (+2.46, 95% CI 1.20 to 3.60, $p = 0.00909$, FDR = 0.0164) and 6 mo (+2.97, 95% CI 2.19 to 3.78, $p = 0.00303$, FDR = 0.0091). Neither Aqp4 nor Snta1 was included in the positive astrocyte-like gating panel; thus, the phenotype was not generated by direct selection on either transcript (Supplementary Tables S20-S23).

The Naive reference regression showed essentially no baseline linear relationship between the two transcripts ($r \approx 0.045$, $R^2 \approx 0.002$, $p \approx 0.923$). Nevertheless, absolute deviation from the fixed Naive reference increased after injury, with median absolute residuals of approximately 0.23 in Naive, 0.73 at 24 h, 1.49 at 7 d, and 0.89 at 6 mo. These residuals quantify expression-level departure only and do not imply physical molecular coupling or uncoupling (Supplementary Table S24).

(D) GSE180862 independent mouse TBI context. The prespecified 7-d comparison did not reproduce the GSE269748 discovery direction after accounting for brain region; secondary 24-h and region-specific contrasts provided only limited directional context. This negative result is retained as an external-replication boundary rather than excluded from the evidence set (Supplementary Table S25).

(E) GSE282909 spatial-transcriptomic context. Astrocyte-enriched mouse-level summaries were numerically higher for log2(Aqp4/Snta1) in TBI than Sham animals (approximately 3.98 versus 3.57), but only two mice were available per condition; the result is therefore interpreted as trend-level spatial context rather than inferential replication (Supplementary Table S26). Chronic human context in the main Figure 3 was also directionally supportive but non-definitive: GSE155114 showed a numerically higher donor-level AQP4/SNTA1 contrast in CTE than Control (n = 8/group), and PXD007694 showed a numerically higher protein contrast in CTE but no significant group difference (Control n = 6, CTE n = 11; median difference = 0.41; Mann-Whitney $p = 0.350$; Supplementary Table S27).

#### Supplementary Figure 4. Human contextual evidence for heterogeneous endfoot and homeostatic remodeling

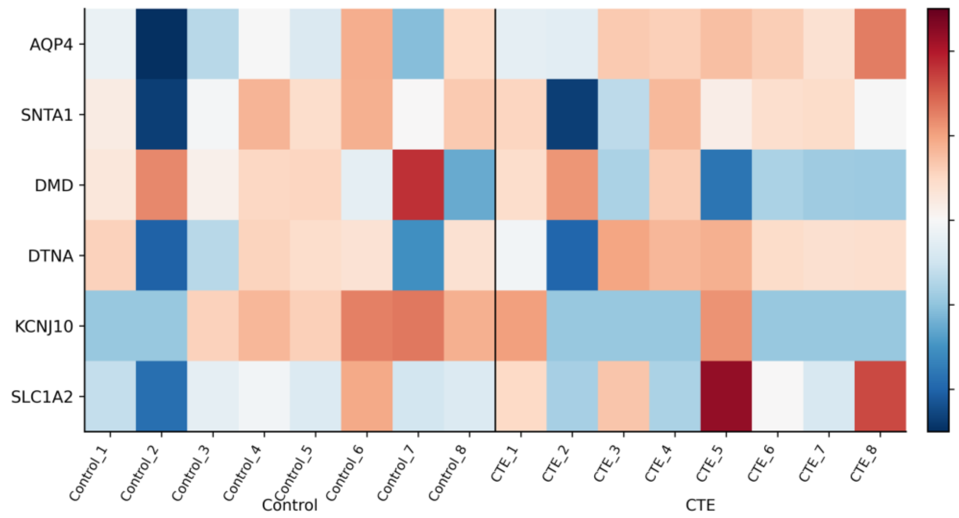


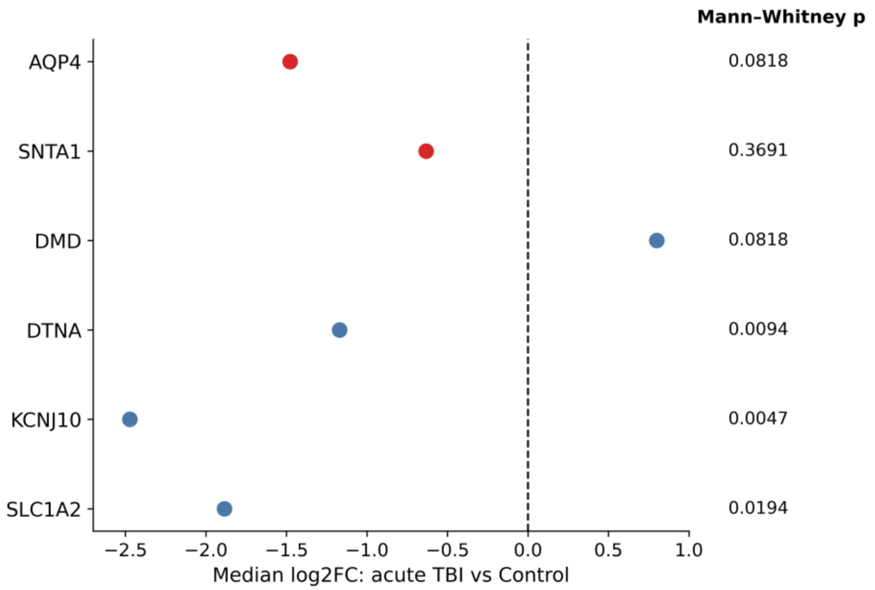

**C**

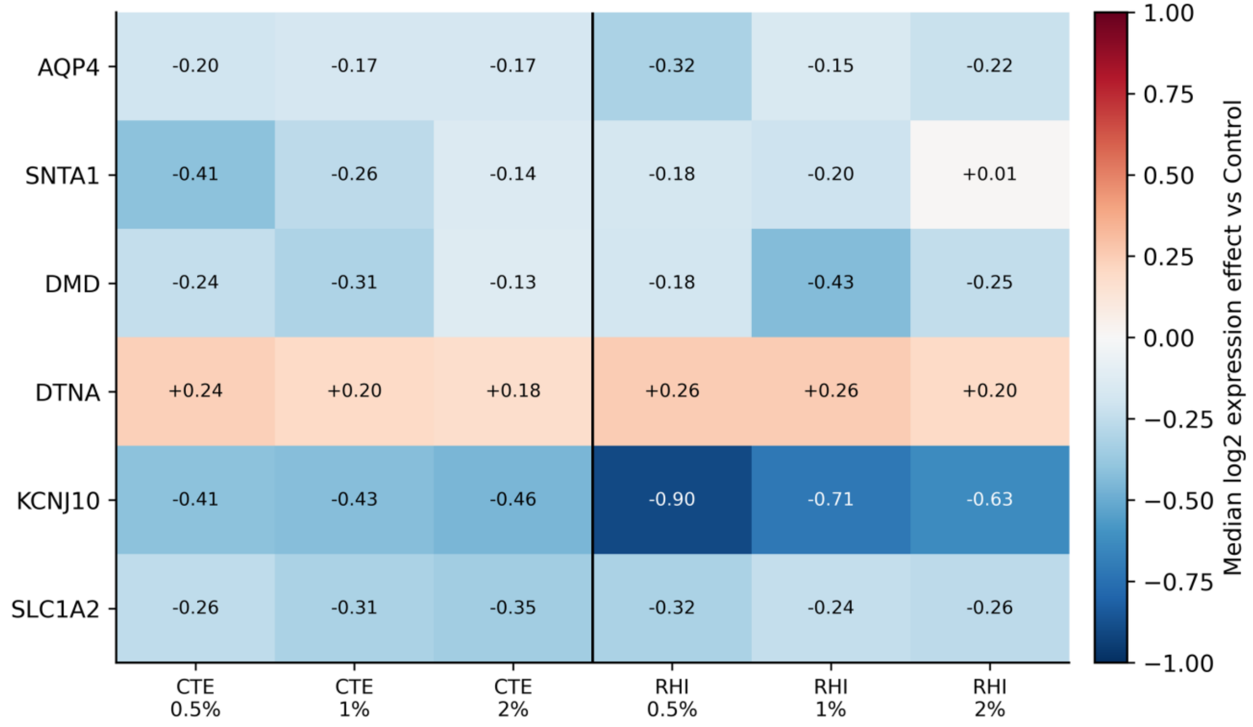


#### Legend

(A) Chronic human CTE transcriptomic context from GSE155114. Donor-level selected-gene expression is shown for AQP4, SNTA1, DMD, DTNA, KCNJ10, and SLC1A2 in Control and CTE donors. The donor-level patterns are heterogeneous and are presented as chronic human context for the broader endfoot/homeostatic domain, not as replication of the mouse CCI state or as evidence of physical AQP4/SNTA1 uncoupling (Supplementary Tables S28-S30).

(B) Acute severe human TBI selected-gene effects from GSE209552. Median TBI-versus-Control log2 expression effects were AQP4 -1.48, SNTA1 -0.63, DMD +0.80, DTNA -1.17, KCNJ10 -2.47, and SLC1A2 -1.88. Mann-Whitney p values shown in the panel were 0.0818, 0.3691, 0.0818, 0.0094, 0.0047, and 0.0194, respectively. The acute human direction differs from the delayed positive homeostatic remodeling in mouse CCI and therefore emphasizes stage, species, tissue, and compartment dependence (Supplementary Tables S31-S32).

(C) GSE261807 repetitive-head-impact/early-CTE threshold sensitivity. Median gene-level effects are shown across the prespecified top 0.5%, 1%, and 2% marker-defined astrocyte-selection thresholds for CTE and RHI comparisons. AQP4, SNTA1, DMD, KCNJ10, and SLC1A2 were generally negative across thresholds, whereas DTNA remained positive. The magnitude of several effects varied with the retained-cell threshold, so this dataset is explicitly treated as threshold-sensitive contextual evidence rather than robust validation (Supplementary Tables S33-S34).

Related primary GSE269748 context retained in Supplementary Tables S28-S36: at 24 h, selected endfoot/homeostatic changes included Aqp4 +0.35, Snta1 -0.47, Kcnj10 -0.34, Gjb6 -0.61, Slc1a2 -1.03, Slc1a3 -1.22, and Glul -0.69 relative to Naive. At 7 d, Aqp4 +1.92, Snta1 -1.73, Dmd +1.07, Dtna +0.72, Kcnj10 +1.81, Gja1 +1.68, and Slc1a2 +0.95 illustrated heterogeneous reorganization. At 6 mo, Aqp4 +2.72, Dmd +2.22, Dtna +1.65, Kcnj10 +2.73, Gja1 +2.76, Gjb6 +2.60, and Slc1a2 +2.77 were increased while Snta1 remained below Naive (-1.04). These contextual datasets collectively support heterogeneous perturbation of the endfoot/homeostatic domain, not universal replication, AQP4 polarization changes, endfoot failure, or causality.

**Supplementary Figure 5. Pathway specificity, redundancy, regulatory context, and robustness of the potassium and ion homeostasis association**

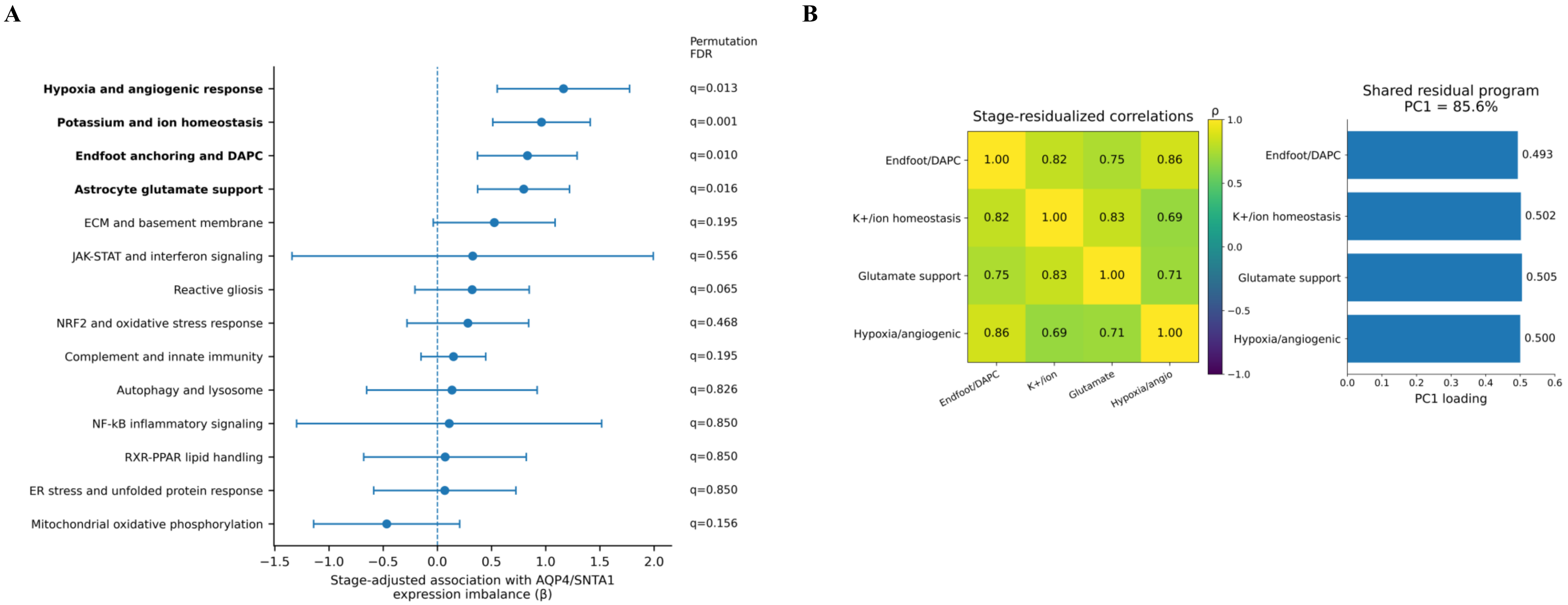

C

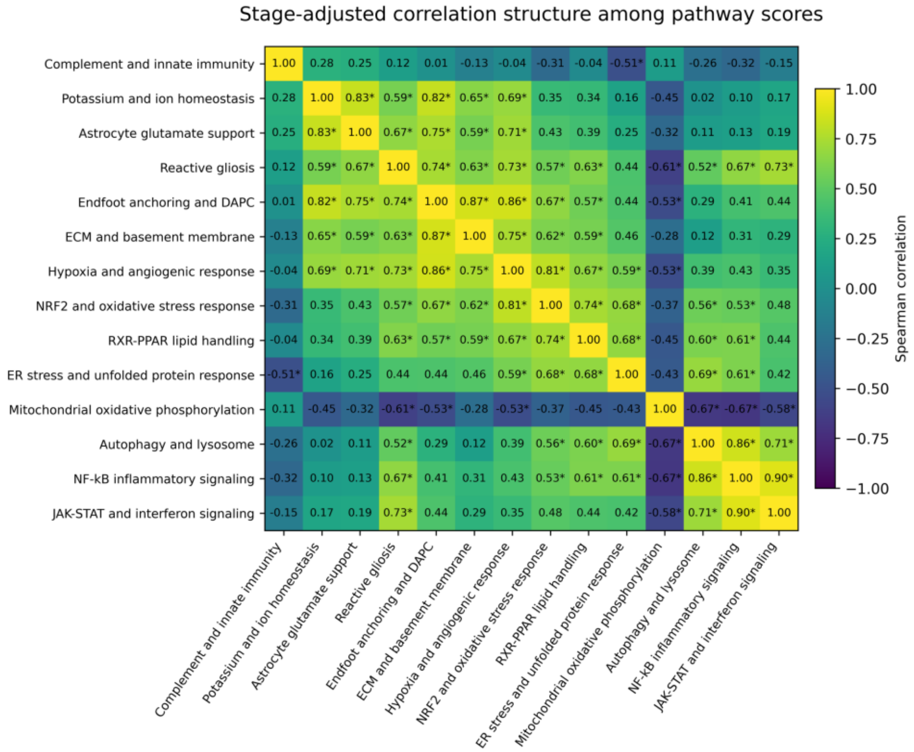
Stage-adjusted correlation structure among pathway scores
Complement and innate immunity
Potassium and ion homeostasis
Astrocyte glutamate support
Reactive gliosis
Endfoot anchoring and DAPC
ECM and basement membrane
Hypoxia and angiogenic response
NRF2 and oxidative stress response
RXR-PPAR lipid handling
ER stress and unfolded protein response
Mitochondrial oxidative phosphorylation
Autophagy and lysosome
NF-kB inflammatory signaling
JAK-STAT and interferon signaling
Spearman correlation
1.00
0.75
0.50
0.25
0.00
−0.25
−0.50
−0.75
−1.00

D

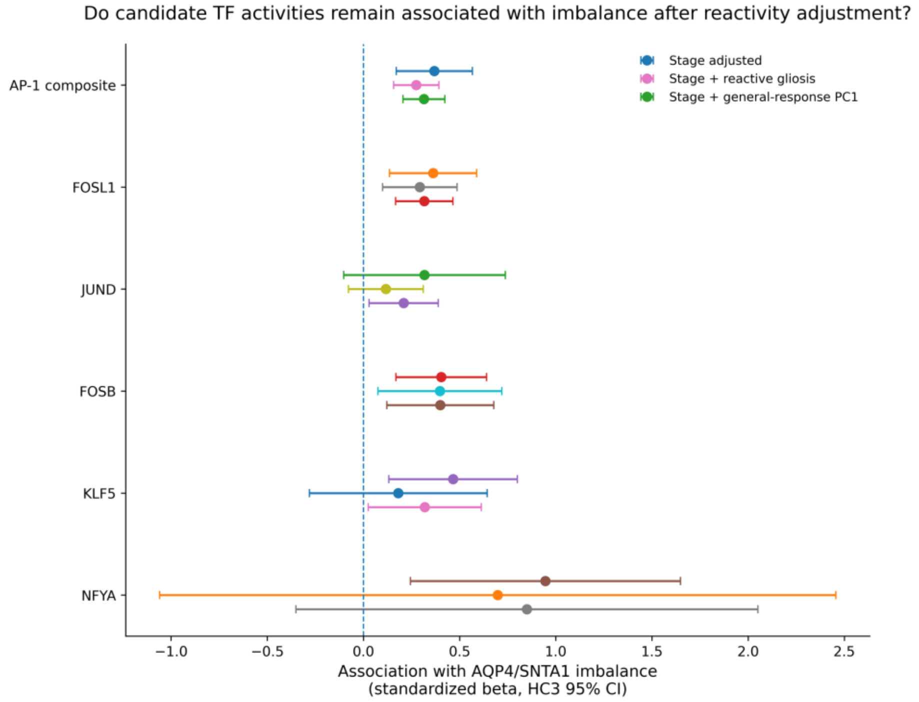
Do candidate TF activities remain associated with imbalance after reactivity adjustment?
Stage adjusted
Stage + reactive gliosis
Stage + general-response PC1
AP-1 composite
FOSL1
JUND
FOSB
KLF5
NFYA
−1.0
−0.5
0.0
0.5
1.0
1.5
2.0
2.5
Association with AQP4/SNTA1 imbalance
(standardized beta, HC3 95% CI)

**E**

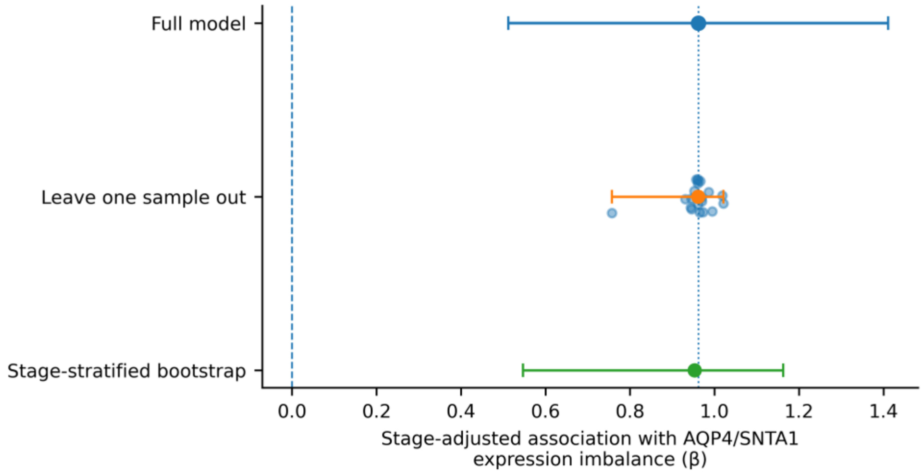

Full model
Leave one sample out
Stage-stratified bootstrap
0.0
0.2
0.4
0.6
0.8
1.0
1.2
1.4
Stage-adjusted association with AQP4/SNTA1
expression imbalance (β)


**F**

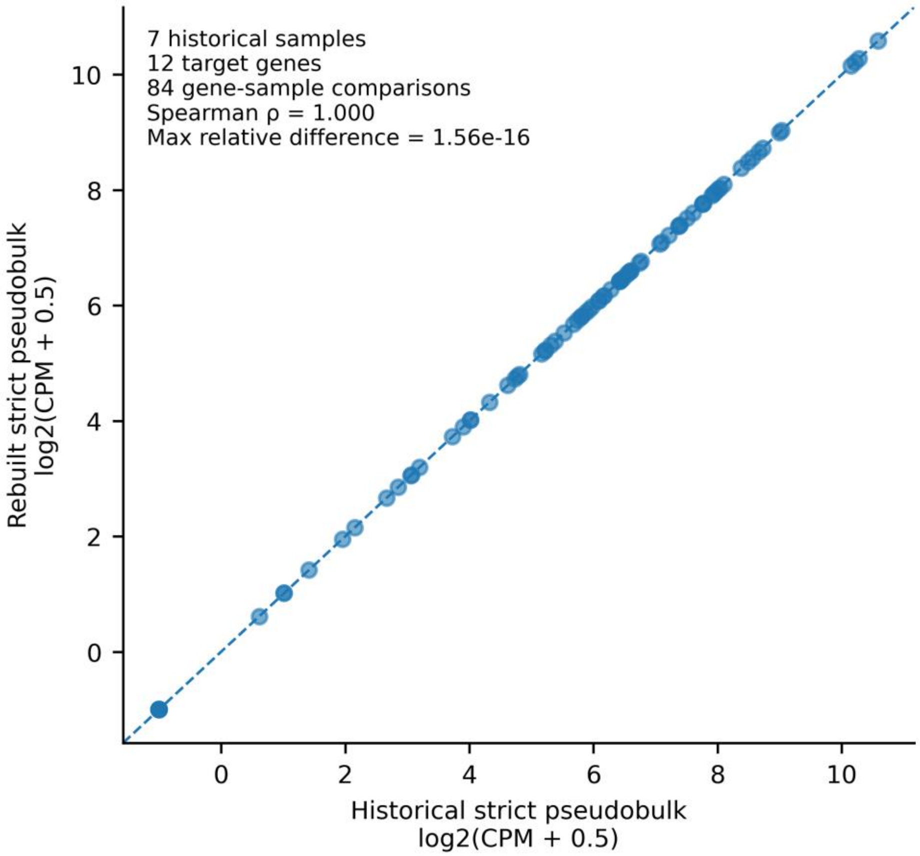

7 historical samples
12 target genes
84 gene-sample comparisons
Spearman ρ = 1.000
Max relative difference = 1.56e-16
Rebuilt strict pseudobulk
log2(CPM + 0.5)
Historical strict pseudobulk
log2(CPM + 0.5)
0
2
4
6
8
10

**G**

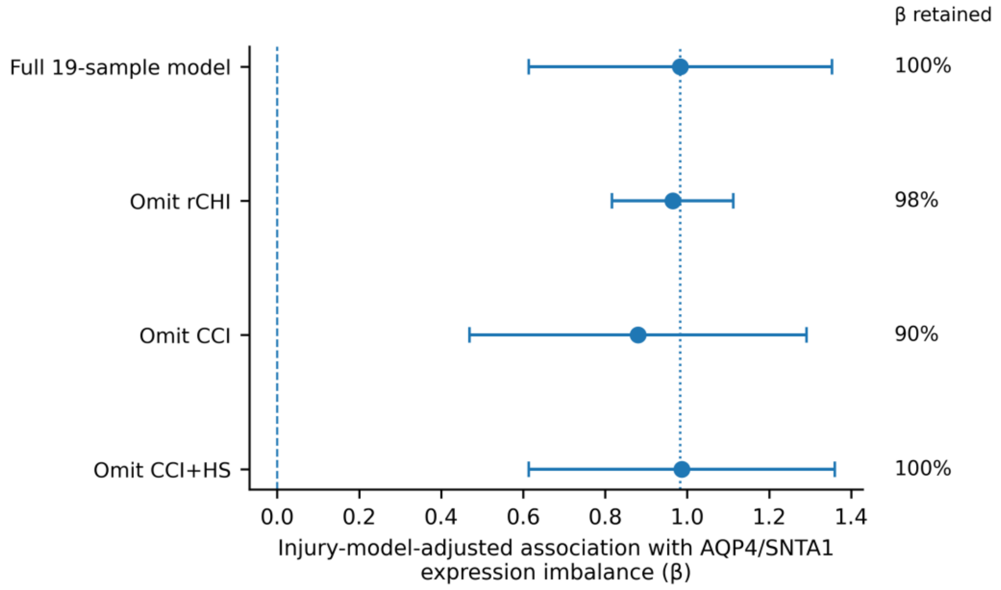
β retained
Full 19-sample model
Omit rCHI
Omit CCI
Omit CCI+HS
100%
98%
90%
100%
0.0
0.2
0.4
0.6
0.8
1.0
1.2
1.4
Injury-model-adjusted association with AQP4/SNTA1 expression imbalance (β)

**H**

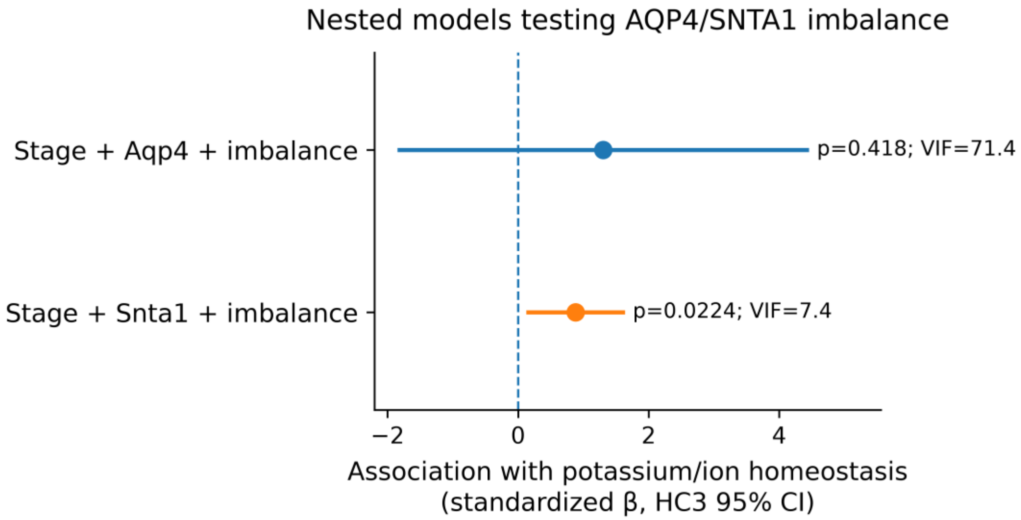


Interpretation: imbalance remains associated after Snta1 adjustment, but the Aqp4-adjusted model shows severe collinearity and should not be interpreted as evidence against an imbalance contribution.

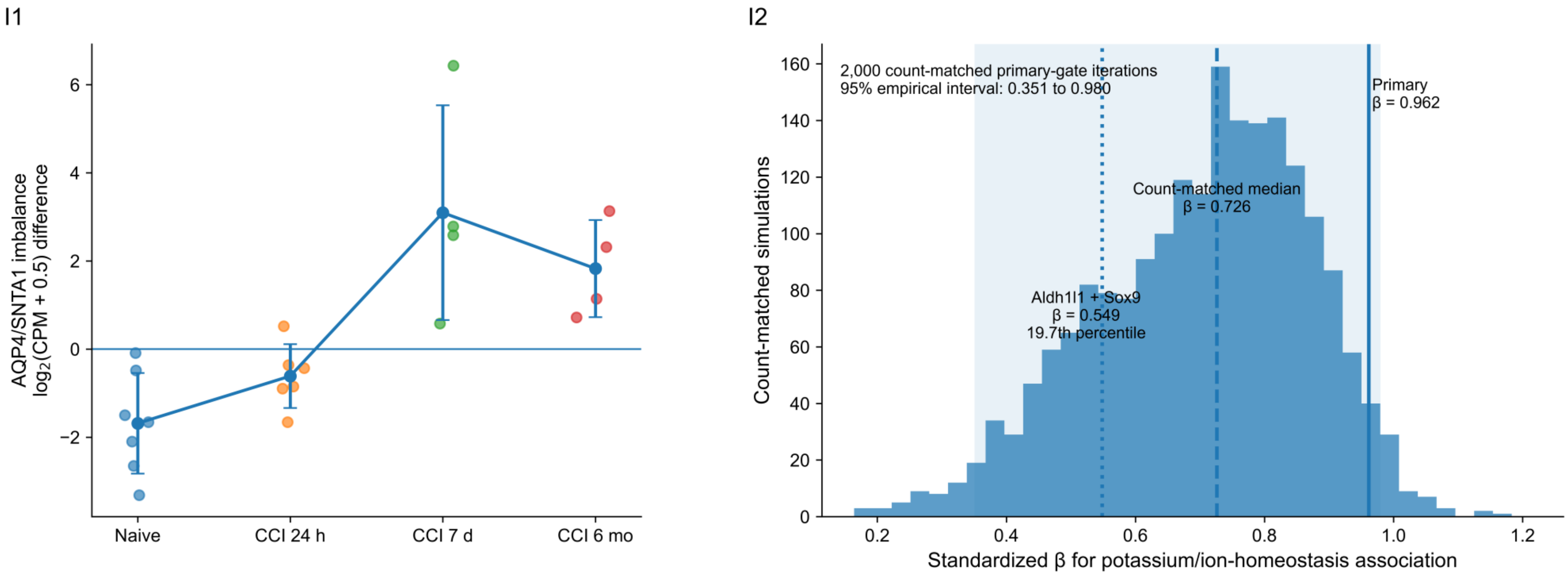


**Legend**

(A) Complete prespecified 14-pathway stage-adjusted screen. Four programs survived permutation-based FDR correction: hypoxia/angiogenic response (β = 1.164, q = 0.013), potassium/ion homeostasis (β = 0.962, q ≈ 0.0014), endfoot anchoring/DAPC (β = 0.831, q = 0.010), and astrocyte glutamate support (β = 0.797, q = 0.016). Reactive gliosis did not survive correction (q = 0.065); ECM/basement membrane and complement/innate immunity each had q = 0.195, NRF2/oxidative stress q = 0.468, JAK-STAT/interferon q = 0.556, autophagy/lysosome q = 0.826, and NF-κB, RXR-PPAR lipid handling, and ER-stress/UPR pathways were unsupported at q ≈ 0.850. Mitochondrial oxidative phosphorylation was also unsupported (q = 0.156). This complete screen shows that potassium/ion homeostasis was prioritized from a prespecified pathway family rather than selected post hoc (Supplementary Table S38).

(B) Residual redundancy among the four supported core programs after adjustment for injury stage. Pairwise Spearman correlations were endfoot/DAPC versus potassium/ion homeostasis ρ = 0.82, endfoot/DAPC versus glutamate support ρ = 0.75, endfoot/DAPC versus hypoxia/angiogenic ρ = 0.86, potassium/ion homeostasis versus glutamate support ρ = 0.83, potassium/ion homeostasis versus hypoxia/angiogenic ρ = 0.69, and glutamate support versus hypoxia/angiogenic ρ = 0.71. A shared residual PC1 explained 85.6% of variance with near-balanced positive loadings: endfoot/DAPC 0.493, potassium/ion homeostasis 0.502, glutamate support 0.505, and hypoxia/angiogenic response 0.500. Thus, the four signals share substantial transcriptional structure but are not assumed to be biologically identical (Supplementary Tables S39-S40).

(C) Full stage-residual correlation structure across all 14 prespecified pathway scores. This panel places the four supported pathways within the broader covariance structure and documents the extent to which inflammatory, homeostatic, metabolic, and stress-response scores share residual variation after accounting for stage. The heatmap is descriptive correlation structure and is not a causal network (Supplementary Table S39).

(D) Candidate transcription-factor specificity after adjustment for broader reactivity. AP-1 composite activity and selected candidates including FOSL1, JUND, FOSB, KLF5, and NFYA were evaluated using inferred CollecTRI/decoupler activities. The potassium association remained supported after adjustment for AP-1 activity, including models additionally adjusted for reactive gliosis or broad-response PC1 (permutation q = 0.011, 0.014, and 0.011). In reciprocal models, AP-1 was no longer independently supported after potassium adjustment (q = 0.455, 0.455, and 0.440). These results indicate comparative statistical specificity, not mediation, causal ordering, direct TF binding, or direct transcriptional regulation (Supplementary Table S42).

(E) Sample-resampling stability of the locked 10-gene potassium/ion-homeostasis association. The reference stage-adjusted model yielded β = 0.962, partial $R^2 = 0.775$, and permutation p = 0.0002. All 21 leave-one-biological-sample-out estimates remained positive (median β = 0.961; range approximately 0.757 to 1.021; minimum retained β fraction ≈ 0.787). Across 5,000 stage-stratified bootstrap resamples, the median β was 0.953 (approximate interval 0.547 to 1.163) and 99.98% of estimates were positive. These are internal stability analyses, not independent replication (Supplementary Table S43).

(F) Historical strict-gate computational reproducibility. The rebuilt workflow included 7 historical samples and 12 target genes, giving 84 gene-sample comparisons. Recomputed pseudobulk values reproduced the archived values to numerical precision (Spearman ρ = 1.000; maximum relative difference = $1.56 \times 10^{-16}$). This establishes reproducibility of the archived computation, not purified cell identity or biological validation of the astrocyte gate (Supplementary Table S44).

(G) Matched 24-h cross-injury-model sensitivity in the strict-gate subset (n = 19: Naive n = 7, rCHI n = 3, CCI n = 6, CCI+HS n = 3). The injury-model-adjusted association remained positive (β = 0.983, 95% CI 0.613 to 1.353; permutation p = 0.00030), with no supported model-by-imbalance interaction (p = 0.628). Leave-one-model-out estimates remained positive after omitting rCHI (β = 0.965; 98% retained), CCI (β = 0.880; 90% retained), or CCI+HS (β = 0.987; approximately 100% retained). Given the small model-specific groups, this supports within-GSE269748 robustness rather than injury-model equivalence or independent replication (Supplementary Table S46).

Additional gene and gene-set robustness corresponding to main Figure 5D-E is retained in Supplementary Tables S43 and S45. The locked 10-gene program comprised Kcnj10, Kcnj16, Atp1a2, Atp1b2, Slc12a2, Slc8a1, Clcn2, Trpv4, Gja1, and Gjb6. Positive distributed support was observed for Kcnj10, Kcnj16, Atp1a2, Atp1b2, Clcn2, Gja1, and Gjb6; Slc8a1 was essentially null, Slc12a2 was negative/unsupported, and Trpv4 showed a negative permutation-supported direction (q ≈ 0.002) with an HC3 interval crossing zero. All 10 single-gene omissions remained positive. Grouped omission coefficients remained positive after removal of Kcnj10 (β ≈ 0.935), connexins Gja1/Gjb6 (β ≈ 0.856), Na+/K+-ATPase components Atp1a2/Atp1b2 (β ≈ 0.840), or inward-rectifier channels Kcnj10/Kcnj16 (β ≈ 0.913). These analyses establish internal robustness and statistical specificity only; they do not demonstrate potassium flux, impaired K+ clearance, AQP4 polarization, mediation, or causality.

(H) Nested stage-adjusted models examining the AQP4/SNTA1 imbalance relative to its component transcripts. When the imbalance and Snta1 were included together, the imbalance remained associated with potassium/ion-homeostasis (standardized β = 0.881, HC3 95% CI 0.125 to 1.637, p = 0.022), whereas Snta1 was unsupported (β = -0.151, p = 0.823). In the corresponding model containing Aqp4 and the imbalance, neither coefficient was independently supported. This model showed severe multicollinearity between Aqp4 and the mathematically related imbalance term (VIF = 52.2 for Aqp4 and 71.4 for imbalance), preventing clean attribution of independent effects. These nested models therefore support information in the relative Aqp4-Snta1 relationship beyond Snta1 alone but do not establish independence from Aqp4.

(I1) Program-independent astrocyte-gate sensitivity of AQP4/SNTA1 imbalance. Astrocyte-like pseudobulk was reconstructed using only Aldh1l1 and Sox9, thereby excluding genes contained in the tested homeostatic programs from cell selection. Points represent individual samples and connected symbols show stage means ± SD. The delayed increase in Aqp4/Snta1 imbalance was preserved, with the largest shift at 7 d and persistence at 6 mo (Kruskal-Wallis p = 0.0019).

(I2) Count-matched sensitivity of the potassium/ion-homeostasis association. The original astrocyte gate was randomly downsampled 2,000 times to match the per-sample cell numbers retained by the Aldh1l1 + Sox9 gate. The histogram shows the distribution of stage-adjusted standardized β values for the potassium/ion-homeostasis association with AQP4/SNTA1 imbalance. The shaded region denotes the 95% empirical interval (0.351 to 0.980), the dashed line marks the count-matched median β = 0.726, the solid line marks the full primary-gate β = 0.962, and the dotted line marks the program-independent-gate β = 0.549, corresponding to the 19.7th percentile of the count-matched distribution.

#### Supplementary Figure 6. Human neuropathology, independent chronic-mouse, and temporal evidence boundaries

**A**

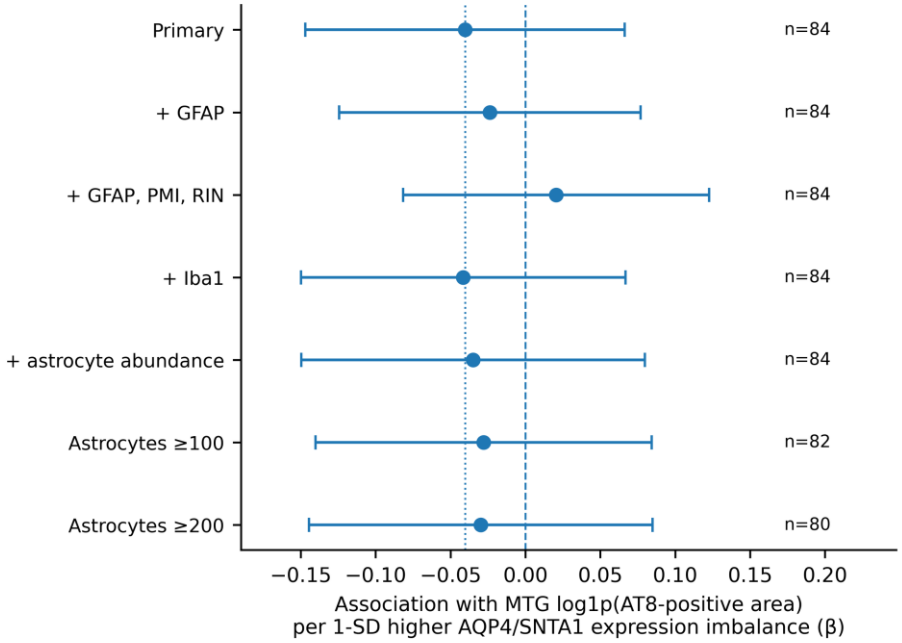

**B**

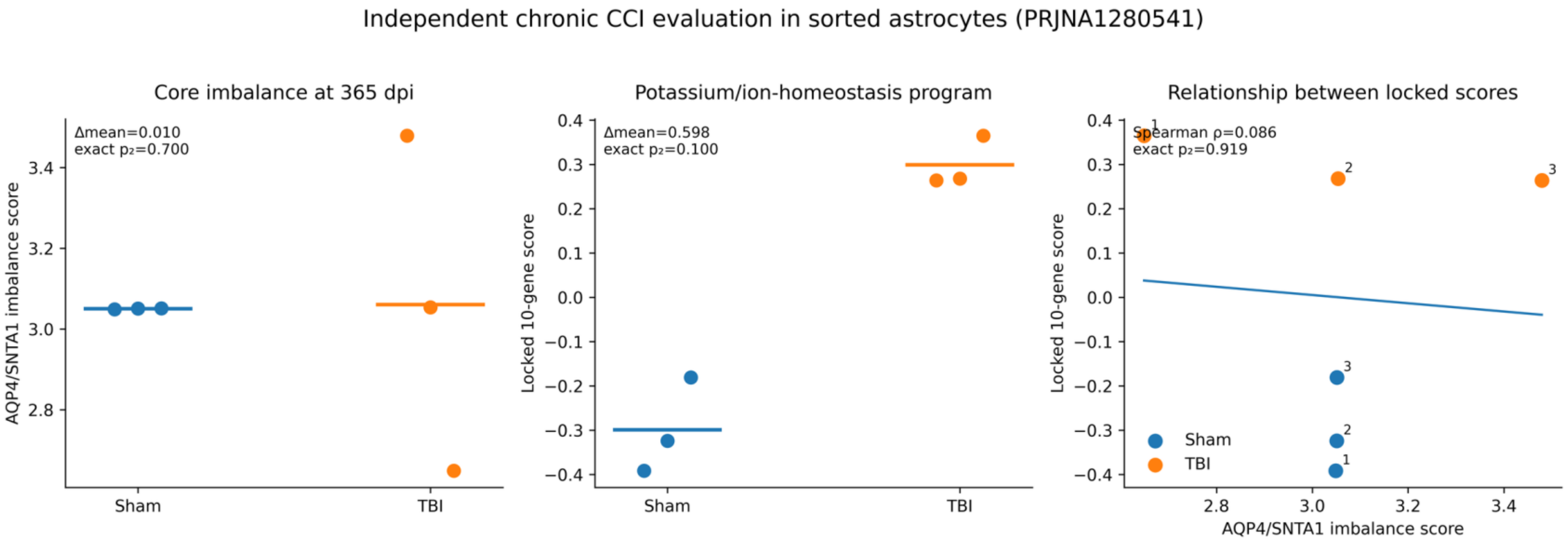
Independent chronic CCI evaluation in sorted astrocytes (PRJNA1280541)
Core imbalance at 365 dpi
Δmean=0.010
exact p₂=0.700
AQP4/SNTA1 imbalance score
Sham
TBI
Potassium/ion-homeostasis program
Δmean=0.598
exact p₂=0.100
Locked 10-gene score
Relationship between locked scores
Spearman ρ=0.086
exact p₂=0.919
AQP4/SNTA1 imbalance score

**C**

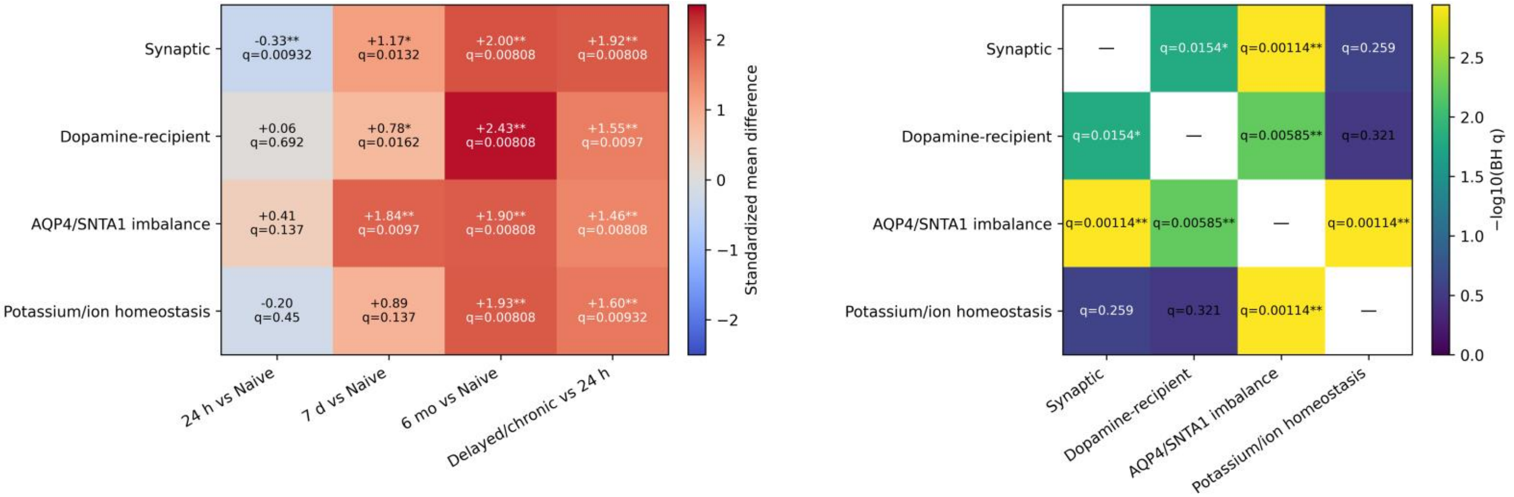
Synaptic
Dopamine-recipient
AQP4/SNTA1 imbalance
Potassium/ion homeostasis
-0.33** q=0.00932
+1.17* q=0.0132
+2.00** q=0.00808
+1.92** q=0.00808
+0.06 q=0.692
+0.78* q=0.0162
+2.43** q=0.00808
+1.55** q=0.0097
+0.41 q=0.137
+1.84** q=0.0097
+1.90** q=0.00808
+1.46** q=0.00808
-0.20 q=0.45
+0.89 q=0.137
+1.93** q=0.00808
+1.60** q=0.00932
24 h vs Naive
7 d vs Naive
6 mo vs Naive
Delayed/chronic vs 24 h
Standardized mean difference
q=0.0154*
q=0.00114**
q=0.259
q=0.00585**
q=0.321
−log10(BH q)

**D**

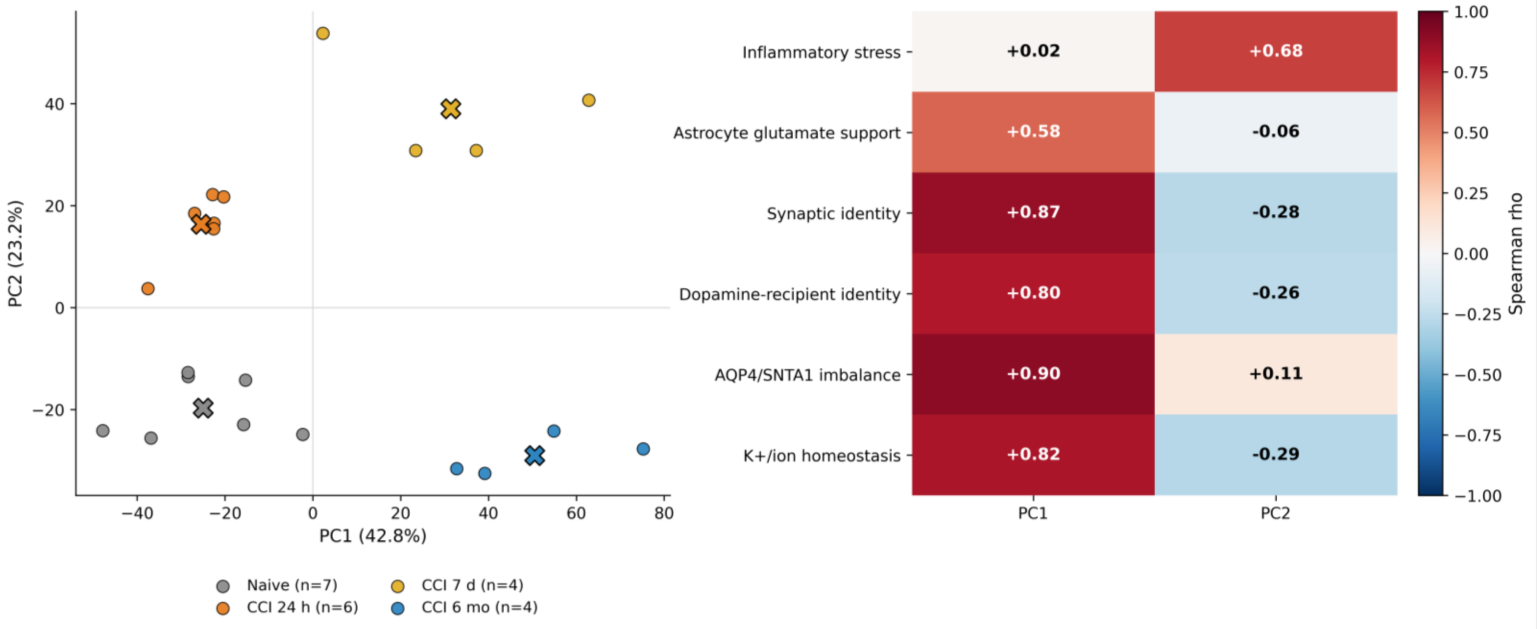


#### Legend

(A) Prespecified SEA-AD middle temporal gyrus sensitivity analyses. The association between astrocyte AQP4/SNTA1 expression imbalance and log1p-transformed AT8-positive tau area remained close to zero after additional adjustment for GFAP, combined GFAP/PMI/RIN, Iba1, or astrocyte abundance and after restricting the analysis to donors with at least 100 or 200 retained astrocytes. Confidence intervals crossed zero across all models, providing no evidence that greater astrocyte AQP4/SNTA1 imbalance predicts greater human tau burden (Supplementary Table S53).

(B) Independent chronic mouse evaluation in sorted astrocytes from PRJNA1280541 at 365 days after TBI (Sham, n = 3; TBI, n = 3). The AQP4/SNTA1 imbalance did not replicate (mean TBI-Sham difference = 0.010; exact two-sided p = 0.700). The locked potassium and ion homeostasis score showed a positive chronic shift (+0.598; exact two-sided p = 0.100), whereas the relationship between the imbalance and potassium scores was weak (Spearman ρ = 0.086; exact p = 0.919). These data provide limited directional support for chronic ion-homeostatic remodeling but not replication of the AQP4/SNTA1 imbalance or its potassium association (Supplementary Table S54).

(C) Detailed temporal comparison of synaptic identity, dopamine-recipient identity, AQP4/SNTA1 imbalance, and potassium and ion homeostasis across the GSE269748 CCI time course. Left, standardized mean differences for 24 h versus Naive, 7 d versus Naive, 6 mo versus Naive, and delayed or chronic stages versus 24 h. Right, pairwise comparison of temporal profiles using globally adjusted q values. These analyses demonstrate partially distinct temporal trajectories among the four transcriptional dimensions.

(D) Relationship between predefined transcriptional responses and the unsupervised whole-transcriptome structure of GSE269748. Left, PCA of the 21 biological samples, with PC1 and PC2 explaining 42.8% and 23.2% of variance, respectively. Right, Spearman correlations between predefined transcriptional scores and PC1 or PC2. PC1 showed strongest correspondence with AQP4/SNTA1 imbalance, synaptic identity, potassium and ion homeostasis, and dopamine-recipient identity, whereas inflammatory stress aligned more strongly with PC2. These correlations arise from the same expression dataset and are interpreted as axis correspondence rather than independent validation

.**Supplementary Figure 7. Spatial organization of AQP4/SNTA1 and related homeostatic programs after mild TBI**

**A**

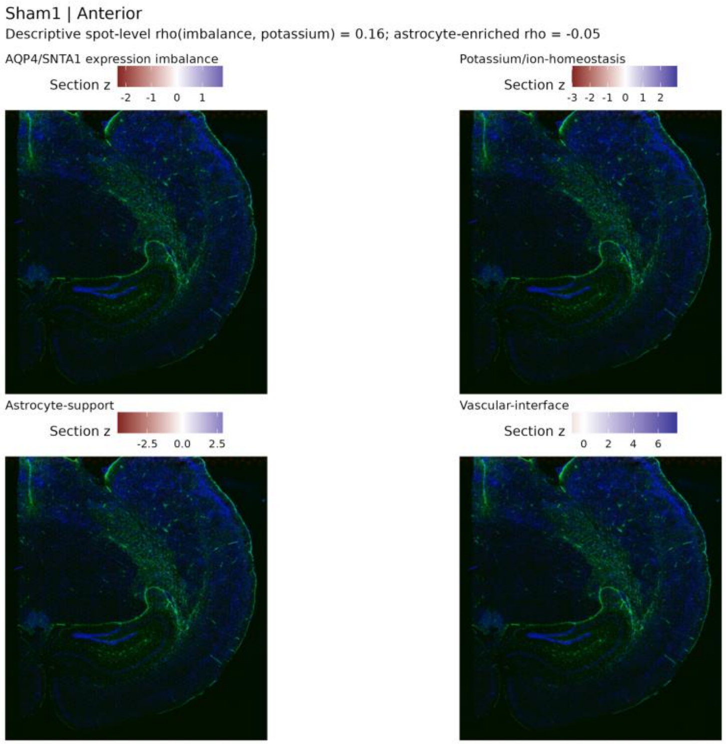


**B**

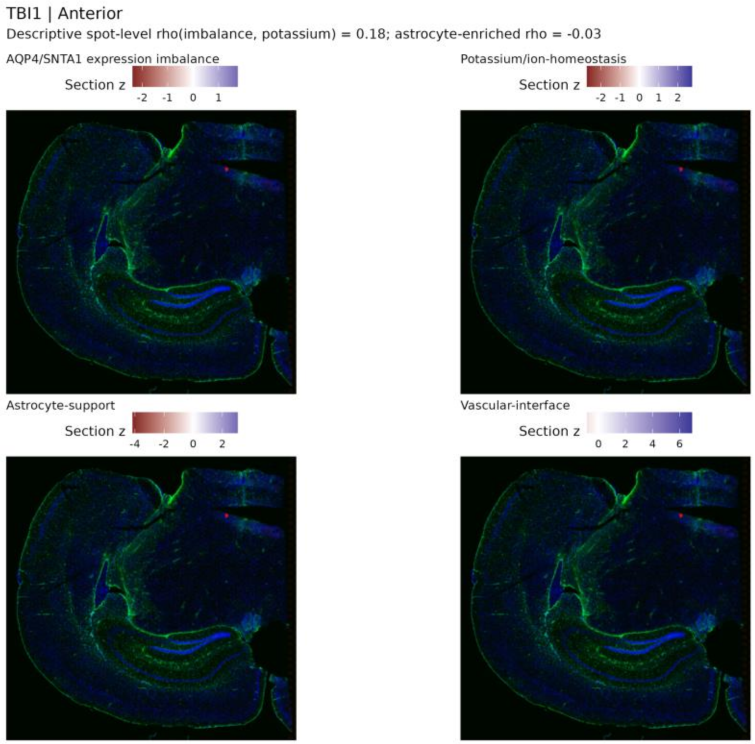

**C**

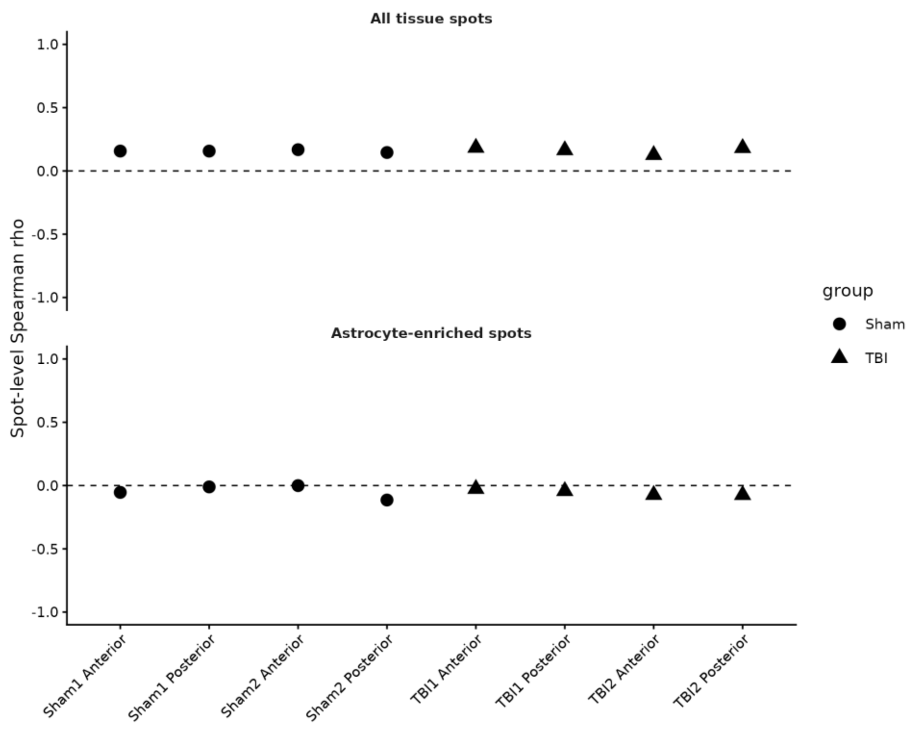


**D**

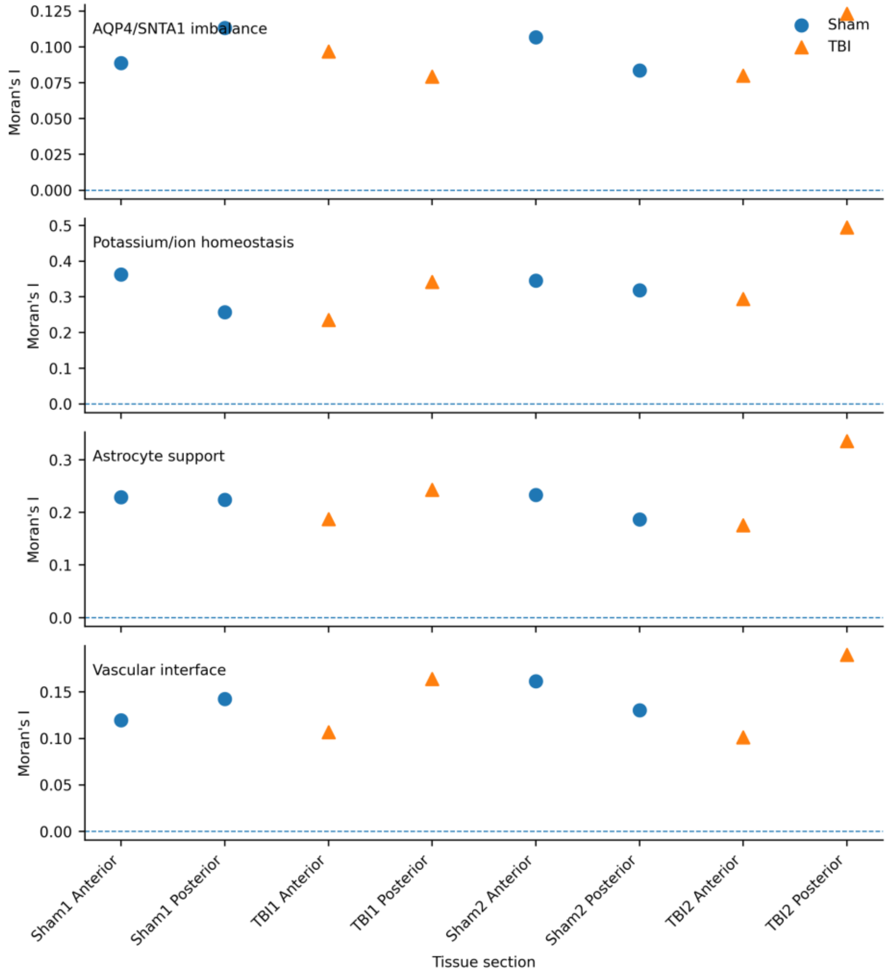


#### Legend

(A,B) Representative Sham and TBI Visium sections from GSE282909 showing within-section z-scored AQP4/SNTA1 expression imbalance, potassium/ion-homeostasis, astrocyte-support, and vascular-interface scores over the corresponding tissue image. Eight sections were analyzed in total: anterior and posterior sections from two Sham and two TBI mice. Within-section z scores are used for visualization and should not be interpreted as between-animal effect sizes. The displayed sections illustrate heterogeneous regional organization of each program (Supplementary Tables S55-S56).

(C) Section-level Spearman correspondence between AQP4/SNTA1 imbalance and potassium/ion-homeostasis scores. Across all tissue spots, correlations were modestly positive in each section. When restricted to astrocyte-enriched spots, defined as the upper quartile of the section-specific astrocyte-support score, the relationship approached zero and was slightly negative in several sections. Thus, the sample-level association observed in GSE269748 does not translate into strong local spot-level coupling in this spatial dataset (Supplementary Table S57).

(D) Global Moran's I for AQP4/SNTA1 imbalance, potassium/ion homeostasis, astrocyte support, and vascular-interface scores across all eight sections. All four programs showed positive within-section spatial autocorrelation, demonstrating non-random anatomical organization. The primary spatial graph used k = 6 nearest neighbors with 999 one-sided permutations; AQP4/SNTA1 spatial autocorrelation remained positive in sensitivity analyses using k = 4, 6, and 8. Because only two mice were available per condition, section-level autocorrelation is interpreted as within-section organization and not as animal-level evidence of a TBI-specific spatial effect (Supplementary Table S58).

The spatial analysis supports anatomical organization of the individual transcriptional programs, not protein colocalization, physical AQP4/SNTA1 uncoupling, perivascular AQP4 polarization, or functional endfoot impairment. Direct imbalance-potassium correspondence is weak at the spot level, especially in astrocyte-enriched spots, and the available n = 2 mice per group precludes strong condition-level inference.

#### Supplementary Figure 8. Additional cross-dataset contextual analyses for the multidimensional TBI framework

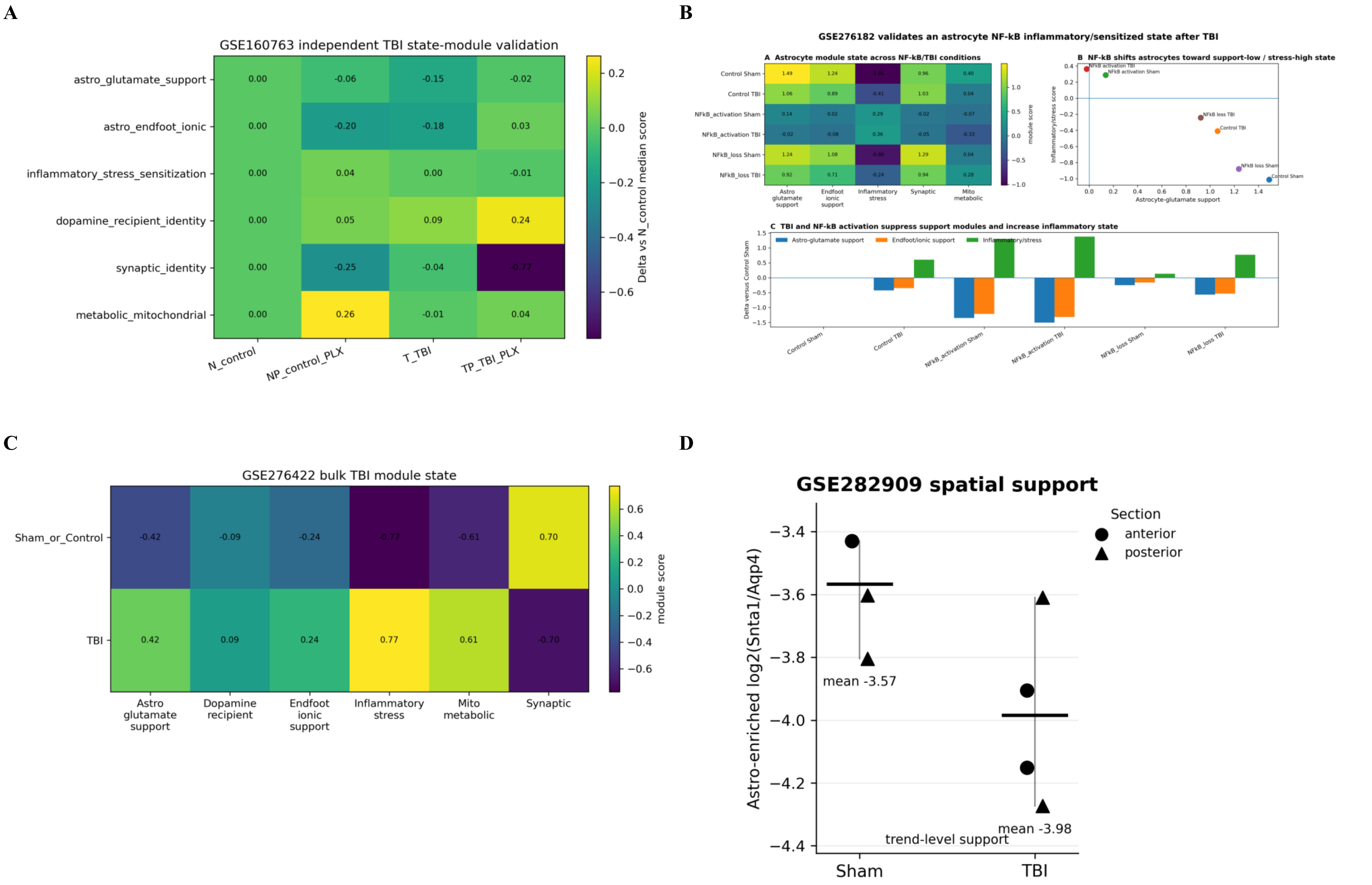

#### Legend

(A) GSE160763 independent mouse subacute-TBI context. Prespecified module definitions were projected onto the available control, PLX5622-control, TBI, and TBI+PLX5622 groups (two biological samples per group in the available reanalysis). Relative to the untreated control reference, the displayed median shifts were modest and context dependent: astrocyte glutamate support 0.00, -0.06, -0.15, -0.02; endfoot/ionic support 0.00, -0.20, -0.18, +0.03; inflammatory stress 0.00, +0.04, 0.00, -0.01; dopamine-recipient identity 0.00, +0.05, +0.09, +0.24; synaptic identity 0.00, -0.25, -0.04, -0.77; and metabolic/mitochondrial 0.00, +0.26, -0.01, +0.04. Given the very small groups and distinct experimental design, this panel is contextual rather than formal replication (Supplementary Table S63).

(B) GSE276182 astrocytic NF-κB perturbational context. NF-κB activation shifted the projected astrocyte programs toward lower glutamate-support and endfoot/ionic scores with higher inflammatory-stress scores. Relative to Control Sham, NF-κB activation produced approximately -1.35 and -1.51 shifts in astrocyte glutamate support in Sham and TBI conditions, -1.22 and -1.32 shifts in endfoot/ionic support, and +1.30 and +1.37 shifts in inflammatory stress. NF-κB-loss conditions were comparatively support-high/stress-low. This perturbational pattern supports partial separability of inflammatory-stress and astrocyte-support dimensions, but does not establish the temporal CCI configurations or a treatment-response phenotype (Supplementary Table S64).

(C) GSE276422 acute bulk-TBI context. Projected module scores in TBI relative to the control-centered display were positive for astrocyte glutamate support (+0.42), dopamine-recipient identity (+0.09), endfoot/ionic support (+0.24), inflammatory stress (+0.77), and metabolic/mitochondrial programs (+0.61), with reduced synaptic identity (-0.70). Because this dataset is bulk tissue, the module pattern is composition sensitive and is used only as acute contextual support (Supplementary Table S65).

(D) GSE282909 spatial endfoot-associated context. Astrocyte-enriched section summaries of log2(Snta1/Aqp4) were numerically lower in TBI than Sham mice (group means approximately -3.98 versus -3.57), which is directionally equivalent to greater Aqp4 relative to Snta1. With only two mice per group, this is trend-level contextual support and not inferential validation. Detailed spatial analyses and the stronger evidence boundaries are provided in Supplementary Figure 7 (Supplementary Table S66).

These panels provide cross-dataset contextual support for separable inflammatory, circuit-associated, and astrocyte-homeostatic transcriptional responses and do not establish categorical molecular states, treatment response, physical AQP4/SNTA1 uncoupling, or causal mechanisms.
