## Supplementary Methods for "Temporal transcriptomic remodeling after controlled cortical impact reveals delayed AQP4/SNTA1 expression imbalance associated with ion-homeostatic remodeling"

Reference numbering corresponds to the main manuscript.

### Dataset acquisition and analytical framework

This study was a secondary analysis of publicly available transcriptomic, proteomic, and spatial datasets. The primary discovery dataset was GSE269748, a murine traumatic brain injury atlas that includes multiple injury paradigms, brain regions, sexes, and post-injury intervals [10]. The GEO series contains 36 samples. For the prespecified controlled cortical impact (CCI) time-course analysis, the primary discovery set was restricted by sample metadata to the 21-sample male focal CCI/Naive temporal series: Naive (n = 7), CCI 24 h (n = 6), CCI 7 d (n = 4), and CCI 6 mo (n = 4). The remaining 15 samples represented the separate female cohort (n = 6), combined CCI plus hemorrhagic shock (n = 3), contralateral CCI tissue (n = 3), or repetitive closed-head injury (n = 3) and were excluded from the primary temporal analysis. This restriction was defined from GEO sample metadata rather than from expression-based sample exclusion. The biological sample, rather than the individual cell or nucleus, was the unit of inference throughout the primary analyses. Analyses were organized a priori around transcriptome-wide remodeling, neural circuit and neurotransmitter-associated programs, AQP4/SNTA1 expression imbalance, astrocyte endfoot and homeostatic programs, transcription-factor activity, and temporal integration. External datasets were used to test contextual convergence, robustness, and evidence boundaries rather than to require uniform replication across injury models, species, tissue compartments, or post-injury intervals.

Public datasets used for contextual analyses included the mouse TBI Drop-seq dataset GSE180862 [21], human chronic traumatic encephalopathy (CTE) single-nucleus dataset GSE155114 [22], human CTE detergent-insoluble proteomics dataset PXD007694 [23], mouse subacute TBI single-cell dataset GSE160763 [24], human repetitive-head-impact/early-CTE single-nucleus dataset GSE261807 [25], mouse CHIMERA spatial transcriptomic dataset GSE282909 [26], and acute human TBI dataset GSE209552 [9]. The astrocytic NF-kB perturbation datasets GSE276182 and GSE276422 were analyzed as perturbational and acute bulk-tissue context, respectively, and originated from Hein et al. [13]. The independent 1-year mouse TBI dataset PRJNA1280541 originated from Abou-El-Hassan et al. [12]. Human neuropathological boundary testing used middle temporal gyrus (MTG) data from the Seattle Alzheimer's Disease Brain Cell Atlas (SEA-AD) [27,28].

### GSE269748 sample selection, cell-level quality control, and pseudobulk construction

Raw GSE269748 10x-style count matrices were linked to the 21 prespecified biological samples using the locked sample-to-stage map. Cells were required to contain at least 500 total counts and at least 200 detected genes. After quality control, each cell received an astrocyte-support score calculated from the mean log1p-normalized expression of seven positive markers (Aldh1l1, Sox9, Slc1a3, Glul, Gja1, Gjb6, and Gfap) and an exclusion-lineage score calculated analogously from Snap25, Rbfox3, Slc17a7, Gad1, Gad2, Mbp, Plp1, Mog, Pdgfra, Cspg4, Pecam1, Cldn5, Kdr, Pdgfrb, Rgs5, Cx3cr1, P2ry12, Aif1, and Lyz2. The primary astrocyte-like gate retained cells with an astrocyte-support score at or above the sample-specific 90th percentile and greater than the exclusion-lineage score by more than 0.15. Importantly, neither Aqp4 nor Snta1 was included in the positive gating panel; thus, subsequent Aqp4 expression and AQP4/SNTA1 expression-imbalance analyses were not generated by direct selection on either transcript. A stricter sensitivity gate used the same marker definitions with a 95th-percentile astrocyte-support threshold and a greater-than-0.50 astrocyte-versus-exclusion score margin. Because this strategy is computationally marker-defined rather than fluorescence-purified or annotation-confirmed, the retained compartment is referred to throughout as astrocyte-like pseudobulk.

For each biological sample, raw counts from retained cells were summed gene-wise to form an animal-level pseudobulk profile. Library size was calculated from the summed pseudobulk counts, and counts per million (CPM) were calculated as the number of gene-level counts per one million total pseudobulk counts. The stricter astrocyte-like workflow was reconstructed directly from the archived source code and used as a reproducibility and sensitivity analysis. The reconstructed workflow reproduced the archived QC-passing and strict-gate cell counts and target-gene pseudobulk values, providing a computational check on gate implementation. Exploratory historical categorical state calls generated by an early pipeline were not used in the present manuscript.

For circuit and neurotransmitter analyses, an all-QC pseudobulk was constructed from the same 21 biological samples after the cell-level count and detected-gene thresholds above, without the astrocyte-marker gate. This broader compartment was used because neural circuit and transmitter-associated analyses were not intended to be astrocyte-specific. Counts from all QC-passing cells were summed within each biological sample, and both raw pseudobulk counts and CPM values were retained for downstream analyses.

### Transcriptome-wide filtering, normalization, differential expression, and unsupervised PCA

Transcriptome-wide differential expression in the primary astrocyte-like pseudobulk was performed with edgeR [29]. Raw sample-level pseudobulk counts were imported into a DGEList object. Genes were filtered using filterByExpr with injury stage supplied as the grouping factor, reducing the common testing universe from 33,451 genes to 16,209 genes. Library-composition normalization factors were calculated using the trimmed mean of M-values (TMM) method [30]. A no-intercept design matrix was fit for Naive, CCI 24 h, CCI 7 d, and CCI 6 mo. Dispersion was estimated with robust estimation, followed by robust quasi-likelihood model fitting and quasi-likelihood F tests [31]. Prespecified contrasts compared each CCI stage with Naive. P values were adjusted by the Benjamini-Hochberg procedure [32]. The primary differential-expression display defined large-effect differentially expressed genes as FDR $< 0.05$ with $|\log2$ fold change$| >= 1$; threshold sensitivity was evaluated using FDR alone and FDR with $|\log2$ fold change$| >= 0.5$.

A chronic-stage gate-sensitivity analysis compared the primary astrocyte-like pseudobulk with the historical stricter astrocyte-like gate in the matched Naive ($n = 7$) versus CCI 6 mo ($n = 4$) cohort. Both count matrices were analyzed with the same edgeR filtering, TMM normalization, and quasi-likelihood framework. Concordance of gene-level log2 fold-change estimates across the common tested genes was assessed by Spearman correlation, and directional concordance was summarized among genes meeting the prespecified significance/effect-size criteria in both analyses. This robustness analysis was restricted to the Naive versus 6-mo comparison and was not interpreted as validation of the full time course.

Whole-transcriptome principal component analysis (PCA) was used to assess unsupervised sample structure. The locked sample-level PCA coordinates were generated from the normalized whole-transcriptome astrocyte-like expression matrix without using injury-stage labels to calculate the principal components. PC1 and PC2 were used for visualization and downstream post hoc program-axis association analyses. Sample-similarity robustness was additionally assessed using a Pearson correlation matrix of the sample-level expression profiles followed by hierarchical clustering. PCA and correlation structure were treated as descriptive representations of transcriptomic organization rather than evidence for discrete categorical disease states.

### Prespecified molecular-program scoring

Prespecified transcriptional programs were calculated from biologically defined gene sets. The astrocyte glutamate-support program comprised Slc1a2, Slc1a3, Glul, Grm3, Gja1, Gjb6, Aldh1l1, and Sox9. The endfoot/ionic program comprised Aqp4, Snta1, Kcnj10, Dmd, Dtna, and Dag1. The inflammatory-stress program comprised Fos, Jun, Junb, Jund, Atf3, Egr1, Dusp1, Arc, Nr4a1, Gfap, Vim, C3, Serpina3n, Lcn2, Nfkbia, Rela, Tnf, Il1b, Ccl2, and Cxcl10. The dopamine-recipient program comprised Drd1, Drd2, Ppp1r1b, Adora2a, Rgs9, Penk, and Tac1. The synaptic-identity program comprised Syn1, Syp, Snap25, Gria1, Gria2, Gria3, Gria4, Grin1, Grin2a, and Grin2b. A metabolic/mitochondrial program was constructed from detected genes beginning with Nduf, Cox, or Atp5 in the archived workflow.

For the initial module landscape, expression was transformed as log2(CPM + 1), and each sample-level program score was calculated as the mean transformed expression of represented genes. Astrocyte glutamate-support, endfoot/ionic, and inflammatory-stress scores were calculated from the astrocyte-like compartment, whereas dopamine-recipient, synaptic-identity, and metabolic/mitochondrial scores were calculated from the all-QC pseudobulk. Sample-level values were centered to the corresponding Naive-group reference for temporal trajectory plots. For the standardized heatmap, values were standardized within program before stage-level means were calculated. These scores quantify coordinated transcriptional representation of prespecified gene sets and were not interpreted as direct measurements of glutamate clearance, synaptic transmission, receptor activation, metabolic flux, water transport, or potassium handling.

### Circuit and neurotransmitter-associated enrichment analyses

Circuit-associated analyses used the all-QC GSE269748 pseudobulk from the same 21 biological samples. Differential-expression statistics were generated with edgeR using filterByExpr, TMM normalization, a no-intercept stage design, robust dispersion estimation, and quasi-likelihood testing as described above [29-31]. For each stage-versus-Naive contrast, genes were ranked by multiplying the sign of the log2 fold change by the square root of the quasi-likelihood F statistic. Preranked gene set enrichment analysis was performed with fgseaMultilevel [33]. Gene sets were obtained with msigdbr from the mouse MSigDB C5 Gene Ontology Biological Process and C2 Reactome collections [34,35]. The transcriptome-wide screen used minimum and maximum pathway sizes of 10 and 500 genes, respectively, with eps = 0 and 10,000 simple permutations for multilevel estimation. Neural, synaptic, and neurotransmission-associated pathways displayed in the focused analysis were extracted from the completed transcriptome-wide results rather than used to prefilter the ranked transcriptome.

Seven transmitter-system gene collections were then assembled from MSigDB pathway families matching dopaminergic, glutamatergic, GABAergic, serotonergic, adrenergic, cholinergic, or purinergic/adenosine terms. Enrichment was recalculated with fgseaMultilevel using minimum and maximum sizes of 10 and 2,000 genes, respectively. To determine whether apparent multi-transmitter remodeling could be explained by shared genes, non-overlapping transmitter modules were constructed. Genes assigned to more than one transmitter system were removed, and remaining genes were required to be detected in at least four biological samples and to have mean CPM >= 0.1. Final non-overlapping modules contained 138 glutamatergic, 69 GABAergic, 46 purinergic/adenosine, 32 dopaminergic, 27 cholinergic, 23 adrenergic, and 14 serotonergic genes. Expression was transformed as log2(CPM + 1), standardized gene-wise across the 21 samples, averaged within each module, and centered to the corresponding Naive-group mean.

Targeted follow-up analysis used the locked 10-gene synaptic-identity set and 7-gene dopamine-recipient set defined above. For each gene, log2(CPM + 1) expression was standardized across the 21 samples. Sample-level module scores were the mean of constituent gene z scores and were re-centered to the module-specific Naive mean. Gene-level heatmaps used stage mean gene z scores relative to the corresponding Naive gene mean. To test single-gene dependence of the dopamine-recipient result, the module was recalculated after sequential omission of each constituent gene; each leave-one-gene-out version was re-centered to its own Naive reference before the chronic-stage direction was evaluated.

To assess whether the targeted circuit-module results could be explained by sample-level cellular composition, the raw GSE269748 matrices from the same 21 animals were independently reprocessed using the locked cell-level quality-control thresholds. All 278,108 QC-passing cells were included. The 17 genes constituting the synaptic-identity and dopamine-recipient outcomes were explicitly excluded from feature selection and clustering to prevent outcome-dependent composition estimation. Batch-aware highly variable gene selection followed by PCA, nearest-neighbor graph construction, and Leiden clustering generated 28 unsupervised clusters. For each biological sample, cluster counts were converted to proportions; after addition of a 0.5-cell pseudocount, proportions were centered-log-ratio transformed and summarized by PCA. Composition PC1 explained 56.7% of sample-level composition variance and PC1 plus PC2 explained 81.5%. The primary sensitivity model regressed the original, unchanged sample-level module score on injury stage and composition PC1 using HC3 heteroskedasticity-robust covariance with small-sample t/F inference. A stage-plus-PC1-plus-PC2 model was evaluated secondarily; because this model produced substantial multicollinearity (maximum variance-inflation factor approximately 53), it was used as an omnibus robustness analysis rather than for strong interpretation of individual stage coefficients. Before examining Figure 2 outcome genes, independent canonical-marker neuronal gates were also audited, but the preferred core-neuronal gate retained as few as 10 cells in one animal; neuron-only animal-level pseudobulk was therefore not considered sufficiently stable for inference and no neuron-specific claim was made.

### AQP4/SNTA1 expression-imbalance analysis

Aqp4 and Snta1 were evaluated as expression-level variables in the marker-defined astrocyte-like pseudobulk. Individual gene trajectories were summarized on the log2(CPM + 1) scale. The primary AQP4/SNTA1 expression-imbalance score was calculated from sample-level CPM as log2(Aqp4 + 0.5) minus log2(Snta1 + 0.5), such that higher scores indicated greater Aqp4 expression relative to Snta1. For stage-centered displays and tests, the Naive-group mean of the imbalance score was subtracted from each sample. Alternative formulations included the direct Aqp4-Snta1 difference on the log2(CPM + 1) scale and the difference between standardized Aqp4 and Snta1 expression. These sensitivity analyses were used to determine whether temporal separation depended on a single scaling choice. The expression-imbalance metric was not interpreted as molecular stoichiometry, physical AQP4-SNTA1 uncoupling, AQP4 depolarization, or perivascular protein mislocalization.

Stage-versus-Naive comparisons for the targeted Aqp4, Snta1, and imbalance analyses used exact two-sided label-permutation tests at the biological-sample level. With the observed group sizes, all possible allocations were enumerated where computationally feasible. Bootstrap confidence intervals for stage mean differences were generated from 20,000 resamples. Multiple comparisons within the planned analysis family were adjusted by the Benjamini-Hochberg procedure [32].

To evaluate departure from the uninjured expression relationship, an ordinary least-squares reference model was fit in Naive samples only with Snta1 log2(CPM + 1) as the dependent variable and Aqp4 log2(CPM + 1) as the predictor. This Naive model was not refit in post-injury samples. For every sample, the signed residual was calculated as observed Snta1 minus the Snta1 value predicted by the Naive reference line; the absolute residual quantified the magnitude of departure from the Naive Aqp4-Snta1 expression relationship. Residual analyses were interpreted as expression-level deviation only and not as evidence of physical protein coupling or uncoupling.

### Endfoot/homeostatic context and inflammatory-stress adjustment

The broader endfoot/homeostatic context was assessed using Aqp4, Snta1, Dag1, Dmd, Dtna, Kcnj10, Gja1, Gjb6, Slc1a2, Slc1a3, Atp1a2, and Glul. Gene expression was transformed as log2(CPM + 0.5), and stage-level changes were calculated relative to each gene-specific Naive mean. The previously validated endfoot/ionic, astrocyte glutamate-support, inflammatory-stress, and metabolic/mitochondrial module scores were reused without redefining gene sets. For comparative temporal visualization, each module was centered to its Naive-group mean and scaled by the overall sample-level standard deviation.

The sample-level association between AQP4/SNTA1 expression imbalance and inflammatory-stress activity was evaluated using Spearman correlation. To determine whether stage-related imbalance remained after accounting for measured inflammatory-stress variation, an ordinary least-squares model included AQP4/SNTA1 imbalance as the outcome, injury stage as a categorical predictor with Naive as the reference, and the inflammatory-stress score as a covariate. Heteroskedasticity-consistent HC3 covariance estimates were used for coefficient uncertainty. Model-adjusted stage values were calculated at the mean inflammatory-stress score. Gene-level chronic effects used Welch tests with Benjamini-Hochberg adjustment and bootstrap confidence intervals for descriptive effect estimates.

### Prespecified pathway screen and stage-adjusted specificity analyses

A prespecified pathway screen evaluated 14 biological programs in the 21 GSE269748 biological samples: endfoot anchoring/dystrophin-associated protein complex (DAPC), extracellular matrix/basement membrane, potassium/ion homeostasis, astrocyte glutamate support, reactive gliosis, complement/innate immunity, NF-kB signaling, JAK-STAT/interferon signaling, hypoxia/angiogenic response, NRF2/oxidative-stress response, RXR-PPAR lipid handling, mitochondrial oxidative phosphorylation, autophagy/lysosome, and endoplasmic-reticulum stress/unfolded-protein response. AQP4 and SNTA1 were excluded from the expression matrix used to construct pathway scores to avoid direct mathematical circularity with the imbalance outcome. Duplicate gene symbols were collapsed by averaging, genes without variation were removed, and each retained gene was z standardized across samples. Pathway scores were the mean of represented constituent-gene z scores. Pathways were required to contain at least four represented genes and at least 35% of their prespecified gene set.

Both the AQP4/SNTA1 imbalance and each pathway score were standardized before regression. For each pathway, the primary ordinary least-squares model included the pathway score as the outcome, standardized AQP4/SNTA1 imbalance as the predictor of interest, and injury stage as a categorical covariate with Naive as the reference. Coefficients and 95% confidence intervals were estimated with HC3 robust covariance. Statistical support was evaluated by 10,000 stage-stratified permutations of the imbalance values, preserving the stage composition of the data. Two-sided empirical permutation P values were calculated using a plus-one correction and adjusted across the pathway family using the Benjamini-Hochberg procedure [32]. Leave-one-sample-out models were used to assess coefficient sign stability. Because stage and imbalance were correlated, the proportion of imbalance variance attributable to stage and variance-inflation diagnostics were examined before interpreting stage-adjusted coefficients.

For pathways that remained supported in the prespecified screen, shared transcriptional structure was evaluated after residualizing each pathway score for injury stage. Pairwise Spearman correlations were calculated among stage residuals. For the four supported core programs, residual values were standardized and subjected to PCA by singular value decomposition; component orientation was set so that the mean loading was positive. This residual PCA was used to quantify redundancy among the core transcriptional programs, not to define a biological latent state.

Specificity of the potassium/ion-homeostasis association was evaluated with sequential covariate adjustment. The potassium score was modeled as a function of AQP4/SNTA1 imbalance and injury stage, followed by models additionally including reactive-gliosis activity, a broad general-response PC1, or both. HC3 covariance estimates were used throughout. Significance after nuisance adjustment was evaluated with a stage-stratified Freedman-Lane permutation procedure: the reduced model containing stage and nuisance covariates was fit, residuals were permuted within stage, the outcome was reconstructed, and the imbalance coefficient was refit in the full model. Ten thousand permutations were used for each model, followed by Benjamini-Hochberg correction across the prespecified comparison family. Covariate-adjusted specificity was interpreted as statistical specificity and not as causal independence.

### Transcription-factor activity inference and reciprocal AP-1 analysis

Transcription-factor (TF) activity was inferred from the GSE269748 expression matrix using the Python implementation of decoupler v2.2.0 [36]. Mouse CollecTRI regulons were obtained through the OmniPath interface [37,38]. AQP4 and SNTA1 were removed before TF inference. Infinite values were treated as missing; genes with more than 20% missing observations were excluded; remaining missing observations were imputed with the within-gene median; and invariant genes were removed. TF activities were inferred using the decoupler univariate linear model implementation (dc.mt.ulm), requiring at least five mapped targets per TF.

Each inferred TF activity was tested for association with standardized AQP4/SNTA1 imbalance in a stage-adjusted HC3 ordinary least-squares model. Statistical support was evaluated with 10,000 stage-stratified permutations and Benjamini-Hochberg correction, and leave-one-sample-out refitting was used to assess directional stability. Candidate TFs were interpreted as inferred regulatory activity rather than measured TF binding or direct transcriptional regulation. AP-1 family activity was subsequently evaluated in reciprocal specificity models with the potassium/ion-homeostasis program. In one direction, potassium/ion homeostasis was modeled with imbalance, stage, and AP-1 activity; in the reciprocal direction, AP-1 activity was modeled with imbalance, stage, and the potassium score. Additional versions included reactive-gliosis or broad-response-PC1 covariates. HC3 uncertainty and stage-stratified permutation inference were used for all reciprocal models. These analyses were not interpreted as mediation or causal ordering.

### Potassium/ion-homeostasis gene-level and resampling robustness

The locked potassium/ion-homeostasis score comprised Kcnj10, Kcnj16, Atp1a2, Atp1b2, Slc12a2, Slc8a1, Clcn2, Trpv4, Gja1, and Gjb6. Each gene was standardized across the 21 biological samples, and the program score was the mean of available gene z scores. Gene-level associations with AQP4/SNTA1 imbalance were evaluated in stage-adjusted HC3 models with the same stage-stratified permutation framework. Because the score was intended as a distributed transcriptional program, constituent genes were not required to change uniformly in direction.

Single-gene dependence was tested by recalculating the program after sequential omission of each of the 10 genes. Grouped omission analyses additionally removed Kcnj10 alone, both connexins (Gja1/Gjb6), both Na+/K+-ATPase components (Atp1a2/Atp1b2), or both inward-rectifier potassium-channel genes (Kcnj10/Kcnj16). Every modified score was re-standardized and refit using the same stage-adjusted HC3 model. The incremental contribution of the imbalance term was summarized using partial R-squared from the full model relative to a stage-only reduced model. Significance was evaluated by 10,000 within-stage Freedman-Lane permutations.

Sample-level stability was assessed by leave-one-biological-sample-out refitting across all 21 samples and by a 5,000-resample stage-stratified bootstrap that resampled biological samples with replacement within stage. The historical strict astrocyte-like gate was reconstructed as described above to verify pseudobulk reproducibility. Cross-injury-model sensitivity was evaluated in the matched 24-h strict-gate subset containing Naive ($n = 7$), repetitive closed-head injury (rCHI; $n = 3$), CCI ($n = 6$), and CCI plus hemorrhagic shock (CCI+HS; $n = 3$). The potassium-imbalance relationship was evaluated after adjustment for injury model, followed by interaction and leave-one-injury-model-out sensitivity analyses. Given the small model-specific groups, these analyses were treated as within-dataset robustness rather than independent replication or evidence of model equivalence.

### Temporal-profile analysis and six-program molecular synthesis

Four transcriptional dimensions developed through the preceding analyses were integrated at the sample level: synaptic identity, dopamine-recipient identity, AQP4/SNTA1 expression imbalance, and potassium/ion homeostasis. Each program was centered to its Naive-group mean and scaled by the overall standard deviation across the 21 samples, fixing the Naive reference to zero. Stage mean differences were summarized with 20,000 bootstrap resamples. Stage-versus-Naive comparisons used exact two-group permutation tests based on the biological samples; a prespecified delayed/chronic-versus-24-h contrast combined the 7-d and 6-mo samples. P values from the planned contrast family were controlled by global Benjamini-Hochberg adjustment, with within-program adjusted values retained as sensitivity summaries.

To test whether the four programs followed a common temporal profile, stage labels were permuted at the biological-sample level. For each program, a one-way stage statistic was evaluated against 200,000 stage-label permutations. Pairwise profile-difference tests were performed on the sample-level difference between each pair of standardized programs and evaluated with a one-way stage statistic; 200,000 stage-label permutations were used for each pair and the six pairwise tests were Benjamini-Hochberg adjusted. An omnibus multivariate profile-difference statistic was calculated from program-difference coordinates and evaluated with 200,000 stage-label permutations. Because the data are cross-sectional, connected stage summaries were interpreted as group-level temporal patterns and not longitudinal trajectories measured repeatedly in the same animals.

For descriptive synthesis, six programs were placed on a common Naive-centered standardized scale: inflammatory stress, astrocyte glutamate support, synaptic identity, dopamine-recipient identity, AQP4/SNTA1 imbalance, and potassium/ion homeostasis. The four locked temporal-analysis variables were reused directly. Inflammatory-stress and astrocyte glutamate-support scores were centered to their respective Naive-group means and scaled by the overall sample-level standard deviation. Stage means were then calculated for the 24-h, 7-d, and 6-mo groups. This six-program panel was a descriptive synthesis and did not introduce a new composite index or additional inferential model.

### Post hoc recovery of predefined programs on unsupervised PCA axes

To determine whether the predefined molecular programs were represented in the dominant unsupervised transcriptome-wide structure, the locked whole-transcriptome PC1 and PC2 coordinates were merged with the six sample-level program scores. Spearman correlations were calculated between each program and PC1 or PC2. PCA coordinates had been generated without using the program scores or injury-stage labels; biological interpretation of the axes was therefore assigned only after PCA through these post hoc correlations. The analysis was treated as axis recovery and correspondence, not as independent proof of discrete molecular states.

### External mouse, perturbational, human, proteomic, and spatial contextual analyses

External datasets were analyzed with a common principle: the original biological unit of replication was preserved, the orientation of AQP4/SNTA1 metrics was harmonized so that higher values represented greater AQP4 relative to SNTA1, and conclusions were restricted to contextual convergence or non-replication unless the external design directly matched the discovery analysis. GSE180862 was analyzed as an independent mouse Drop-seq dataset [21]. Astrocytes were pseudobulked by mouse and brain region, and the same AQP4/SNTA1 expression-imbalance definition used in the discovery analysis was applied to sample-level CPM values. The prespecified 7-d TBI-versus-Sham comparison pooled cortex and hippocampus while adjusting for region in an ordinary least-squares model. Exact mouse-level permutation assigned three TBI labels among six mice while preserving paired regional observations. The 24-h contrast and region-specific contrasts were secondary contextual analyses.

GSE155114 was analyzed at the donor level as chronic human transcriptomic context [22]. Marker-inferred astrocyte summaries from eight neuropathologically confirmed CTE donors and eight controls were reoriented to a common log2(AQP4/SNTA1) direction. PXD007694 provided a cross-modal human CTE proteomic context [23]. Sample-level AQP4/SNTA1 protein contrasts were calculated for six control and 11 CTE samples from detergent-insoluble proteomics; the group comparison used a two-sided Mann-Whitney test. These datasets were not treated as mechanistic validation of mouse AQP4 localization or SNTA1-dependent anchoring.

GSE160763 provided independent mouse subacute TBI context [24]. The prespecified program definitions were projected onto sample-level expression summaries from control, PLX5622-treated control, TBI, and TBI plus PLX5622 groups, with two biological samples per group in the available reanalysis. The dataset was used to assess whether the discovery configuration was a generic feature of 7-d injury rather than as a required replication cohort. GSE261807 provided human repetitive-head-impact and low-stage CTE context [25]. Because marker-inferred astrocyte summaries varied with the retained-cell threshold, analyses were repeated across the prespecified 0.5%, 1%, and 2% astrocyte-selection thresholds and interpreted as threshold sensitivity rather than robust validation.

The astrocytic NF-kB perturbation dataset GSE276182 and acute bulk-TBI dataset GSE276422 were obtained from the study by Hein et al. [13]. The locked transcriptional programs were projected onto the processed expression matrices to provide perturbational context for inflammatory-stress versus astrocyte-support relationships and acute bulk-tissue context, respectively. Bulk-tissue results were explicitly treated as composition-sensitive. PRJNA1280541, comprising sorted astrocytes at 365 days after TBI, was analyzed as an independent chronic mouse boundary dataset [12]. The locked AQP4/SNTA1 and potassium/ion-homeostasis definitions were applied without redefining the discovery gene sets, with exact small-sample comparisons and leave-one-sample/gene sensitivity used to characterize directional support.

### Acute human TBI module projection

Acute human TBI context was evaluated in GSE209552 [9]. The final analysis included 12 TBI and 5 control samples. The six locked transcriptional programs were calculated in the corresponding analysis compartment and centered to the control-group median. For each module, the primary descriptive effect was the TBI median shift relative to the control median. Uncertainty intervals were generated from 10,000 bootstrap resamples of the control and TBI sample scores. Because the available sample structure did not support independent donor-level replication of the chronic mouse state, these intervals were treated as descriptive sample-level uncertainty and the analysis was interpreted as acute human context rather than validation of chronic CCI transcriptional configurations.

### SEA-AD donor-level human neuropathology boundary analysis

SEA-AD MTG single-nucleus transcriptomic and quantitative neuropathology resources were used to test whether the mouse-derived AQP4/SNTA1 transcriptional phenotype generalized to human tau burden [27,28]. Eighty-four donors with matched MTG transcriptomic and quantitative neuropathology data were retained. For each donor, nuclei annotated as Subclass = Astrocyte and Used in analysis = True were selected from the final-nuclei objects. Raw UMI counts were extracted from the UMIs layer and summed across retained astrocyte nuclei to generate donor-level raw-count pseudobulk. Donor-level CPM was calculated from the astrocyte pseudobulk library size. AQP4 and SNTA1 expression values were transformed using the same pseudocount-based log2 scale as the discovery imbalance analysis, and the donor-level AQP4/SNTA1 expression-imbalance score was calculated as transformed AQP4 minus transformed SNTA1. The imbalance score was standardized to one sample-level standard deviation for regression.

The prespecified primary neuropathology outcome was log1p-transformed MTG percent AT8-positive area. The primary ordinary least-squares model used HC3 robust covariance and included standardized AQP4/SNTA1 imbalance, standardized age at death, sex as a categorical covariate, and standardized log1p-transformed MTG 6E10 amyloid-positive area. The incremental variance associated with the imbalance term was summarized using partial R-squared. A 10,000-permutation conditional test was implemented by residualizing the exposure with respect to the covariates, permuting the exposure residuals across donors, reconstructing the exposure, and refitting the full model. Leave-one-donor-out HC3 refitting was used to assess influence.

Prespecified sensitivity models additionally adjusted for log1p GFAP burden, postmortem interval and RNA integrity number, Iba1 burden, or log astrocyte count; restricted analyses to donors with at least 100 or at least 200 retained astrocyte nuclei; and evaluated secondary tau outcomes including AT8-positive cell density and continuous pTau-related measures with corresponding amyloid adjustment. This analysis was designed as an evidence-boundary test. A null association was therefore retained and reported rather than being treated as a failed validation or excluded from the study.

### GSE282909 spatial transcriptomic analysis

Spatial organization was evaluated in GSE282909, a mouse CHIMERA mild-TBI spatial transcriptomic dataset [26]. Eight Visium sections were analyzed, comprising anterior and posterior sections from two Sham and two TBI mice. Spatial assay counts were extracted from archived Seurat-compatible RDS objects [39]. Within each section, spot library sizes were used to calculate CPM and required genes were transformed as log2(CPM + 0.5). Spatial scores were calculated for AQP4/SNTA1 imbalance, the locked 10-gene potassium/ion-homeostasis program, a 9-gene astrocyte-support program (Aldh1l1, Sox9, Slc1a3, Slc1a2, Glul, Gja1, Gjb6, Aqp4, Aldoc), an endfoot program (Aqp4, Snta1, Dmd, Dtna, Dag1, Agrn, Kcnj10), and a vascular-interface program (Pecam1, Cldn5, Kdr, Emcn, Vwf, Pdgfrb, Rgs5, Cspg4, Acta2, Tagln). Within-section z scores were used for visualization only. Astrocyte-enriched spots were defined as spots in the upper quartile of the section-specific astrocyte-support score.

Spatial autocorrelation was quantified with global Moran's I [40]. A symmetric binary k-nearest-neighbor graph was constructed from spot image coordinates with $k = 6$ as the primary neighborhood definition. Sensitivity analyses used $k =$ 4, 6, and 8. For each section and score, a one-sided permutation test for positive spatial autocorrelation used 999 random permutations, and the primary spatial-test family was adjusted using the Benjamini-Hochberg procedure [32]. Section-level spatial tests were interpreted as evidence of within-section anatomical organization. For condition-level summaries, the mouse was the biological replicate ($n = 2$ per condition); sections nested within a mouse were not treated as independent biological replicates. Spot-level correspondence between AQP4/SNTA1 imbalance and potassium/ion-homeostasis scores was treated as descriptive spatial context and not as evidence of physical colocalization or endfoot dysfunction.

### Statistical analysis and software

All statistical tests were two-sided unless a directional spatial-autocorrelation test or other prespecified directional sensitivity analysis is explicitly identified. Biological samples, mice, or human donors were used as the unit of inference according to the original experimental design; cells, nuclei, and spatial spots were not treated as independent biological replicates for group-level inference. Exact permutation tests were preferred for small group comparisons when all label allocations could be enumerated. Larger permutation analyses used fixed random seeds recorded in the analysis scripts. Bootstrap procedures resampled the appropriate biological unit with replacement, and stage-stratified procedures preserved the observed number of samples in each injury stage. Multiple-testing correction used the Benjamini-Hochberg false-discovery-rate procedure [32]. Spearman correlation was used for monotonic sample-level associations and for post hoc program-to-PC correspondence. Ordinary least-squares models used HC3 heteroskedasticity-consistent covariance estimates when coefficient inference could be sensitive to unequal residual variance. No causal, mediation, or longitudinal inference was assigned to cross-sectional regression or temporal-profile analyses.

The principal computational environments recovered from the archived final analyses were Python 3.12.3 and R 4.3.3. The Figure 5 regulatory/specificity environment used decoupler 2.2.0, pandas 3.0.3, NumPy 2.4.6, SciPy 1.18.0, statsmodels 0.14.6, anndata 0.13.2, and scanpy 1.12.2. The R differential-expression and enrichment environment used edgeR 4.0.16, limma 3.58.1, fgsea 1.28.0, msigdbr 26.1.0, ggplot2 4.0.3, dplyr 1.2.1, tidyr 1.3.2, and data.table 1.18.4. General Python visualization and tabular processing additionally used matplotlib 3.10.9 and seaborn 0.13.2. Archived spatial scripts were executed in R 4.3.1 and used Seurat-compatible objects together with Matrix 1.6.1.1 and patchwork 1.3.2; an independently verifiable Seurat package version was not retained in the archived environment and is therefore not inferred here. SRA Toolkit fasterq-dump 3.4.1 was used where raw SRA retrieval was required. Software versions are reported from the audited analysis environments rather than reconstructed from current package releases.

### Data and code availability

All primary and contextual datasets analyzed in this study are publicly available through the accessions listed above: GSE269748, GSE180862, GSE155114, GSE160763, GSE209552, GSE261807, GSE276182, GSE276422, GSE282909, PXD007694, and PRJNA1280541, together with the publicly released SEA-AD MTG transcriptomic and quantitative neuropathology resources. Figure-level source tables, supplementary tables, locked gene-set definitions, and the custom scripts used for the analyses are maintained in the study reproducibility package. A permanent public repository identifier for the final code and source-data archive should be inserted here before submission: [repository/DOI to be added].

All analyses used publicly available datasets. The complete reference list is provided in the main manuscript.
